# Marine phytoplankton and heterotrophic bacteria rapidly adapt to future pCO_2_ conditions in experimental co-cultures

**DOI:** 10.1101/2024.02.07.579367

**Authors:** Zhiying Lu, Elizabeth Entwistle, Matthew D. Kuhl, Alexander R. Durrant, Marcelo Malisano Barreto Filho, Anuradha Goswami, J. Jeffrey Morris

**Affiliations:** Department of Biology, University of Alabama at Birmingham

## Abstract

The CO_2_ content of Earth’s atmosphere is rapidly increasing due to human consumption of fossil fuels. Models based on short-term culture experiments predict that major changes will occur in marine phytoplankton communities in the future ocean, but these models rarely consider how the evolutionary potential of phytoplankton or interactions within marine microbial communities may influence these changes. Here we experimentally evolved representatives of four phytoplankton functional types (silicifiers, calcifiers, coastal cyanobacteria, and oligotrophic cyanobacteria) in co-culture with a heterotrophic bacterium, *Alteromonas*, under either present-day or predicted future pCO_2_ conditions. Growth rates of cyanobacteria generally increased under both conditions, and the growth defects observed in ancestral *Prochlorococcus* cultures at elevated pCO_2_ and in axenic culture were diminished after evolution, possibly due to regulatory mutations in antioxidant genes. Except for *Prochlorococcus*, mutational profiles suggested phytoplankton experienced primarily purifying selection, but most *Alteromonas* lineages showed evidence of directional selection, especially when co-cultured with eukaryotic phytoplankton, where evolution appeared to favor a broad metabolic switch from growth on small organic acids to catabolism of more complex carbon substrates. Evolved *Alteromonas* were also poorer “helpers” for *Prochlorococcus*, supporting the assertion that the interaction between *Prochlorococcus* and heterotrophic bacteria is not a true mutualism but rather a competitive interaction stabilized by Black Queen processes. This work provides new insights on how phytoplankton will respond to anthropogenic change and on the evolutionary mechanisms governing the structure and function of marine microbial communities.

---

As a result of human fossil fuel use, Earth’s atmospheric pCO_2_ has increased by ∼40% since the industrial revolution and is projected to further double by the end of the century (*1*). Elevated pCO_2_ affects ocean biogeochemistry primarily by raising ocean temperature and lowering ocean pH, and marine ecosystems may experience extensive changes as organisms better adapted to the new conditions locally replace those that are currently dominant (*2, 3*). For example, based on short-term culture experiments, many smaller phytoplankton such as cyanobacteria have been predicted to become more abundant in warmer and/or more acidic oceans (*4, 5*), whereas *Prochlorococcus*, which is numerically dominant in current oligotrophic oceans, has shown pronounced reductions in growth rate at elevated pCO_2_ (*6–8*). However, making predictions about future seas from culture experiments is complicated by their simplicity, and many experiments have shown that phytoplankton respond differently to predicted future conditions depending on the community context in which they are observed. *Prochlorococcus*, for instance, displays remarkably different responses to increased pCO_2_ depending on whether it is in axenic culture, in co-culture with heterotrophic (*6, 8*) or other photosynthetic microbes (*7*), or in intact *in situ* communities (*9*). Heterotrophic bacteria in general have profound impacts on the metabolisms of phytoplankton in culture experiments (*10–15*), and very little is known about how anthropogenic change will impact pelagic bacterial assemblages, or how those changes may reverberate through the phytoplankton. Moreover, natural selection will act on phytoplankton populations to adapt them to changing environments (*16*), and especially for microbial populations, may be able to mitigate negative impacts of environmental change (*17*). On the other hand, adaptation to ocean acidification may also alter microbial physiologies or community interactions in ways that are difficult to predict, but that may nevertheless have far-reaching impacts on ecosystem structure and function.

With few exceptions (*18–22*), studies of phytoplankton responses to future pCO_2_ have not considered long-term evolutionary responses, and to our knowledge no study has observed co-evolution between phytoplankton and heterotrophic bacteria under simulated global change conditions. We therefore sought to explore how the growth rates, metabolisms, and genomes of simple assemblages of marine phytoplankton and heterotrophic bacteria adapted to ocean acidification using long-term experimental evolution. We grew a variety of unicellular phytoplankton in low-density semicontinuous cultures either under current atmospheric pCO_2_ conditions or projected year 2100 conditions (400 vs. 800 ppm, respectively) for approximately 500 generations. We selected taxa representing major ecological functional groups: a brackish-water cyanobacterium *Synechocystis* PCC6803, the open ocean diatom *Thalassiosira oceanica* CCMP1005, the important bloom-forming coccolithophore *Emiliania huxleyi* CCMP371, the coastal picocyanobacterium *Synechococcus* CC9311, and the highly abundant oligotrophic picocyanobacterium *Prochlorococcus* MIT9312. These groups are ecologically important not just because they interact with ocean biogeochemistry in distinct ways (e.g.,, *Prochlorococcus* as an oligotrophic specialist, *T. oceanica* and *E. huxleyi* as silicifiers and calcifiers, respectively), but also because they have conspicuously different growth rate responses to elevated pCO_2_ in short-term culture experiments (i.e., positive for *Synechococcus,* negative for *Prochlorococcus*, and no net effect for diatoms or coccolithophores) (*23*).

In addition to these photoautotrophic taxa, each culture (apart from *Synechocystis*, which was evolved axenically in a pilot experiment) also included a single heterotrophic bacterial strain, the Gammaproteobacterium *Alteromonas macleodii* EZ55. Strains of *Alteromonas* are commonly found inhabiting phytoplankton cultures (*24, 25*) and are ubiquitous in ocean waters worldwide (*26*). The decision to use mixed instead of axenic cultures was made for two reasons. First, the bacteria facilitated carbon cycling of photosynthetic exudates, preventing the environmentally unrealistic overaccumulation of metabolites in the cultures that might obscure any evolutionary responses of the phytoplankton to pCO_2_ manipulation (*27, 28*). Second, one of our phytoplankton subjects, *Prochlorococcus*, grew very poorly in axenic culture especially under elevated pCO_2_ conditions (*6, 8, 29*), and in preliminary experiments we were unable to sustain axenic *Prochlorococcus* in semicontinuous low-density cultures for more than a few transfers.

We tracked phytoplankton growth in six replicate cultures of each phytoplankton at each pCO_2_ condition, achieving at least 500 generations for most cultures (Table S1). While there was substantial variability transfer-to-transfer in Malthusian growth rates (MGR), regression lines fit to the overall data for each of the 48 lineages revealed clear fitness trends (Fig. 1A–E), with all cyanobacterial lineages and one eukaryotic phytoplankton treatment (*E. huxleyi*, 400 ppm pCO_2_) exhibiting significant increases in growth rate over the course of the experiment (Fig. 1F). The pace of evolution was only significantly different between pCO_2_ conditions for two species however: *Prochlorococcus’* growth rate evolved faster at 800 ppm pCO_2_, and *E. huxleyi*’s evolved faster at 400 ppm (Fig. 1F).

**Figure 1.**
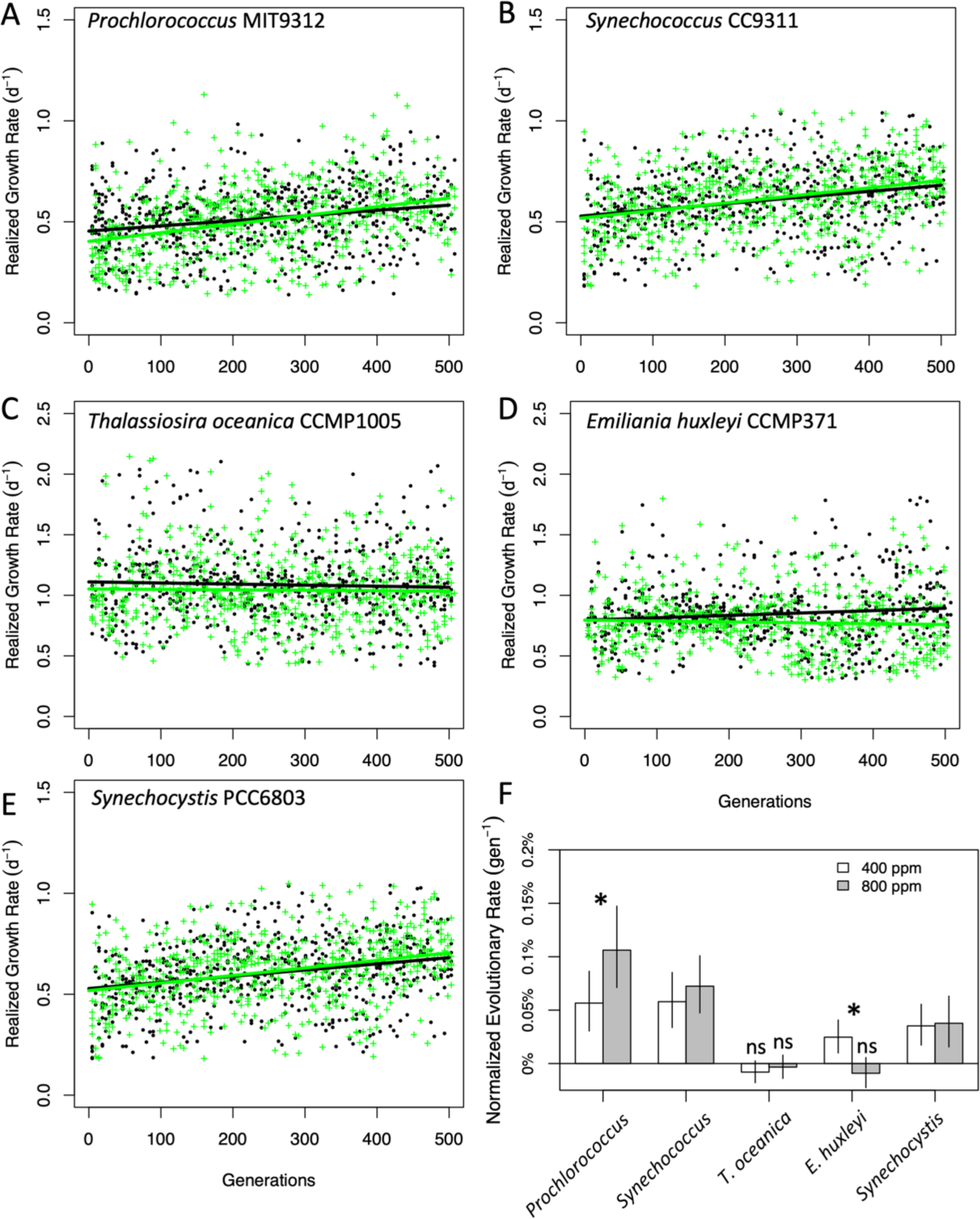
Evolution of the Malthusian growth rates of phytoplankton at 400 and 800 ppm pCO2. A-E. Each point represents the Malthusian growth rate of one transfer (log_2_ 26 = 4.7 generations) of the indicated phytoplankton strain. Trendlines are regressions of growth rate on generation. Black points and lines = 400 ppm cultures, green crosses and lines = 800 ppm cultures. F. Normalized evolutionary rates were expressed as the slope of growth rate change (i.e. regression lines in panels A-E) divided by the estimated ancestral growth rate (i.e. the y-intercept of the regression). Error bars are 95% confidence intervals of the estimates. Asterisks represent p < 0.05 in post-hoc tests of the evolutionary rates of 400 ppm vs. 800 ppm pCO2 cultures. All evolutionary rates were significantly greater than 0 except for those marked “ns” or not significant.

At the end of the experiment, we investigated whether evolution at 400 or 800 ppm pCO_2_ had produced a correlated response (*30*) in MGR and/or exponential growth rate (EGR) in the opposite treatment condition for any of the phytoplankton species (Fig. 2), potentially indicating adaptive specialization to the changed environment. At the beginning of the evolution experiment, *Prochlorococcus* had a significantly slower MGR at 800 ppm (Fig. 1A); however, when the 800 ppm-evolved lineages were grown under both pCO2 conditions, there was no longer a significant difference in MGR between the pCO_2_ treatments (Fig. 2A). In contrast, 400 ppm-evolved *Prochlorococcus*, while growing faster overall than its ancestor, retained significantly reduced MGR and EGR at 800 ppm (Fig. 2A). Evolved *Synechococcus* lineages from both pCO_2_ conditions grew faster than their ancestor, but (as with their ancestor) pCO_2_ treatment had no effect on either their EGRs or MGRs (Fig. 2B). For both *E. huxleyi* and *T. oceanica*, the evolved lineages had greater MGRs and/or EGRs in their evolved milieu than in the opposite pCO_2_ condition (Fig. 2B–C). *T. oceanica* was the only strain to show evidence of adaptive trade-offs, however, with both evolved strains growing more slowly than their ancestor under the pCO_2_ condition opposite from their evolutionary condition (Fig. 2C).

**Figure 2.**
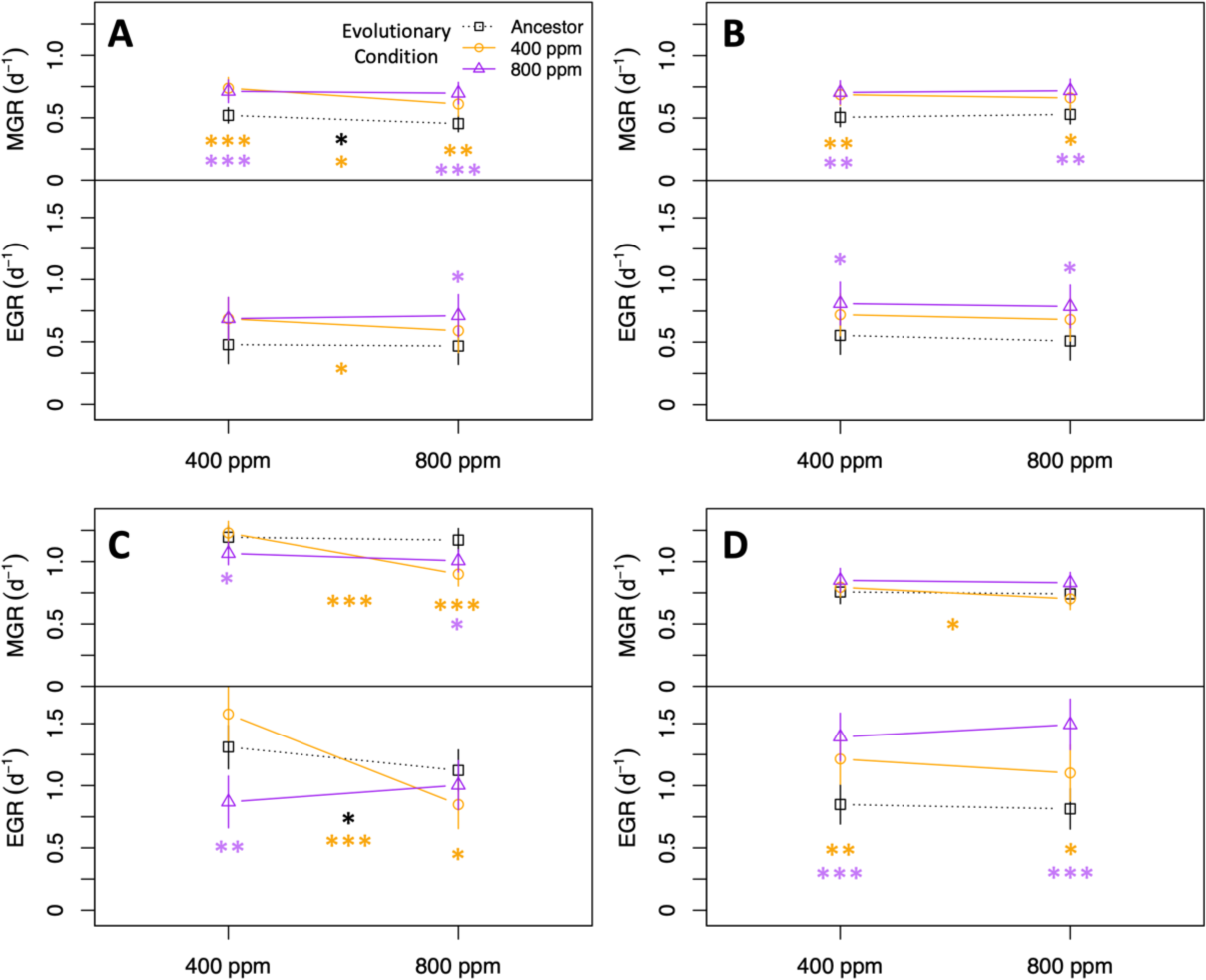
Pre- and post-evolution reaction norms for four phytoplankton species. A. *Prochlorococcus*, B. *Synechococcus*, C. *T. oceanica*, and D. *E. huxleyi*. MGR, Malthusian growth rate; EGR, exponential growth rate. The legend in panel A applies to all panels. Error bars represent 95% confidence intervals for the growth rate estimate. Asterisks in the center of plots indicate significant differences between pCO_2_ treatments; asterisks at ends of reaction norms indicate significant differences between the indicated evolution treatment and the ancestor at the assay pCO_2_. Colors of asterisks correspond to colors of lines. *, p < 0.05; **, p < 0.01; ***, p < 0.001.

To better understand the genetic mechanisms behind the evolution of growth rates and pCO_2_ responses in these organisms, we obtained shotgun metagenomic sequences (>50X coverage for prokaryotes, > 15X coverage eukaryotes, Table S2, Figs. S1–6) for each evolved population and predicted single nucleotide polymorphism (SNP) and simple insertion/deletion (indel) mutations in each of the genomes compared to the reference ancestral genome. We observed thousands of mutations existing at various abundances above our 5% per population cutoff (Table S3), with strikingly different trends for the different species. The three cyanobacterial taxa had several dozen fixed mutations observed across the replicate lineages, with large numbers of rarer mutations (Fig. S7). Similar patterns were observed for *Alteromonas*, albeit with a much greater number of rare mutations relative to fixed ones (Fig. S8). In contrast, *T. oceanica* and *E. huxleyi* genomes showed very large numbers of fixed mutations, and whereas the cyanobacterial non-fixed mutational distributions were skewed toward rarer mutations, the eukaryotes showed a roughly normal distribution of mutation frequencies, centered on 50% (Fig. S9). We believe these differences between bacteria and eukaryotes reflect differences in how natural selection affects sexual and asexual genomes. Because of their obligate asexuality, fixation of a new mutation in a bacterial genome requires a selective sweep, such that each fixed mutation purges all previously existing diversity, generating a bimodal distribution weighted heavily toward rarer, more recent mutations. The eukaryotes, on the other hand, presumably engaged in sexual reproduction at least some of the time, allowing beneficial mutations to fix without a loss of diversity, and at a much greater rate due to an absence of clonal interference (*31*). An accumulation of heterozygotic strains may account for the normal distribution of novel mutations in eukaryotic populations.

The spectrum of mutational types also suggests that natural selection rather than random drift or sequencing errors was responsible for the detected variants. The relative abundance of nonsynonymous and nonsense mutations, both of which result in altered proteins upon translation and are therefore more likely to result in phenotype changes than synonymous or intergenic mutations, varied greatly among phytoplankton taxa (Fig S10) and among *Alteromonas* populations based on which phytoplankter they evolved alongside (Fig S11). The ratio of nonsynonymous to synonymous substitutions (dN/dS ratio) for *Prochlorococcus* was significantly greater than 1 under both pCO_2_ conditions (Fig 3A), suggesting that directional natural selection drove the evolutionary process. In contrast, all other phytoplankton taxa had significantly lower dN/dS ratios than *Prochlorococcus* did, and most were also significantly less than 1, suggesting that purifying selection, not drift, dominated evolution in these cases. Similar trends were observed for the abundance of nonsense mutations, which were very rare for eukaryotes under both pCO_2_ conditions and for *Synechocystis* under current atmospheric pCO_2_ conditions but were significantly elevated for *Prochlorococcus* evolved at 800 ppm pCO_2_; similarly, the transition:transversion ratio (where lower values generally correspond to greater likelihood of phenotypic change) was much higher than expected under neutral conditions for both eukaryotes and *Synechocystis* but was significantly reduced for *Prochlorococcus* at 800 ppm pCO_2_ (Fig. S12Aa–B). In short, evolved *Prochlorococcus* possessed numerous mutations suggestive of changes to protein function especially during adaptation to future predicted pCO_2_, whereas eukaryotic phytoplankton accumulated mutations that were less likely to alter or disrupt protein function and the abundance of these was unaffected by pCO_2_.

**Figure 3.**
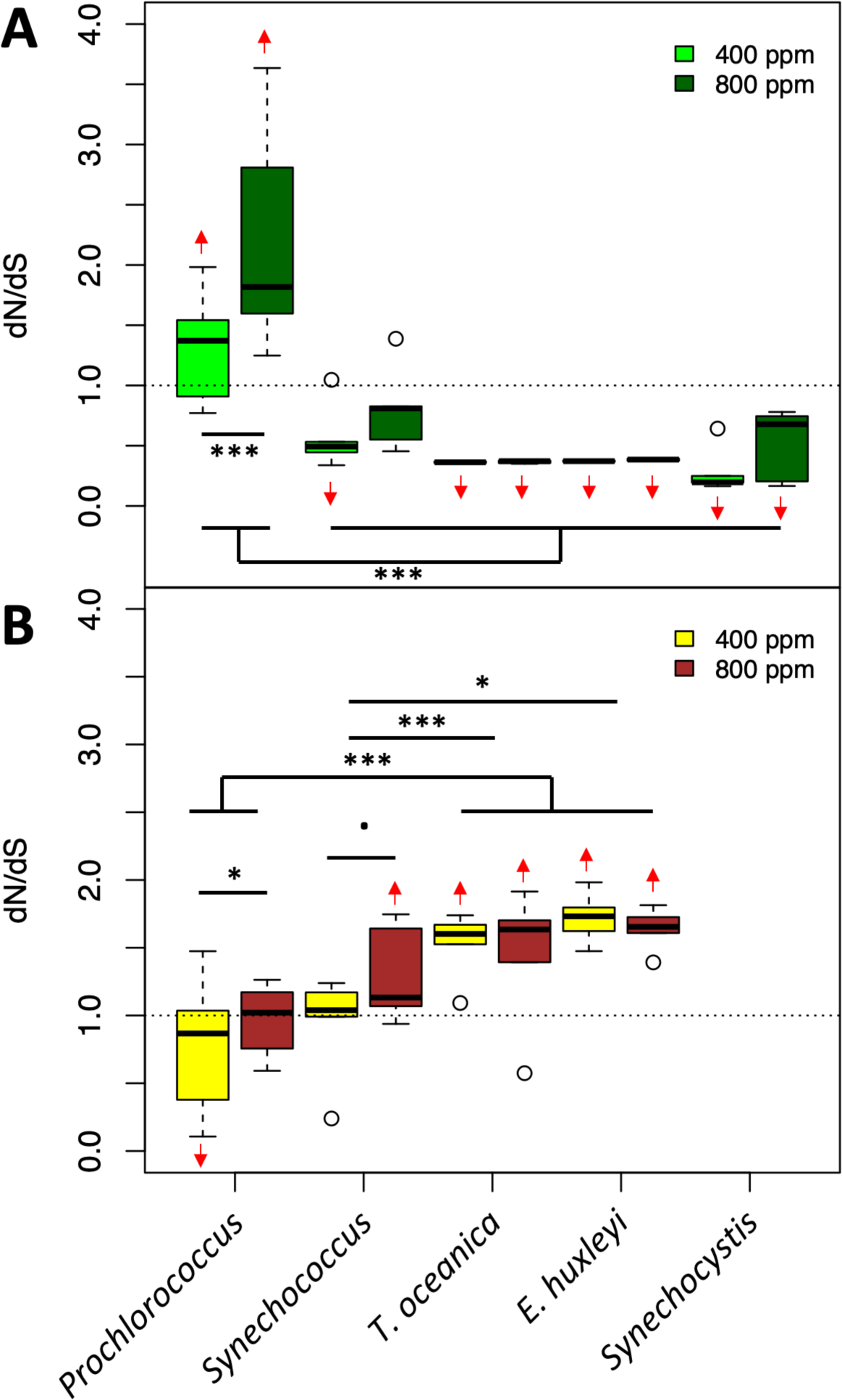
Genomic evidence of adaptive evolution. Plots show the ratio of nonsynonymous to synonymous (dN/dS) amino acid changes in coding sequences of phytoplankton species (A) or *Alteromonas* strains paired with the indicated phytoplankton species (B). Dashed lines indicate the expected value under neutral evolution; red arrows indicate predicted means significantly higher or lower than this expected value (linear model, 95% confidence interval of the extended marginal mean). Asterisks indicate significant differences between pCO_2_ treatments within a species, or between species or groups of species: *, p < 0.05; ***, p < 0.001, **·**, p < 0.1.

In many cases, these trends were reversed for *Alteromonas*: when the phytoplankton partner exhibited evidence of purifying selection and conservation, *Alteromonas* showed signs of directional change, and vice versa (Fig. 3B, Fig. S12C–D). *Alteromonas* EZ55 was originally isolated from a *Prochlorococcus* culture from the same HLII ecotype as the strain used in this study (*32*), so it is not surprising that all three mutational type metrics (i.e., dN/dS ratio, abundance of nonsense mutations, and transition:transversion ratio) support conservation instead of change with *Prochlorococcus* under current pCO_2_ conditions. In general, all these metrics moved in favor of directional evolution as the partner became more evolutionary distant from *Prochlorococcus*. In all cases, these metrics were significantly greater than the value expected by chance for *Alteromonas* evolved alongside eukaryotic phytoplankton. Additionally, the dN/dS and/or nonsense ratios were significantly elevated at 800 ppm pCO_2_ for *Alteromonas* evolved with *Prochlorococcus* and *Synechococcus*, but no pCO_2_ effects were observed either for eukaryotic phytoplankton or the *Alteromonas* strains with which they evolved. We thus conclude that the primary driver of adaptation for *Alteromonas* paired with cyanobacteria was the evolutionary pCO_2_ condition, whereas adaptation to the novel partner drove phytoplankton adaptation in co-cultures with eukaryotes.

To detect specific mutational targets, we searched for genes with more observed mutations than expected by chance. First, we assessed whether overall mutational distribution patterns deviated from random expectations using a Monte Carlo bootstrapping process. In almost all cases, the maximum number of mutations in a single gene observed in the actual dataset was much higher than any bootstrapped dataset (Table S3). However, nonrandom mutational distributions were observed for both nonsynonymous and synonymous mutations, suggesting the presence of mutational hotspots (*33*). We therefore curated a list of individual genes that were most likely to be adaptive evolutionary targets by selecting only nonsynonymous mutations (including mutations within promoter regions up to 50 bp upstream of a prokaryotic start codon) that either i) received more mutations than were observed in any bootstrap dataset or ii) had mutations in at least half of eligible replicate lineages. We further removed any genes that also fit either of these criteria for synonymous mutations.

Using these conservative criteria, relatively few mutations remained for cyanobacteria (Table S4, Fig. S13), although several pathways were nevertheless statistically over-represented (Fig. S14). Interestingly, every *Prochlorococcus* lineage contained one of several promoter-region mutations upstream of the gene for plastoquinol terminal oxidase (PTOX), and most also had similar mutations upstream of thioredoxin reductase; both of these proteins are involved in protection against oxidative stress in cyanobacteria (*34, 35*) and could underlie the increased resilience of evolved axenic MIT9312 cultures (see below). Most *Prochlorococcus* populations also had expansions of AT repeats (Fig. S1) in the promoter/5ʹ region of the apolipoprotein N-acyltransferase gene that encodes an integral membrane protein involved in outer membrane lipoprotein maturation in *E. coli* (*36*), with significantly more polymorphisms observed in 800 ppm pCO_2_ evolved cultures. Lipoproteins are involved in many intercell interactions in bacterial pathogens (*37*) and may affect how *Prochlorococcus* and *Alteromonas* interact with each other and possibly also with secreted membrane vesicles in their environment (*38*). A diverse group of genes were mutated in all *Synechocystis* cultures, with a nitrate transporter being the only gene in these populations significantly more represented in high pCO_2_ treatments. In contrast, no single gene was mutated in more than half of the *Synechococcus* cultures, and only a single uncharacterized protein was differentially mutated between pCO_2_ treatments.

Far more mutations passed the filter for eukaryotic phytoplankton (Table S4, Fig. S13), with genes involved in biosynthetic pathways, carbon metabolism, and peroxisome functions strongly overrepresented (Fig. S15). The diatom *T. oceanica* had dozens of pathways statistically overrepresented, mostly in populations evolved at 400 ppm. In contrast, *E. huxleyi* had more pathways overrepresented in lineages evolved at 800 ppm pCO_2_, including several pathways related to lipid metabolism. Interestingly, most genes that passed our cutoff for *T. oceanica* only did so in one or the other pCO_2_ treatment, whereas most *E. huxleyi* multiply mutated genes passed the filter under both treatments (Fig. S13).

The mutational targets for *Alteromonas* differed dramatically between populations evolved with different phytoplankton partners (Table S5, Fig. S16). Relatively few mutations and pathways passed our filters in strains paired with cyanobacteria, consistent with our observation of mostly purifying selection under these conditions for *Alteromonas* except when paired with *Prochlorococcus* under 800 ppm pCO_2_. When evolved alongside eukaryotes, however, many pathways were found to be significantly over-represented in the mutational data (Fig. S17), and conspicuously more genes were observed to be multiply mutated under 800 ppm than under 400 ppm pCO_2_ (Fig. S16). In general, all these metrics moved in favor of directional evolution as the partner became more evolutionary distant from *Prochlorococcus*. Regardless of the phytoplankton partner or pCO_2_ treatment, two-component regulatory systems and genes related to chemotaxis and flagellar synthesis were convergently mutated across *Alteromonas* lineages, consistent with these genes previously being shown to be differentially regulated in ancestral *Alteromonas* based on partner and pCO_2_ conditions (*8*). The most mutated EZ55 gene was an unannotated putative lipoprotein (EZ55_02425) observed in 47 out of 48 *Alteromonas* lineages, with almost all mutations falling within the 50-bp promoter region we considered in our analysis, suggesting possible cell-surface alterations like those we speculated above may have occurred in *Prochlorococcus*.

Relatively few *Alteromonas* genes that passed our filter were shared between *Alteromonas* evolved with *Prochlorococcus* vs. *Synechococcus* or between *Alteromonas* evolved with cyanobacteria vs. eukaryotes, whereas a large suite of genes was shared between *Alteromonas* partnered with each of the two eukaryotic phytoplankton (Table S6, Fig. S18), with strong overrepresentation for genes involved in amino acid metabolism (Fig. S19). Genes that were significantly more mutated with a single partner tended to be regulatory or sensory gene, but also included several specific carbon catabolism genes, which implies variability existed in the carbon compounds exuded by the different phytoplankton (Table S6). Frequently, genes that were specific to a given partner were also differentially mutated between pCO_2_ treatments (Table S6). For instance, the *Synechococcus*-specific gene fucose permease was much more likely to be mutated at 800 ppm pCO_2_, as were eukaryote-specific regulatory genes.

Substantial evolution of *Alteromonas’* single plasmid was also detected, with most evolved populations having significant accumulations of plasmid-free segregants (Fig. S20) or possible insertion of a fragment of the plasmid by homologous recombination into the primary chromosome (Fig. S21). Notably, most populations evolved alongside *Synechococcus* lost the free plasmid entirely, whereas most populations evolved with *E. huxleyi* appear to have retained it intact, with those evolved at 800 ppm pCO_2_ in some cases carrying multiple plasmid copies per cell (Fig. S20B). Overall, *Alteromonas* evolved with cyanobacteria were significantly more likely to lose the free plasmid entirely than were those evolved alongside eukaryotes (Fisher’s exact test, p = 0.009, Fig. S20B). Coverage patterns also suggested the evolution of a sub-population of plasmids of reduced size in some lineages (Fig. S20C), perhaps indicating negative frequency-dependent interactions between plasmid carriers and segregants (*39*). The plasmid primarily consists of transposable elements, metal and antibiotic resistance genes, and genes for metabolizing toluene and xylene (*40*), so it is unclear what selective pressures underlie these trends.

Collectively, these observations suggest that the five phytoplankton and *Alteromonas* were experiencing varying levels of natural selection largely working to alter their carbon metabolism. There was relatively little evidence that mutational responses in the phytoplankton were CO_2_ level-specific, even though *Prochlorococcus* had been able to effectively eliminate its high pCO_2_ growth defect after evolving at 800 ppm for 500 generations. Instead, for the four species co-cultured with *Alteromonas*, most of the convergent mutations for both *Alteromonas* and phytoplankton were consistent with the organisms adapting their metabolism to their partner. *Alteromonas* clearly experienced stronger selection as its chemical and biological environment became more divergent from its ancestral condition as a long-term partner of *Prochlorococcus*, but its strongest and most diverse evolutionary responses were observed in co-culture with eukaryotic phytoplankton at projected year 2100 pCO_2_. Of the phytoplankton, only *Prochlorococcus* showed strong evidence of general directional selection (Figs. 3, S12), and the fact that multiple redox-oriented targets were detected suggested that the “helping” activity of *Alteromonas* may have been downgraded during evolution, forcing *Prochlorococcus* to evolve stronger antioxidant defenses of its own potentially by co-option of thioredoxins and PTOX proteins. The large differences between the mutational spectra of *Alteromonas* with cyanobacteria compared to that with eukaryotic phytoplankton also raised the possibility that *Alteromonas* may specialize on particular partners, which might also impact its “helper” ability.

We therefore sought to measure changes in *Alteromonas’* “helping” ability by conducting a series of experiments where various combinations of evolved and ancestral *Prochlorococcus* and *Alteromonas* were paired in co-culture. First, we discovered that evolved *Prochlorococcus* cultures retained a need for “help” from *Alteromonas*, with significantly elevated mortality risk in axenic culture and at elevated pCO_2_ even after 500 generations of evolution (Fig. 4A, S22). However, the magnitude of this impediment had significantly decreased during evolution, suggesting that at least some protective mutations (e.g., the PTOX and thioredoxin reductase regulatory mutations described above) had arisen in *Prochlorococcus* genomes. We also found that the community context under which *Alteromonas* evolved affected its helping ability. *Alteromonas* isolates from the evolved *Prochlorococcus* cultures (regardless of evolutionary pCO_2_ treatment) were generally less effective as helpers of ancestral *Prochlorococcus*, allowing greater mortality than ancestral *Alteromonas* (Fig. 4A). Moreover, whereas the *Alteromonas* ancestor increased the *Prochlorococcus* growth rate relative to axenic cultures, the evolved *Alteromonas* isolates either did not enhance or even decreased *Prochlorococcus* growth rates relative to axenic cultures for *Alteromonas* evolved alongside *Prochlorococcus* and eukaryotes, respectively (Figs. 4B, S23).

**Figure 4.**
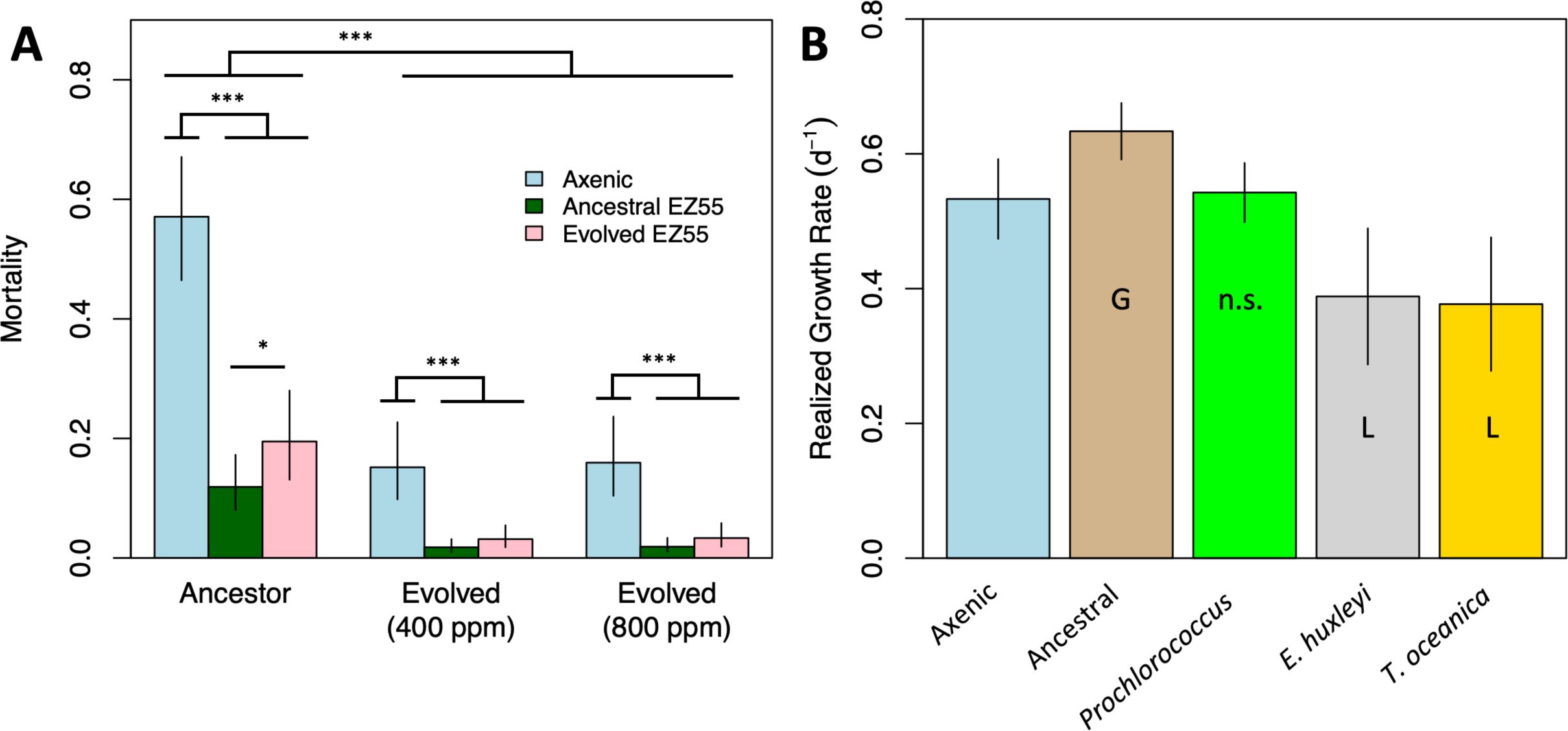
Effects of experimental evolution on the “helping” ability of *Alteromonas* EZ55. A) Bars represent model estimates from a logistic mixed effects regression model predicting the likelihood of culture failure/death for *Prochlorococcus* cultures grown either axenically or as co-cultures with ancestral EZ55 or with EZ55 evolved with *Prochlorococcus*. Because EZ55 clones from both pCO_2_ treatments had statistically similar effects on mortality reduction, they are included together in these estimates. The impact of pCO_2_ on mortality was independent of the EZ55 treatment and is shown in Figure S22. *, p < 0.05; ***, p < 0.001. B) *Prochlorococcus* was grown either axenically, in co-culture with ancestral EZ55, or with clones of EZ55 isolated from cultures of *Prochlorococcus*, *E. huxleyi*, or *T. oceanica* after 500 generations of evolution at 400 ppm pCO_2_. G and L indicate significantly (p < 0.05) greater or lower growth rates based on the results of a Dunnett’s test comparing each EZ55 treatment to the axenic control, whereas n.s. indicates the result of the comparison was nonsignificant. Error bars represent the 95% confidence intervals of the extended marginal means estimate of the growth rate.

A possible explanation for changes in *Alteromonas’* interactions with *Prochlorococcus* was revealed by examining the correlational network of mutations in the *Alteromonas* genome across lineages (Fig. 5). For 17 genes in the *Alteromonas* genome, the presence of mutations in one gene was significantly predictive of mutations in at least one other gene. Most of these were genes related to transcriptional regulation, environmental sensing, or transmembrane transport, and many included frameshift or nonsense mutations, suggesting that the adaptive benefit of the mutations may involve loss of function. The gene with the most incoming predictive edges was only found mutated in *Alteromonas* paired with eukaryotic phytoplankton, and it encodes a gene matching the *aceK* phosphatase/kinase that regulates the switch between the full TCA cycle and the abbreviated glyoxylate shunt in *E. coli*.

**Figure 5.**
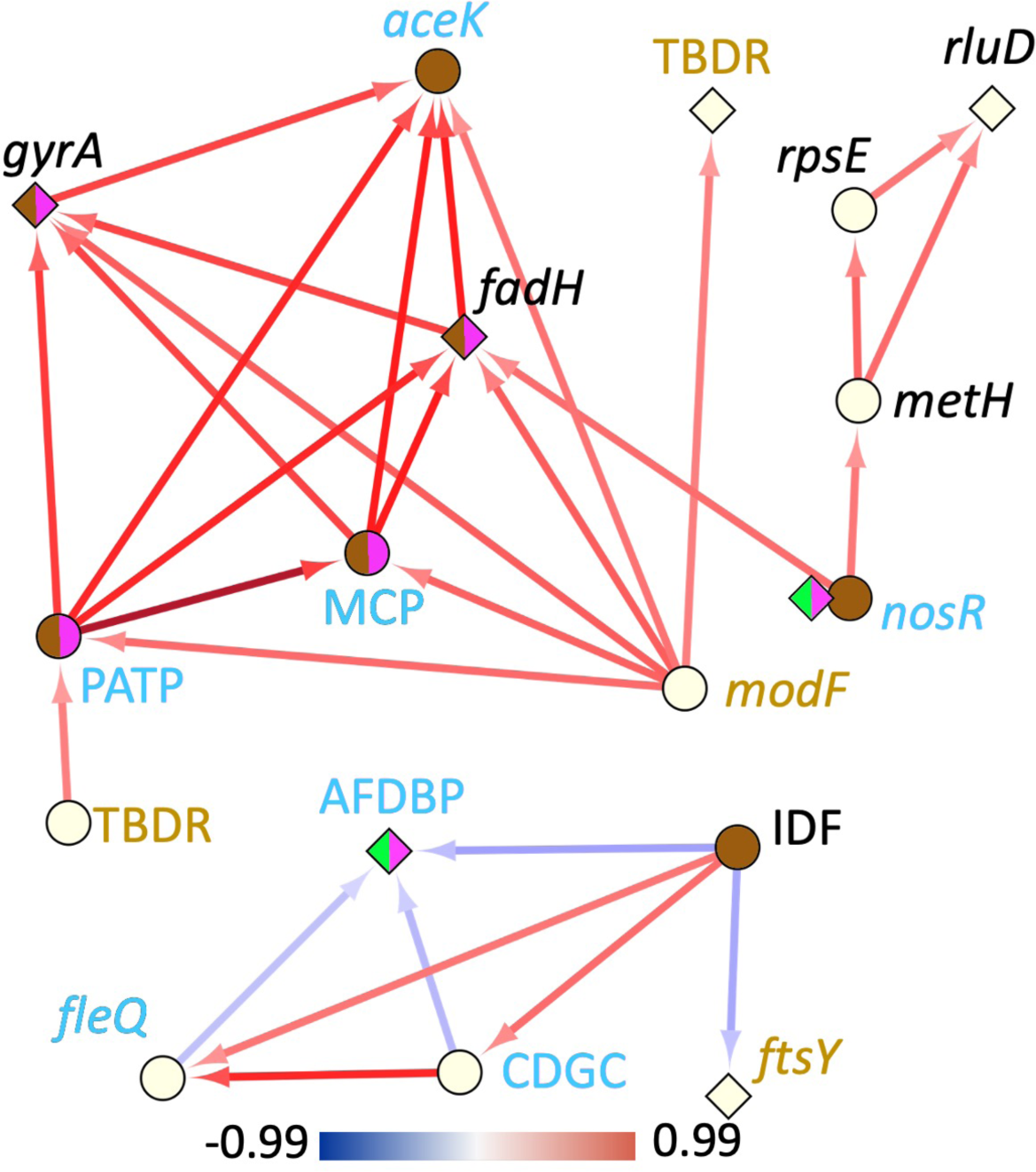
Mutational network of *Alteromonas* EZ55. Nodes indicate genes and arrows indicate that mutations in one gene significantly predict mutations in the second gene (at the end of the arrow) in the same evolved EZ55 culture. Diamond nodes indicate genes where only nonsynonymous or promoter region mutations were observed; circular nodes indicate genes with potential knock-out mutations (e.g., frameshifts and nonsense mutations). For *nosR*, mutational profiles were very different for cyanobacteria and eukaryotes, so two node types are included to reflect this. Node color indicates in which phytoplankton co-cultures mutations were observed: green, *Prochlorococcus*; pink, *Synechococcus*; brown, both eukaryotic strains. Gene names in italics represent the names of the closest match in *Escherichia* or *Pseudomonas* genomes; capitalized letters represent genes without clearly annotated identities. TBDR, TonB dependent receptor; MCP, methyl-accepting chemotaxis protein; PATP, PepSY associated transmembrane protein; AFDBP, Arc family DNA binding protein; CDGC, Cyclic-di-GMP cyclase. Gene names in gold have products involved in transmembrane transport; names in blue have products involved in environmental sensing and transcriptional regulation. Arrow color reflects the Spearman correlation coefficient between the two genes according to the color bar at the bottom of the figure.

Phosphorylation of isocitrate dehydrogenase by AceK bypasses the loss of 2 C atoms during oxidative growth, allowing growth on simple compounds such as acetate or glycolate (*41*). Mutations in genes related to growth on complex carbon compounds strongly predicted mutations in *aceK* in eukaryotic cultures, suggesting a shift away from simple carbon substrates as *Alteromonas* evolves away from partnering with *Prochlorococcus*. Previous work suggested that catabolism of small organic acids was an important component of *Alteromonas’* response to short-term co-culture with both *Prochlorococcus* and *Synechococcus* (*8*), so the inactivation of a key pathway for metabolizing these compounds in long-term co-culture with eukaryotes is particularly striking. Potentially loss of function mutations in genes related to denitrification (e.g., molybdenum transport gene *modF* and nitrite reductase gene *nosR*) were also predictive of mutations in the cluster of genes connecting to *aceK*, possibly signaling changes in *Alteromonas’* N demand when co-cultured with eukaryotes. These changes are suggestive of broad alterations in core metabolic processes as *Alteromonas* adapts to partners other than *Prochlorococcus* and could underlie the apparently antagonistic relationship between *Prochlorococcus* and eukaryote-evolved *Alteromonas*.

We draw three central conclusions from the results of our evolution experiment. First, all phytoplankton we observed evolved rapidly under both pCO_2_ regimes, in most cases increasing their growth rates significantly in the milieu in which they evolved. *Prochlorococcus*, which had the most negative response to future pCO_2_ conditions prior to evolution, was able to compensate for its growth deficiencies within 500 generations, suggesting that *in situ* populations will be able to evolve fast enough to avoid major changes to their range in coming decades (e.g. (*5*)). Second, in the case of the diatom *T. oceanica*, we measured growth rate evidence of physiological trade-offs to adaptation to high pCO_2_ growth that resulted in a decrease in growth rate in the ancestral environment. Further work will be necessary to understand the metabolic basis for this trade-off, but it is possible that the trade-offs will cause unexpected secondary effects that may have important consequences for community function and nutrient cycling. That more convergently mutated genes were observed between *Alteromonas* partnered with *T. oceanica* and *E. huxleyi* at 800 ppm than at 400 ppm suggests that metabolic rearrangements may be more common among marine microbes at projected year 2100 pCO2, and this could have downstream impacts for the broader marine food web.

Third and finally, our results conclusively demonstrate that the “helper” interaction between *Alteromonas* and *Prochlorococcus* is not strictly or stably mutualistic. Instead, we saw evidence that each organism adapted to improve its own fitness and potentially disentangle itself from its partner. For instance, *Prochlorococcus* clearly became less dependent on *Alteromonas* for stress protection, evidenced by its reduced mortality risk in axenic culture compared to its ancestor (Fig. 4A). At the same time, *Alteromonas’* capacity to protect *Prochlorococcus* decreased even in clones isolated from evolved *Prochlorococcus* cultures, and this effect was more pronounced after co-evolution with more distantly related phytoplankton partners (Fig. 4B). In previous work involving short-term co-cultures (*6, 8*) we showed that *Prochlorococcus*’ poor growth at 800 ppm occurred because *Alteromonas* decreased expression of its catalase genes under these conditions, possibly because of changes in the carbon compounds *Prochlorococcus* excreted under a higher CO_2_:O_2_ ratio atmosphere. We hypothesized that, over evolutionary time, *Alteromonas* might adjust its gene expression to restore its “helper” ability at 800 ppm pCO_2_, but instead the evidence suggests that the two partners evolved greater independence from each other. This result is consistent with the concept that the outwardly mutualistic interaction between *Prochlorococcus* and helpers like *Alteromonas* is a result of reductive Black Queen evolution (*42, 43*). Whereas true mutualisms are positively reinforcing, Black Queen mutualisms are the result of dynamic equilibria between a more efficient, streamlined beneficiary and a helper. Laboratory experiments show that these equilibria fluctuate as one partner or the other obtains beneficial mutations during evolution (*39, 44*). Both chemostat (*45*) and transcriptomic evidence (*8*) indicate that *Alteromonas* competes for N with phytoplankton in co-culture, so it is reasonable to expect that its apparent “helping” ability should decrease as it evolves to more efficiently compete with *Prochlorococcus*, but that the general dynamic would persist. An open question, however, is why *Prochlorococcus* in nature does not appear to favor the putative antioxidant mutations observed in this experiment, but instead maintains a much higher degree of vulnerability to oxidative stress than is strictly necessary given its genomic capacities.

In conclusion, this work demonstrates that phytoplankton are sufficiently evolutionarily plastic that results from short-term experiments are unlikely to be strong predictors of the behavior of taxa under future pCO_2_ conditions, and likely other environmental changes such as warming or mixed layer shoaling as well. However, some taxa may be extensively modified during this adaptation as evidenced by the hundreds of nonsynonymous mutations sampled during this relatively short period, and metabolic changes are likely to alter the character of exuded photosynthates which will result in compensatory alterations in pelagic bacterial communities with unknown impacts on the rest of the food web. Finally, we see evidence of rapid diversification and partner specialization in *Alteromonas*, suggesting that this ubiquitous taxon may undergo similar evolution in nature in response to local phytoplankton communities. Future work should explore the mechanisms underlying the shift for *Alteromonas* between cyanobacteria and eukaryote specialization, and why this shift results in apparently exploitative interactions with *Prochlorococcus*. We propose that the evolved organisms from this experiment present an opportunity for future investigators to study this phenomenon and other aspects of algal:bacterial co-evolution in greater detail.

## Acknowledgements

We are grateful to Erik Zinser, Steven Wilhelm, and Sonya Dyhrman for assistance with obtaining strains and developing methods; to Irene Chiang for laboratory assistance; to Michael Crowley and the UAB Genomics Core for sequencing; to Jeffrey Barrick for assistance with breseq; and to the UAB Cheaha HPCC and staff for computing resources. This project was supported by grants OCE-1540158 and OCE-1851085 from the National Science Foundation, an early career fellowship in Marine Microbial Ecology and Evolution from the Simons Foundation, and startup funds from the UAB College of Arts and Sciences.

## Materials and methods

### Cultures and media

The phytoplankton used in this study as well as the media in which they were grown are listed in Table S1. All media types were derived from media commonly used to cultivate each organism (*46*). P concentrations were standardized at 2 mM NaH_2_PO_4_ across all media types, with N added at Redfield proportions (16:1 N:P). *Prochlorococcus* was grown in PEv medium, which was a 1/25 dilution of Pro99 (32 mM NH_4_Cl, 40 µL L^-1^ Pro99 trace metals). *Synechococcus* was grown in SEv medium, which was modeled on SN, but with N concentrations lowered to be at Redfield proportions with P (final concentrations 32 mM NaNO_3_, 20 µL L^-1^ SN trace metals, 20 µL F/2 vitamins). *T. oceanica* and *E. huxleyi* were grown in FEv medium, which was a 1/25 dilution of F/2 medium, replacing the F/2 trace metal solution with 40 µL L^-1^ of L1 trace metals. All media were prepared in an artificial seawater (ASW) base (per L, 28.4 g NaCl, 7.21 g MgSO_4_*7H_2_O, 5.18 g MgCl_2_*6H_2_O, 1.58 g CaCl_2_, 0.79 g KCl). This ASW base was autoclaved in 1 or 2L batches, then filter-sterilized nutrient solutions and 4 mL of filter-sterilized ∼0.58 M NaHCO_3_ solution were added. The precise concentration of NaHCO_3_ of each batch was determined by titration (see below). Completed media were bubbled for at least 24 h with sterile air to equilibrate the carbonate system with the atmosphere. All media storage bottles and culture glassware were acid washed prior to use.

Prior to use in experiments, phytoplankton cultures were rendered clonal and axenic. First, cultures were diluted to approximately 1 cell mL^-1^ and then 100 µL aliquots were distributed to 96-well plates, such that each well was likely to have either 0 or 1 cells; replicate ancestral cultures of each strain were picked from these plates, cryopreserved (see below), and used for all subsequent experiments. We used a streptomycin-resistant *Prochlorococcus* strain (*32*) and each of our six ancestral cultures was rendered axenic according to (*29*), then checked for bacterial contamination by adding 1 mL aliquots to liquid YTSS medium (*47*); if no growth was observed after 48 h, the *Prochlorococcus* cultures were diluted 100-fold into fresh PEv medium to initiate experiments. *Synechococcus* and *Synechocystis* were made axenic by dilution-to-extinction. *T. oceanica* and *E. huxleyi* were both axenic upon receipt from the National Collection of Marine Algae (Boothbay Harbor, ME).

*Alteromonas* sp. EZ55 was streaked for isolation on YTSS agar and individual colonies were used to inoculate clonal liquid YTSS cultures. Prior to addition to axenic phytoplankton, *Alteromonas* cells were pelleted by centrifugation (2000 × *g* for 2 min) and washed twice with sterile saline (8 g L^-1^ NaCl). *Alteromonas* clones were re-isolated from post-evolution cultures by spread plating on YTSS agar.

### Carbonate system manipulation

We manipulated the carbonate system in our cultures by careful additions of HCl, NaHCO_3_, and/or NaOH (*48*). The exact concentration of each solution was determined by titrating the alkalinity of ASW before and after addition of gravimetrically determined masses of solution. Alkalinity titrations were performed with factory-standardized 0.1 N HCl using a Mettler Toledo T5 titrator according to (*49*). Medium pH was determined by addition of m-cresol purple according to (*49*), but with the protocol modified to use a BioTek Synergy H1 plate reader instead of a standard spectrophotometer. Using the measured pH and alkalinity of a batch of media, we calculated the additions of either HCl and NaHCO_3_, or NaOH, necessary to achieve 400 ppm or 800 ppm pCO_2_ conditions using the package *seacarb* in R (*50*). We used calibrated 0.2 M NaHCO_3_, 0.2M HCl, and 0.1M NaOH solutions for carbonate system manipulation. All solutions were filter sterilized prior to calibration and were monitored for bacterial contamination by periodically adding aliquots to YTSS broth.

In general, we only assessed the alkalinity and atmosphere-equilibrated pH of a 2L batch of media once, prior to its utilization for experiments. However, approximately one year into the experiment we discovered that the atmospheric pCO_2_ in the laboratory varied considerably on a seasonal basis, which led to some drift in pCO_2_ concentrations in the media over the period necessary to consume an entire 2L batch. We therefore began monitoring laboratory CO_2_ levels and re-assessed pH (and re-calculated necessary additions) when substantial shifts in CO_2_ were observed. To the best of our knowledge these fluctuations did not produce pH deviations greater than 0.1 unit from our intended targets and did not result in overlap between our ambient and year 2100 pCO_2_ treatment groups.

### Cryopreservation of cultures

Using dilution-to-extinction, we isolated 5 (PCC6803) or 6 clones (all other strains) of each phytoplankton strain to initiate evolution experiments. We also isolated 6 *Alteromonas* clones from isolated colonies on YTSS agar. These clonal populations represent the ancestors of our experiment and were each cryopreserved immediately. *Alteromonas* clones were frozen at −80°C in YTSS with 20% sterile glycerol. Cyanobacteria were flash frozen in culture media with 7.5% sterile DMSO by immersion in liquid nitrogen (*51*). Eukaryotic phytoplankton were also treated with 7.5% DMSO, but were slowly frozen using a Mr. Frosty device according to the manufacturer’s instructions (*46*). After the freezing process, all cultures were placed in long-term storage in liquid nitrogen vapor.

When necessary, *Alteromonas* cultures were revived by scraping some frozen material from the top of the sample with a sterile wooden dowel and streaking for isolation onto YTSS agar; experiments were always initiated from fresh clonal isolates. Phytoplankton were revived by placing frozen samples into room-temperature water in the dark just long enough to fully thaw. Then, working under very low light, 500 µL were inoculated into fresh media, and the tube was refrozen using the same techniques described above. Freshly revived cultures were placed into an incubator under very low light (∼5 µmol photons m^-2^ s^-1^) for 48 h, then moved to moderate (∼30 µmol photons m^-2^ s^-1^) light and monitored for growth by flow cytometry (see below).

### Experimental evolution

Cultures for experimental evolution were initiated with 12.3 mL of media, 0.2 mL of acid or base additions for carbonate chemistry manipulation, and 0.5 mL of a previous culture. All cultures were grown at 22°C under approximately 75 µmol photons m^-2^ s^-1^ in acid-washed conical-bottom glass tubes with airtight caps; with 13 mL of culture, almost no headspace existed in these tubes. All cultures except *Prochlorococcus* were grown on a rotating test tube wheel with illumination from the top and bottom; preliminary observations indicated that *Prochlorococcus* cells did not settle noticeably when grown in static test tube racks, whereas all other taxa did. Each phytoplankton clone was split into two culture lines, one maintained at 400 ppm pCO_2_ and the other at 800 ppm pCO_2_. For all strains except *Synechocystis*, each clone was co-cultured with a single *Alteromonas* clone. Thus, all phytoplankton designated “1” in Table S1 were co-cultured with *Alteromonas* clone 1, all phytoplankton labeled “2” were co-cultured with *Alteromonas* clone 2, and so forth.

Phytoplankton growth was measured every 48 h using a Guava HT1 flow cytometer equipped with a 488 nm laser. Phytoplankton populations were identified by their clustering pattern on logarithmic plots of forward light scatter vs. 660 nm (chlorophyll) fluorescence. Cell densities within user-defined gates encircling the phytoplankton were calculated automatically by the Guava software. When phytoplankton cell densities crossed a cutoff value (Table S1), cultures were diluted 26-fold (0.5 mL into a total volume of 13 mL) into fresh media. We targeted 108 transfers for each lineage, representing log_2_26 or 4.7 generations (although we did not achieve this goal for some lineages due to repeated crashes or contamination, Table S1). Samples from each lineage were cryopreserved every 25 generations and again at the end of the experiment.

We monitored both the media, media additions, and cultures for external contamination throughout the experiment. Each time transfers were performed, 0.5 mL of any media batch used were added to 5 mL of YTSS broth and monitored for at least 7 days for evidence of growth. *Synechocystis* cultures, which were grown axenically, were directly tested for bacterial contamination by transferring 0.5 mL into YTSS broth after transfer. For cultures containing *Alteromonas*, we periodically spread-plated cultures on YTSS media and examined colony morphology for evidence of bacteria other than *Alteromonas*, which forms distinctive large, shiny brown colonies.

After transfer into fresh media, the previous generation’s tube from a given replicate evolutionary lineage was placed back in the same incubator under low (<30 µmol photons m^-2^ s^-1^) light conditions to slow growth. At least three previous transfers were stored in this manner. In rare instances (see Table S1) when a culture failed to grow after transfer or contamination was detected in a culture or in the media, we re-started the line from the last trustworthy transfer tube. In a small number of cases, we had to revive a culture from the last cryopreserved sample due to slow-growing contaminants.

### Growth experiments

At the end of the evolution period, we subcultured clonal evolved *Alteromonas* strains by spread-plating evolved cultures on YTSS agar and selecting single, isolated colonies for growth in YTSS broth. *Prochlorococcus* was rendered axenic by addition of 100 µg mL^-1^ streptomycin. All growth experiments were initiated by mixing axenic *Prochlorococcus* with a specific *Alteromonas* clone (or else remaining axenic) and acclimating the co-culture for 3 transfer cycles (approximately 14 generations) at the target pCO_2_ concentration. Growth was then monitored by flow cytometry as described above for at least 3 subsequent transfers under constant conditions. We calculated realized and exponential growth rates as well as lag duration as described in (*6*).

### Statistical analysis of growth data

The impact of experimental treatments on growth parameters was statistically analyzed using linear models in R with post-hoc statistical testing using extended marginal means with the *emmeans* package (*52*). Malthusian and exponential growth rates were calculated as described in (*6*). Because these experiments involved thousands of measurements collected over several years, a variety of clearly erroneous data points were recorded that led to several outlier growth rates that had a disproportionate impact on model output; rather than attempt to manually curate all growth rates, we simply removed either the most extreme 5% high and low Malthusian growth rates or eliminated exponential growth rates with r^2^ values lower than 0.95 for each strain in each experiment before conducting statistical tests. Because growth rates evolved differently in different lineages, contrasts involving evolved strains were performed using linear mixed effects models with the *lme4* package in R (*53*) with lineage as the random contrast.

### Whole genome re-sequencing

Post-evolution cultures were split into 5 replicate 13 mL culture tubes, grown to the cutoff transfer cell density, and then collected by gentle vacuum filtration on 0.2 µm pore size polycarbonate filters, then flash frozen in liquid nitrogen and stored at −80°C. Genomic DNA was extracted from the filters using MoBio ProSoil kits, with the bead-beating step accomplished using a MP FastPrep-24 homogenizer. DNA was fragmented, ligated with Illumina adapters, and sequenced on an Illumina NextSeq500 device.

FastQ files (NCBI BioSamples SAMN34542194 through SAMN34542251) were analyzed using breseq (*54*) with default settings. *Synechocystis* PCC6803 cultures were assembled against the chromosomal reference genome (NCBI accession number NC_000911) as well as its four plasmids (NC_005229, NC_005230, NC_005231, and NC_005232). *Prochlorococcus* MIT9312 and *Synechococcus* CC9311 were assembled against their RefSeq genomes (respectively, CP000111 and NC_008319). The *T. oceanica* CCMP1005 and *E. huxleyi* CCMP371 reference genomes were still in draft form and were assembled in breseq with the contig flag activated; all contigs were downloaded as a single GenBank format file from NCBI BioProject PRJNA36595 (CCMP1005) or BioSample SRX112492 (CCMP371, also known as strain 12-1). *E. huxleyi* CCMP371 (also known as strain 12-1 (*55*)) The *T. oceanica* CCMP1005 sequence file contained the *T. oceanica* chloroplast sequence but *E. huxleyi* CCMP371 did not; we therefore used the chloroplast sequence of another *E. huxleyi* strain, CCMP373, also with downstream curation, to analyze mutational changes to those sequences. For both *T. oceanica* and *E. huxleyi*, breseq was set to not attempt to predict structural variants due to the much greater computational demand required to assemble these large genomes relative to the bacteria.

*Alteromonas* genomes were assembled against the most recent EZ55 genome (*40*), including both the chromosomal (CABDXN010000001) and plasmid (CABDXN010000002) sequences in the same GenBank format file. For *Prochlorococcus* and *Synechococcus* lineages, EZ55 genomes were assembled in the same breseq run as the phytoplankton genome. However, because of the computational demand of assembling *T. oceanica* and *E. huxleyi* genomes, the *Alteromonas* portion of these cultures was assembled separately to allow for structural variant prediction.

Any specific mutational call present in 100% of replicate lineages for a given organism was considered to have occurred prior to the initiation of the experiment (i.e., it was ancestral) and was removed from further consideration. Because of additional genomic complexity in the eukaryotic genomes of *T. oceanica* and *E. huxleyi*, we undertook additional efforts to curate these datasets. First, coverage for the eukaryote genomes was lower than for the bacterial genomes, and some loci had insufficient coverage to make predictions about mutations; all these loci were removed from breseq output files prior to further analysis. Also, because the GenBank files we used as reference sequences represented haploid sequences, we sought to discover potentially heterozygous loci in the ancestral genomes. To do this, we re-assembled the raw read files from the original genomic sequencing runs for both organisms against the GenBank reference sequences using breseq (SRX112492 for *E. huxleyi* and all sequencing runs within BioProject PRJNA36595 for *T. oceanica*). We identified several variants present in 100% of reads compared to the published GenBank sequences, perhaps suggesting differences between breseq’s handling of data and the programs used to produce the published sequences; all mutations assigned by breseq to these loci were therefore removed from consideration as untrustworthy. All variants present in 0% < *n* < 100% of sequences in the reference genome reassemblies were assumed to represent loci that were heterozygous in the ancestral population. Mutations mapped to these loci were only considered further if they became fixed in evolved lineages; if they remained at intermediate frequencies they were removed from subsequent analyses. Mutational frequency analysis indeed suggested (Figs. S7–S9) that *T. oceanica* and *E. huxleyi* retained substantial levels of heterozygosity during the experiment and were possibly undergoing frequent sexual reproduction.

### Mutational analysis

We used the application gdtools from the breseq package to convert breseq output genome difference files into long-format data files for subsequent analysis. Custom python scripts were used to bin all mutations within a given gene in each lineage, producing mutational count tables of genes vs. lineages. For bacterial genomes, we produced count tables either with or without considering insertions, deletions, and intergenic mutations within 50 bp upstream of a gene’s start codon (i.e., putative cis-acting promoter or other regulatory elements) in addition to mutations within a gene’s coding region. Eukaryotic count tables only considered coding sequences; all intergenic and intron regions were removed from analysis. We used python scripts to analyze three metrics of directional selection: dN/dS (*56*), transition:transversion ratio (*57*), and nonsense mutation ratio (*58*). All three metrics exclusively used SNP data from coding regions of annotated genes. Effects of treatment groups on these metrics were analyzed using linear models in R.

We sought to test for convergent evolution of gene targets by determining which, if any, genes received more mutations than would be expected by chance under a completely random mutational model. This analysis was complicated by the fact that larger genes represented larger mutational targets than smaller ones, precluding the use of a simple Poisson model. Instead, we used a bootstrapped Monte Carlo procedure to produce randomized genomes with the same number of coding sequence mutations observed in our real dataset, only distributed randomly across the genome. Given the total coding genome size *g* as the sum of the lengths of all coding sequences in the genome, and the total number of observed mutations *n*, the probability *p* of any given base pair receiving a mutation is *p* = *n*/*g*, and the probability *λ* of a gene of length *l* receiving a mutation is thus *λ* = *p x l*. For each lineage, we produced 100 randomized matrices where each gene in the genome was assigned several mutations drawn from a Poisson distribution with average rate of occurrence *λ*. Each random matrix was compared to the real matrix of mutations by first converting each into an empirical cumulative distribution function (ECDF) and then applying either a Kolmogorov-Smirnov or dts test (*59*). The Kolmogorov-Smirnov (KS) test only considers the single value in the ECDF where the gap between the two samples is greatest, whereas the dts test considers the entire distribution. Because the greatest difference between our observed and Monte Carlo matrices was the fact that the real dataset had many more genes with large numbers of mutations than were ever observed in any simulation, the dts test was generally more sensitive in detecting significant differences. These procedures were conducted separately for nonsynonymous and synonymous mutations and revealed that the distribution of mutations was significantly different from random expectations.

Because genomes are known to include mutational hot-spots (*33*), we performed additional curation of multiply-mutated genes to find genes that were most likely targets of natural selection instead of accelerated mutational rates. We only considered genes as convergently evolved if they i) accumulated more mutations across replicate evolved lineages than *any* gene received in any randomized bootstrap trial or ii) were observed in at least half of all replicate evolved lineages. Additionally, reasoning that mutational hot spots should not be biased for or against silent mutations, the gene had to meet these criteria for the nonsynonymous mutation dataset but not also the synonymous mutation dataset. We applied these tests separately for each treatment group (e.g., pCO_2_ treatments for phytoplankton genomes, or pCO_2_ × partner for *Alteromonas* genomes). We further supported this curation step by performing a gene-by-gene linear model test in R to discover genes significantly more mutated under one treatment than another. Genes that met the curation criteria in one treatment group only and that also demonstrated statistical enrichment in that group were considered optimal candidates for genes under natural selection for a given treatment.

Genes remaining after this curation process were then analyzed for function. First, we binned genes into KEGG pathway groups using Over Representation Analysis (ORA) with the function *enrichKEGG* in *clusterProfiler* in R (*60*). The argument ‘*organism*’ was set to the KEGG organism codes ‘*pmi*’ and ‘*syn*’ for *Prochlorococcus* and *Synechocystis*, respectively, whereas it was to set to ‘ko’ (KEGG Orthology) for *Synechococcus*, *T. oceanica*, *E. huxleyi* and *Alteromonas*. KO identifiers to individual genes were assigned for each organism using the KEGG automatic annotation server website by the bi-directional best hit (BBH) method (*61*). P-values correspond to the comparison between the gene ratio (i.e., the number of genes that match that gene set divided by the number of genes in the ‘hit’ database) and the background ratio (i.e., the number of genes in the gene set divided by the total number of unique genes in the genome database). Under specific contrast comparisons for *Alteromonas* considering the mixed effect of co-cultures (i.e., MIT9312:CC9311, MIT9312:CCMP1005, MIT9312:CCM3171, MIT9312: CCMP1005:CCM3171 and all phytoplankton partners), KO identifiers were examined directly using the KEGG database website (*62*).

Second, the degree of association between mutating genes in EZ55 was analyzed using the Species Pairwise Association Analysis (SPAA) algorithm in R (*63*), reasoning that presence vs. absence of mutations in a given gene in a culture was mathematically comparable to presence vs. absence of species in an ecosystem. The mutation data was transformed into a binary presence vs. absence matrix, with genes having at least 1 mutation in each lineage coded as 1 and those without mutations coded as 0. The logistic regression function was used to calculate the odds ratio for mutations in each gene as described in (*63*), using cutoffs of 0.5 and 0.9 for the minimum and maximum mutation frequencies, respectively. SPAA estimated Spearman’s rank correlation coefficient for each gene pair, measuring the monotonic predictive relationship, i.e., the likelihood of a mutation in one gene predicting a mutation in the other. After manually examining the data, we further filtered the pairwise predictions to consider only the most positively and negatively correlated pairs (coefficients between 0.5 to 0.9 for positive relationships and −0.3 to −0.2 for negative relationships). The curated matrix was imported in Cytoscape 3.8.2 for network visualization (*64*). The network was analyzed as a directed graph to obtain the indegree and outdegree of each node, i.e., the number of other genes to which each gene is connected as a predictive target or source, respectively. The target gene of each set of genes was illustrated as an arrow pointed towards the target gene, and the gradient of the edge connecting genes was set to reflect either a positive or negative correlation.

Finally, we manually examined several genes of interest identified from these analyses using BLAST (*65*) against the NCBI nr database to attempt to provide superior annotations for hypothetical proteins or other vaguely described gene products.

### Copy number analysis

We used R scripts to extract coverage data from all breseq-assembled genomes (summarized in Table S2). Estimated copy numbers for plasmids reported in Table S2 were initially obtained simply by dividing average plasmid coverage by average chromosome coverage. However, inspection of coverage maps for the *Alteromonas* plasmid revealed highly uneven coverage (Fig. S6). Closer analysis revealed three plasmid regions with different patterns of coverage. One region was generally present at the same or greater coverage as the chromosome, one was often completely absent, and a third often existed at an intermediate coverage level. We reasoned that these differences may reflect homologous recombination events leading to the excision of parts of the plasmid and/or insertion of plasmid regions into the *Alteromonas* chromosome and therefore looked for homologous regions between the plasmid and chromosome sequences using a dot plot analysis via the D-GENIES web interface (*66*). This analysis revealed several points of homology, including one approximately 9kb sequence, thus suggesting two things. First, a reduced size plasmid likely evolved as a subpopulation in several lineages, with approximately 50kb deleted following an unknown but reproducible event. Second, an approximately 70kb section likely integrated into the chromosome of most lineages, and in several lineages became the only surviving portion of the plasmid, with no evidence remaining of subpopulations carrying the remaining 150kb of the plasmid. Because of these trends, we calculated average coverage of the plasmid at three different regions: 170kb to 180kb to estimate the “insertable” portion of the plasmid, 120kb to 130kb to estimate the full, free plasmids, and 40kb to 50kb to represent the possible reduced-size plasmid. We tested the effects of phytoplankton partner and pCO_2_ treatment on *Alteromonas* plasmid copy number using pairwise Mann-Whitney tests in R with manually calculated Holm-Bonferroni corrections for multiple tests (*67*). We also compared the likelihood of plasmid loss (defined as less than 1% coverage of plasmid region 120kb to 130kb) in *Alteromonas* paired with cyanobacteria vs. eukaryotic phytoplankton using Fisher’s exact test in R.

Examination of coverage maps further revealed the presence of elevated copy numbers for some chromosomal regions, suggesting the presence of duplications. We therefore investigated these more closely using R scripts, retrieving any regions where the average coverage was 5× the coverage (or 5× the standard deviation of coverage for low-coverage eukaryote genomes) across the chromosome. We excluded from further analysis all duplications in eukaryotic genomes where repetitive elements led to tens of thousands of qualifying duplications, as well as duplications falling outside of coding sequences or cis-acting promoter regions. After this process, only one duplication remained of interest: a promoter region mutation in the apolipoprotein N-acyltransferase gene in *Prochlorococcus* MIT9312, clearly visible in coverage maps (Fig. S1) and present in most lineages with up to 120× duplication.

