## Supplemental Material for "Marine phytoplankton and heterotrophic bacteria rapidly adapt to future pCO_2_ conditions in experimental co-cultures"

Tables S1 – S3
Tables S4 – S6 are provided as .csv files for download

Figures S1 – S23

**Table S1. Culture properties.**

| Strain | Designation | Organism | Partner | pCO2 | Medium | Transfer Density | Start Date | End Date | Transfers | Generations | Crash Restarts | Contamination Restarts | Move Restarts |
|---|---|---|---|---|---|---|---|---|---|---|---|---|---|
| LTPE191 | C-1 | Synechocystis PCC6803 | None | 400ppm | SEv | 2.60E+05 | 8/21/14 | 5/16/17 | 125 | 588 | 0 | 3 | 1 |
| LTPE192 | C-2 | Synechocystis PCC6803 | None | 400ppm | SEv | 2.60E+05 | 8/14/14 | 8/7/17 | 129 | 606 | 0 | 3 | 1 |
| LTPE193 | C-3 | Synechocystis PCC6803 | None | 400ppm | SEv | 2.60E+05 | 8/14/14 | 8/7/17 | 136 | 639 | 0 | 4 | 1 |
| LTPE194 | C-4 | Synechocystis PCC6803 | None | 400ppm | SEv | 2.60E+05 | 8/21/14 | 8/9/17 | 133 | 625 | 0 | 2 | 1 |
| LTPE195 | C-5 | Synechocystis PCC6803 | None | 400ppm | SEv | 2.60E+05 | 8/21/14 | 8/4/17 | 129 | 606 | 0 | 4 | 1 |
| LTPE196 | C+1 | Synechocystis PCC6803 | None | 800ppm | SEv | 2.60E+05 | 8/21/14 | 7/12/17 | 129 | 606 | 0 | 3 | 1 |
| LTPE197 | C+2 | Synechocystis PCC6803 | None | 800ppm | SEv | 2.60E+05 | 8/14/14 | 8/20/17 | 107 | 503 | 0 | 3 | 1 |
| LTPE198 | C+3 | Synechocystis PCC6803 | None | 800ppm | SEv | 2.60E+05 | 8/14/14 | 8/22/17 | 120 | 564 | 0 | 2 | 1 |
| LTPE199 | C+4 | Synechocystis PCC6803 | None | 800ppm | SEv | 2.60E+05 | 8/20/14 | 8/7/17 | 117 | 550 | 0 | 4 | 1 |
| LTPE200 | C+5 | Synechocystis PCC6803 | None | 800ppm | SEv | 2.60E+05 | 8/21/14 | 8/7/17 | 121 | 569 | 0 | 3 | 1 |
| LTPE397 | M-1 | Prochlorococcus MIT9312 | Alteromonas EZ55 | 400ppm | PEv | 2.60E+06 | 1/23/16 | 7/1/18 | 108 | 508 | 5 | 0 | 0 |
| LTPE398 | M-2 | Prochlorococcus MIT9312 | Alteromonas EZ55 | 400ppm | PEv | 2.60E+06 | 1/10/16 | 10/31/18 | 77 | 362 | 3 | 1 | 0 |
| LTPE399 | M-3 | Prochlorococcus MIT9312 | Alteromonas EZ55 | 400ppm | PEv | 2.60E+06 | 1/10/16 | 4/14/18 | 107 | 503 | 1 | 0 | 0 |
| LTPE400 | M-4 | Prochlorococcus MIT9312 | Alteromonas EZ55 | 400ppm | PEv | 2.60E+06 | 1/10/16 | 10/15/18 | 90 | 423 | 3 | 0 | 0 |
| LTPE401 | M-5 | Prochlorococcus MIT9312 | Alteromonas EZ55 | 400ppm | PEv | 2.60E+06 | 1/10/16 | 4/7/18 | 108 | 508 | 2 | 0 | 0 |
| LTPE402 | M-6 | Prochlorococcus MIT9312 | Alteromonas EZ55 | 400ppm | PEv | 2.60E+06 | 1/21/16 | 2/17/18 | 108 | 508 | 0 | 0 | 0 |
| LTPE403 | M+1 | Prochlorococcus MIT9312 | Alteromonas EZ55 | 800ppm | PEv | 2.60E+06 | 1/23/16 | 10/15/18 | 108 | 508 | 4 | 1 | 0 |
| LTPE404 | M+2 | Prochlorococcus MIT9312 | Alteromonas EZ55 | 800ppm | PEv | 2.60E+06 | 1/17/16 | 6/29/18 | 108 | 508 | 2 | 1 | 0 |
| LTPE405 | M+3 | Prochlorococcus MIT9312 | Alteromonas EZ55 | 800ppm | PEv | 2.60E+06 | 1/10/16 | 8/12/18 | 108 | 508 | 3 | 1 | 0 |
| LTPE406 | M+4 | Prochlorococcus MIT9312 | Alteromonas EZ55 | 800ppm | PEv | 2.60E+06 | 1/10/16 | 12/3/18 | 93 | 437 | 3 | 1 | 0 |
| LTPE407 | M+5 | Prochlorococcus MIT9312 | Alteromonas EZ55 | 800ppm | PEv | 2.60E+06 | 1/10/16 | 10/15/18 | 73 | 343 | 3 | 1 | 0 |
| LTPE408 | M+6 | Prochlorococcus MIT9312 | Alteromonas EZ55 | 800ppm | PEv | 2.60E+06 | 1/17/16 | 6/25/18 | 108 | 508 | 3 | 1 | 0 |
| LTPE421 | W-1 | Synechococcus CC9311 | Alteromonas EZ55 | 400ppm | SEv | 2.60E+05 | 6/20/16 | 6/19/18 | 107 | 503 | 3 | 0 | 0 |
| LTPE422 | W-2 | Synechococcus CC9311 | Alteromonas EZ55 | 400ppm | SEv | 2.60E+05 | 6/20/16 | 3/12/18 | 107 | 503 | 3 | 0 | 0 |
| LTPE423 | W-3 | Synechococcus CC9311 | Alteromonas EZ55 | 400ppm | SEv | 2.60E+05 | 6/20/16 | 6/3/18 | 107 | 503 | 5 | 0 | 0 |
| LTPE424 | W-4 | Synechococcus CC9311 | Alteromonas EZ55 | 400ppm | SEv | 2.60E+05 | 6/20/16 | 12/3/18 | 96 | 451 | 3 | 0 | 0 |
| LTPE425 | W-5 | Synechococcus CC9311 | Alteromonas EZ55 | 400ppm | SEv | 2.60E+05 | 6/20/16 | 6/3/18 | 107 | 503 | 3 | 0 | 0 |
| LTPE426 | W-6 | Synechococcus CC9311 | Alteromonas EZ55 | 400ppm | SEv | 2.60E+05 | 6/20/16 | 4/2/18 | 107 | 503 | 2 | 0 | 0 |
| LTPE427 | W+1 | Synechococcus CC9311 | Alteromonas EZ55 | 800ppm | SEv | 2.60E+05 | 6/28/16 | 7/13/18 | 107 | 503 | 3 | 1 | 0 |
| LTPE428 | W+2 | Synechococcus CC9311 | Alteromonas EZ55 | 800ppm | SEv | 2.60E+05 | 6/28/16 | 5/7/18 | 107 | 503 | 2 | 1 | 0 |
| LTPE429 | W+3 | Synechococcus CC9311 | Alteromonas EZ55 | 800ppm | SEv | 2.60E+05 | 6/28/16 | 5/9/18 | 107 | 503 | 1 | 1 | 0 |
| LTPE430 | W+4 | Synechococcus CC9311 | Alteromonas EZ55 | 800ppm | SEv | 2.60E+05 | 6/28/16 | 9/11/18 | 107 | 503 | 7 | 1 | 0 |
| LTPE431 | W+5 | Synechococcus CC9311 | Alteromonas EZ55 | 800ppm | SEv | 2.60E+05 | 6/28/16 | 4/19/18 | 107 | 503 | 2 | 1 | 0 |
| LTPE432 | W+6 | Synechococcus CC9311 | Alteromonas EZ55 | 800ppm | SEv | 2.60E+05 | 6/28/16 | 6/12/18 | 107 | 503 | 5 | 1 | 0 |
| LTPE445 | T-1 | Thalassiosira oceanica CCMP1005 | Alteromonas EZ55 | 400ppm | FEv | 2.60E+04 | 5/10/17 | 4/2/18 | 107 | 503 | 1 | 0 | 0 |
| LTPE446 | T-2 | Thalassiosira oceanica CCMP1005 | Alteromonas EZ55 | 400ppm | FEv | 2.60E+04 | 5/10/17 | 4/7/18 | 107 | 503 | 1 | 0 | 0 |
| LTPE447 | T-3 | Thalassiosira oceanica CCMP1005 | Alteromonas EZ55 | 400ppm | FEv | 2.60E+04 | 5/10/17 | 4/14/18 | 107 | 503 | 1 | 0 | 0 |
| LTPE448 | T-4 | Thalassiosira oceanica CCMP1005 | Alteromonas EZ55 | 400ppm | FEv | 2.60E+04 | 5/10/17 | 5/11/18 | 107 | 503 | 1 | 0 | 0 |
| LTPE449 | T-5 | Thalassiosira oceanica CCMP1005 | Alteromonas EZ55 | 400ppm | FEv | 2.60E+04 | 5/10/17 | 4/16/18 | 106 | 498 | 1 | 0 | 0 |
| LTPE450 | T-6 | Thalassiosira oceanica CCMP1005 | Alteromonas EZ55 | 400ppm | FEv | 2.60E+04 | 5/10/17 | 5/11/18 | 107 | 503 | 1 | 0 | 0 |
| LTPE451 | T+1 | Thalassiosira oceanica CCMP1005 | Alteromonas EZ55 | 800ppm | FEv | 2.60E+04 | 5/10/17 | 5/5/18 | 108 | 508 | 1 | 0 | 0 |
| LTPE452 | T+2 | Thalassiosira oceanica CCMP1005 | Alteromonas EZ55 | 800ppm | FEv | 2.60E+04 | 5/10/17 | 5/12/18 | 107 | 503 | 1 | 0 | 0 |
| LTPE453 | T+3 | Thalassiosira oceanica CCMP1005 | Alteromonas EZ55 | 800ppm | FEv | 2.60E+04 | 5/10/17 | 4/19/18 | 107 | 503 | 1 | 0 | 0 |
| LTPE454 | T+4 | Thalassiosira oceanica CCMP1005 | Alteromonas EZ55 | 800ppm | FEv | 2.60E+04 | 5/10/17 | 6/9/18 | 107 | 503 | 1 | 0 | 0 |
| LTPE455 | T+5 | Thalassiosira oceanica CCMP1005 | Alteromonas EZ55 | 800ppm | FEv | 2.60E+04 | 5/10/17 | 6/14/18 | 107 | 503 | 1 | 1 | 0 |
| LTPE456 | T+6 | Thalassiosira oceanica CCMP1005 | Alteromonas EZ55 | 800ppm | FEv | 2.60E+04 | 5/10/17 | 5/9/18 | 107 | 503 | 1 | 0 | 0 |
| LTPE469 | E-1 | Emiliania huxleyi CCMP371 | Alteromonas EZ55 | 400ppm | FEv | 2.60E+04 | 5/11/17 | 6/23/18 | 108 | 508 | 1 | 0 | 0 |
| LTPE470 | E-2 | Emiliania huxleyi CCMP371 | Alteromonas EZ55 | 400ppm | FEv | 2.60E+04 | 5/12/17 | 6/28/18 | 107 | 503 | 1 | 0 | 0 |
| LTPE471 | E-3 | Emiliania huxleyi CCMP371 | Alteromonas EZ55 | 400ppm | FEv | 2.60E+04 | 5/13/17 | 7/9/18 | 107 | 503 | 1 | 0 | 0 |
| LTPE472 | E-4 | Emiliania huxleyi CCMP371 | Alteromonas EZ55 | 400ppm | FEv | 2.60E+04 | 5/14/17 | 7/20/18 | 107 | 503 | 1 | 0 | 0 |
| LTPE473 | E-5 | Emiliania huxleyi CCMP371 | Alteromonas EZ55 | 400ppm | FEv | 2.60E+04 | 5/15/17 | 7/20/18 | 107 | 503 | 1 | 0 | 0 |
| LTPE474 | E-6 | Emiliania huxleyi CCMP371 | Alteromonas EZ55 | 400ppm | FEv | 2.60E+04 | 5/16/17 | 7/14/18 | 108 | 508 | 1 | 0 | 0 |
| LTPE475 | E+1 | Emiliania huxleyi CCMP371 | Alteromonas EZ55 | 800ppm | FEv | 2.60E+04 | 5/17/17 | 8/30/18 | 107 | 503 | 0 | 2 | 0 |
| LTPE476 | E+2 | Emiliania huxleyi CCMP371 | Alteromonas EZ55 | 800ppm | FEv | 2.60E+04 | 5/18/17 | 8/18/18 | 107 | 503 | 0 | 2 | 0 |
| LTPE477 | E+3 | Emiliania huxleyi CCMP371 | Alteromonas EZ55 | 800ppm | FEv | 2.60E+04 | 5/19/17 | 8/24/18 | 107 | 503 | 0 | 2 | 0 |
| LTPE478 | E+4 | Emiliania huxleyi CCMP371 | Alteromonas EZ55 | 800ppm | FEv | 2.60E+04 | 5/20/17 | 9/13/18 | 107 | 503 | 0 | 2 | 0 |
| LTPE479 | E+5 | Emiliania huxleyi CCMP371 | Alteromonas EZ55 | 800ppm | FEv | 2.60E+04 | 5/21/17 | 9/30/18 | 107 | 503 | 0 | 2 | 0 |
| LTPE480 | E+6 | Emiliania huxleyi CCMP371 | Alteromonas EZ55 | 800ppm | FEv | 2.60E+04 | 5/22/17 | 9/8/18 | 107 | 503 | 0 | 2 | 0 |

Table S2. Genomic re-sequencing coverage.

| Culture | Phytoplankton Coverage[1] | EZ55 Chromosome Coverage | EZ55 Plasmid Coverage[2] | pSYSA coverage | pSYSG coverage | pSYSM Coverage | pSYSX coverage |
|---|---|---|---|---|---|---|---|
| PCC6803 | 102 +/- 20.9 | n/a | n/a | 111 +/- 14.1 | 111 +/- 15.4 | 146 +/- 16.3 | 142 +/- 14.1 |
| MIT9312 | 253 +/- 86.6 | 112 +/- 19.8 | 72.2 +/- 22.6 | n/a | n/a | n/a | n/a |
| CC9311 | 123 +/- 52.4 | 613 +/- 194 | 271 +/- 121 | n/a | n/a | n/a | n/a |
| CCMP371 | 20.9 +/- 4.16 | 491 +/- 255 | 426.5 +/- 250 | n/a | n/a | n/a | n/a |
| CCMP1005 | 32.5 +/- 6.37 | 567 +/- 215 | 293 +/- 89.0 | n/a | n/a | n/a | n/a |

[1] Coverage values are given as means plus/minus 95% confidence intervals of replicate lineages.
[2] EZ55 plasmid coverage is given as the average estimate across the entire ancestral plasmid. See text for descriptions of various plasmid variants that may have evolved.

Table S3. Mutational multiplicity.

| Species | Partner | Mutation Type | Mutations observed[1] | Maximum observed mutations[2] | Maximum dummy mutations[3] |
|---|---|---|---|---|---|
| MIT9312 | EZ55 | Synonymous | 104 | 8 | 4 |
| | | Nonsynonymous | 389 | 20 | 6 |
| CC9311 | EZ55 | Synonymous | 802 | 98 | 9 |
| | | Nonsynonymous | 1,382 | 36 | 11 |
| CCMP1005 | EZ55 | Synonymous | 1,475,350 | 3,297 | 864 |
| | | Nonsynonymous | 1,648,927 | 2,603 | 975 |
| CCMP371 | EZ55 | Synonymous | 536,313 | 13,634 | 954 |
| | | Nonsynonymous | 570,103 | 9,028 | 996 |
| PCC6803 | None | Synonymous | 1,689 | 226 | 14 |
| | | Nonsynonymous | 1,358 | 87 | 11 |
| EZ55 | MIT9312 | Synonymous | 1,875 | 93 | 13 |
| | | Nonsynonymous | 1,638 | 31 | 14 |
| EZ55 | CC9311 | Synonymous | 463 | 48 | 7 |
| | | Nonsynonymous | 1,427 | 43 | 12 |
| EZ55 | CCMP1005 | Synonymous | 1,665 | 22 | 13 |
| | | Nonsynonymous | 8,499 | 69 | 48 |
| EZ55 | CCMP371 | Synonymous | 2,573 | 88 | 21 |
| | | Nonsynonymous | 9,618 | 66 | 52 |

[1] Sum of all unique mutations observed across replicate lineages.
[2] Maximum number of mutations observed in a single coding sequence across all replicate lineages
[3] Maximum number of mutations observed in a single coding sequence in any of 100 bootstrapped monte carlo mutational distributions (see methods)

Tables S4, S5, and S6 are provided as .csv files available for download.

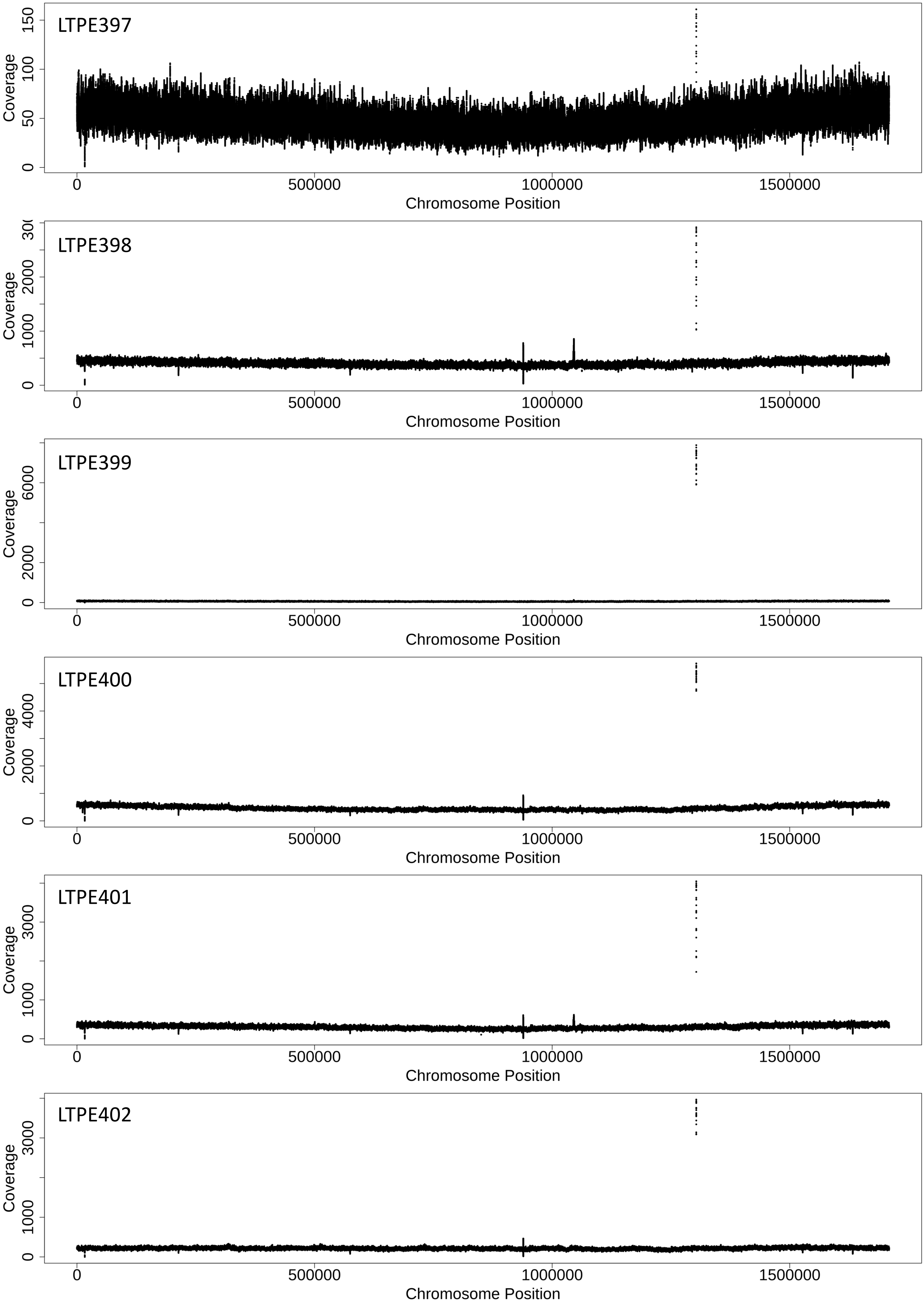
LTPE397
Coverage
0
50
100
150
0
500000
1000000
1500000
Chromosome Position
LTPE398
Coverage
0
1000
2000
LTPE399
Coverage
0
2000
6000
LTPE400
Coverage
0
2000
4000
LTPE401
Coverage
0
1000
3000
LTPE402
Coverage
0
1000
3000
Chromosome Position

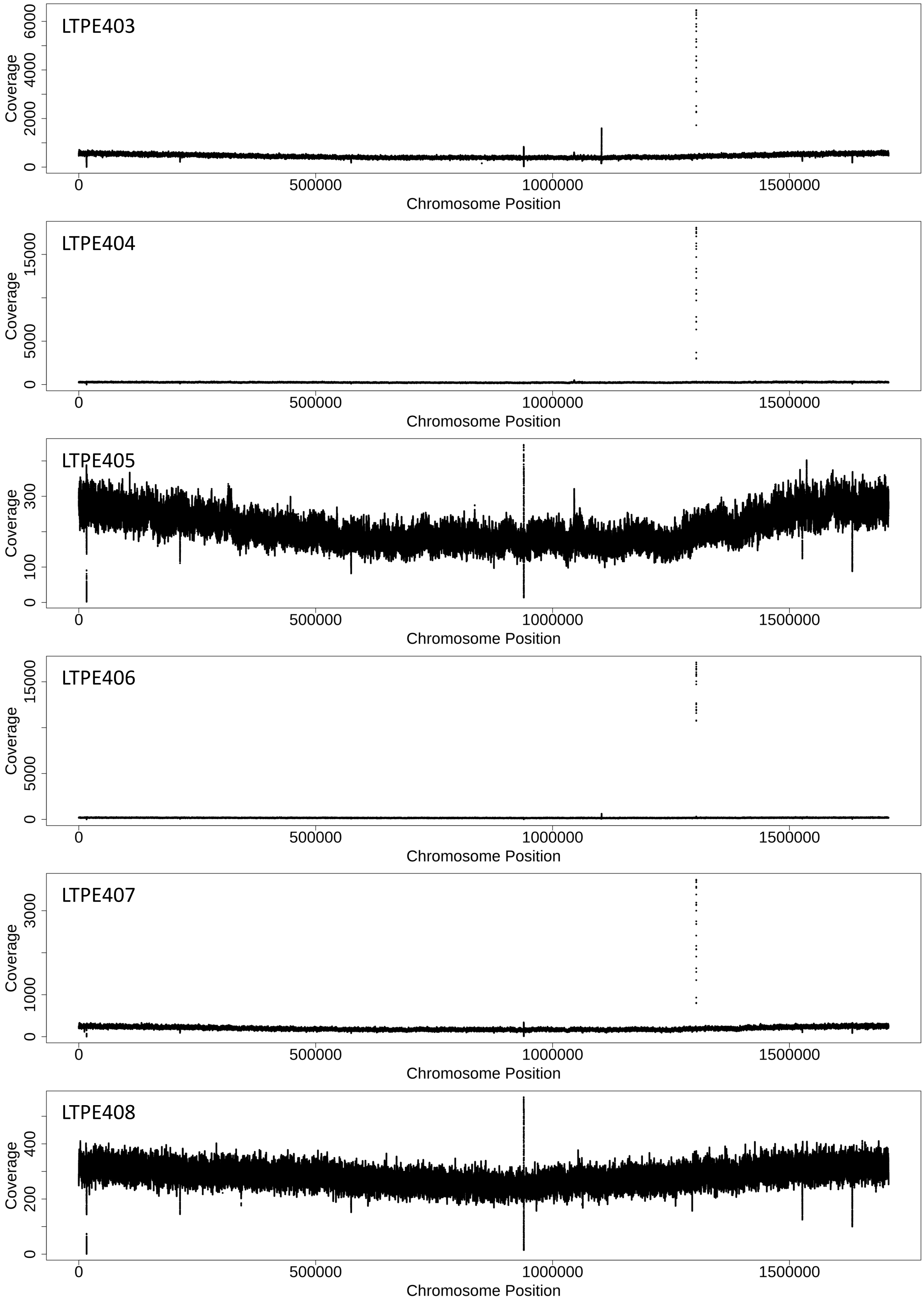
LTPE403
Coverage
0
2000
4000
6000
0
500000
1000000
1500000
Chromosome Position
LTPE404
Coverage
0
5000
15000
0
500000
1000000
1500000
Chromosome Position
LTPE405
Coverage
0
100
300
0
500000
1000000
1500000
Chromosome Position
LTPE406
Coverage
0
5000
15000
0
500000
1000000
1500000
Chromosome Position
LTPE407
Coverage
0
1000
3000
0
500000
1000000
1500000
Chromosome Position
LTPE408
Coverage
0
200
400
0
500000
1000000
1500000
Chromosome Position

**Figure S1. Sequencing coverage for evolved *Prochlorococcus* MIT9312 genomes.** Strains LTPE397 to LTPE402 were evolved at 400 ppm $pCO_2$; strains LTPE403 to LTPE408 were evolved at 800 ppm $pCO_2$. The spikes in coverage in most genomes around 1.3MB correspond to AT duplications in the promoter/5' coding sequence of the apolipoprotein N-acyltransferase gene described in the text.

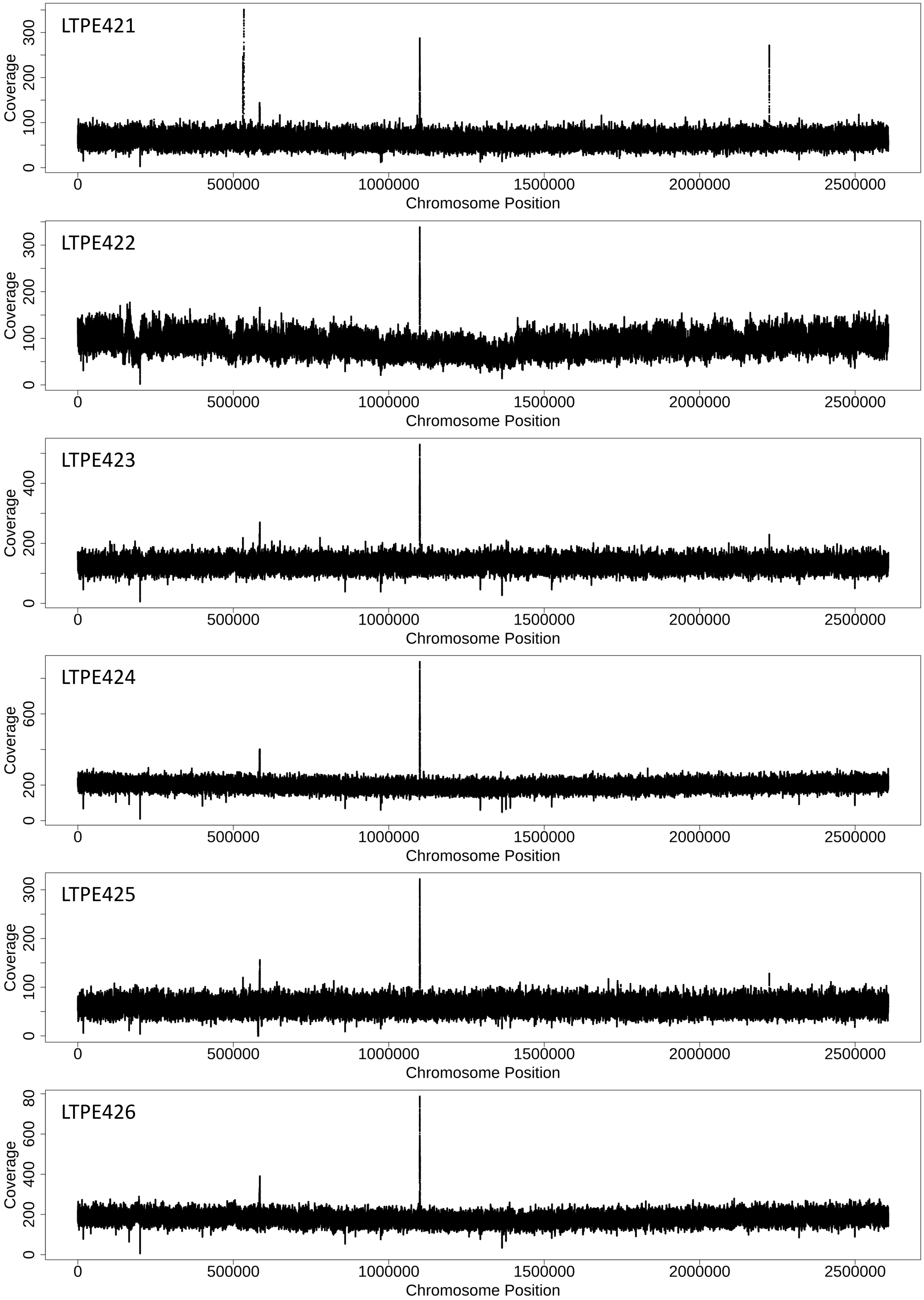
LTPE421
Coverage
0
100
200
300
0
500000
1000000
1500000
2000000
2500000
Chromosome Position
LTPE422
Coverage
0
100
200
300
0
500000
1000000
1500000
2000000
2500000
Chromosome Position
LTPE423
Coverage
0
200
400
0
500000
1000000
1500000
2000000
2500000
Chromosome Position
LTPE424
Coverage
0
200
600
0
500000
1000000
1500000
2000000
2500000
Chromosome Position
LTPE425
Coverage
0
100
200
300
0
500000
1000000
1500000
2000000
2500000
Chromosome Position
LTPE426
Coverage
0
200
400
600
80
0
500000
1000000
1500000
2000000
2500000
Chromosome Position

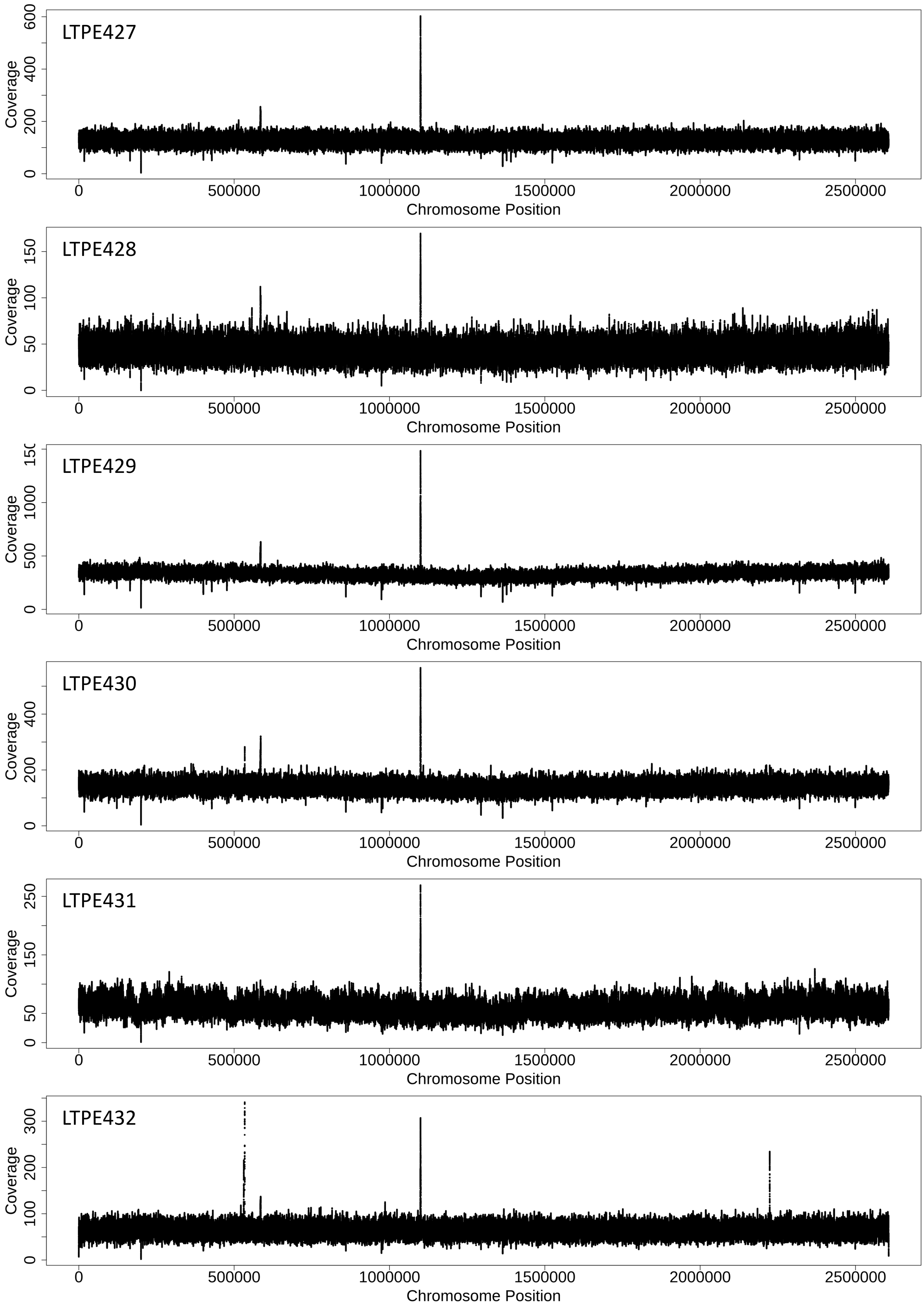
LTPE427
LTPE428
LTPE429
LTPE430
LTPE431
LTPE432
Coverage
Chromosome Position
0
500000
1000000
1500000
2000000
2500000

**Figure S2. Sequencing coverage for evolved *Synechococcus* CC9311 genomes.** Strains LTPE421 to LTPE426 were evolved at 400 ppm $pCO_2$; strains LTPE427 to LTPE432 were evolved at 800 ppm $pCO_2$.

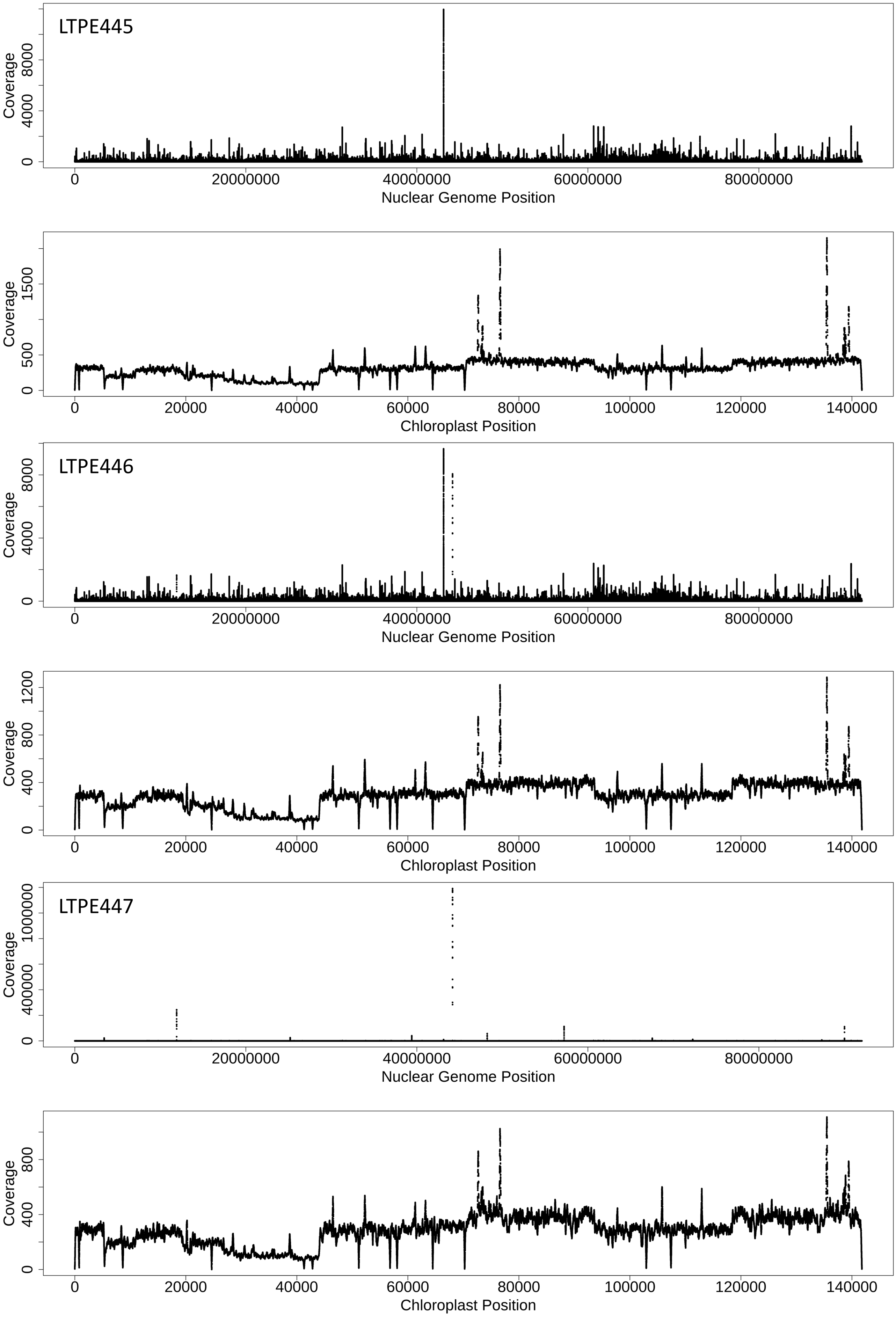

LTPE445
Coverage
0
4000
8000
0
20000000
40000000
60000000
80000000
Nuclear Genome Position
Coverage
0
500
1500
0
20000
40000
60000
80000
100000
120000
140000
Chloroplast Position
LTPE446
Coverage
0
4000
8000
0
20000000
40000000
60000000
80000000
Nuclear Genome Position
Coverage
0
400
800
1200
0
20000
40000
60000
80000
100000
120000
140000
Chloroplast Position
LTPE447
Coverage
0
400000
1000000
0
20000000
40000000
60000000
80000000
Nuclear Genome Position
Coverage
0
400
800
0
20000
40000
60000
80000
100000
120000
140000
Chloroplast Position

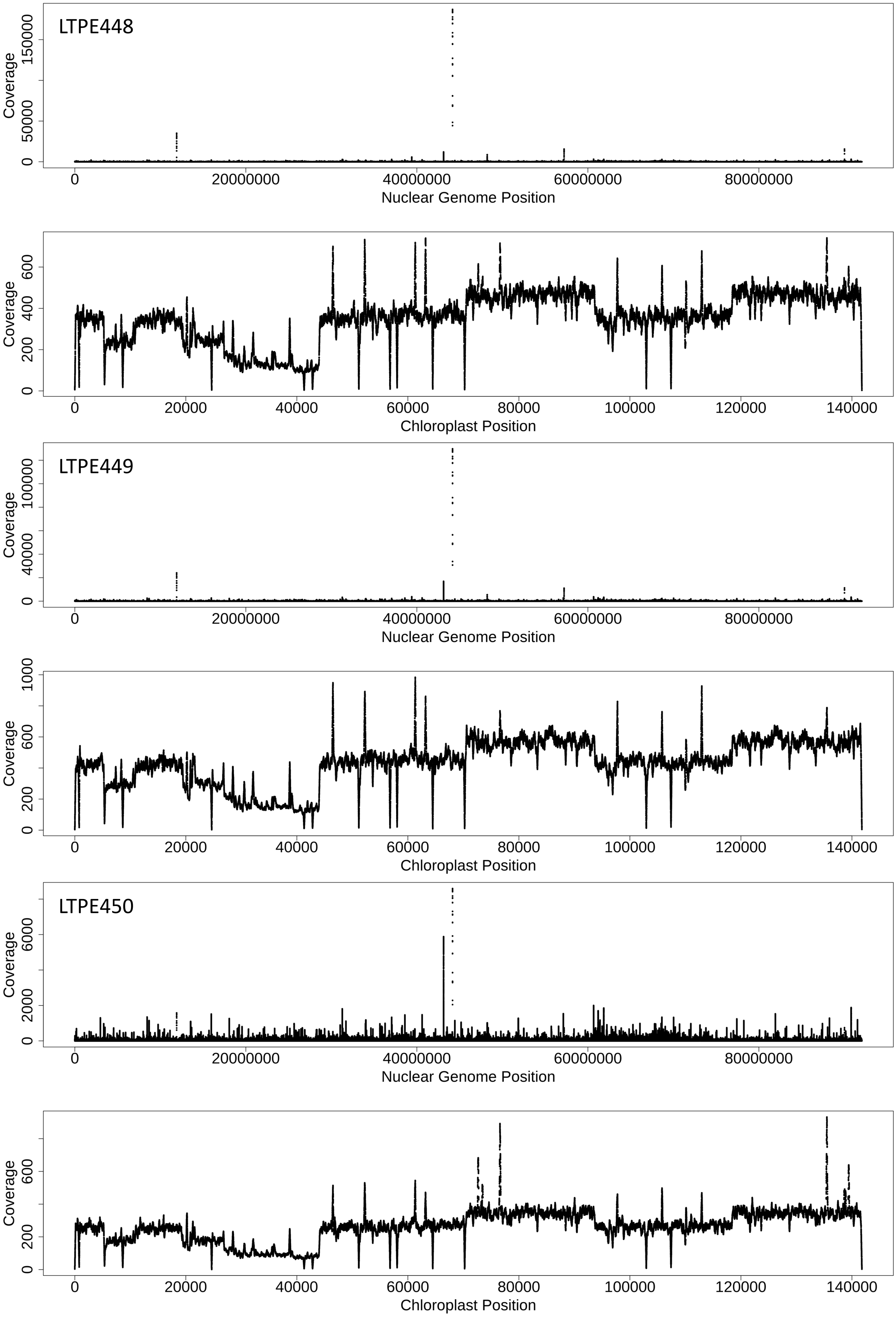

LTPE448
Coverage
0
50000
150000
0
20000000
40000000
60000000
80000000
Nuclear Genome Position
Coverage
0
200
400
600
0
20000
40000
60000
80000
100000
120000
140000
Chloroplast Position
LTPE449
Coverage
0
40000
100000
0
20000000
40000000
60000000
80000000
Nuclear Genome Position
Coverage
0
200
600
1000
0
20000
40000
60000
80000
100000
120000
140000
Chloroplast Position
LTPE450
Coverage
0
2000
6000
0
20000000
40000000
60000000
80000000
Nuclear Genome Position
Coverage
0
200
600
0
20000
40000
60000
80000
100000
120000
140000
Chloroplast Position

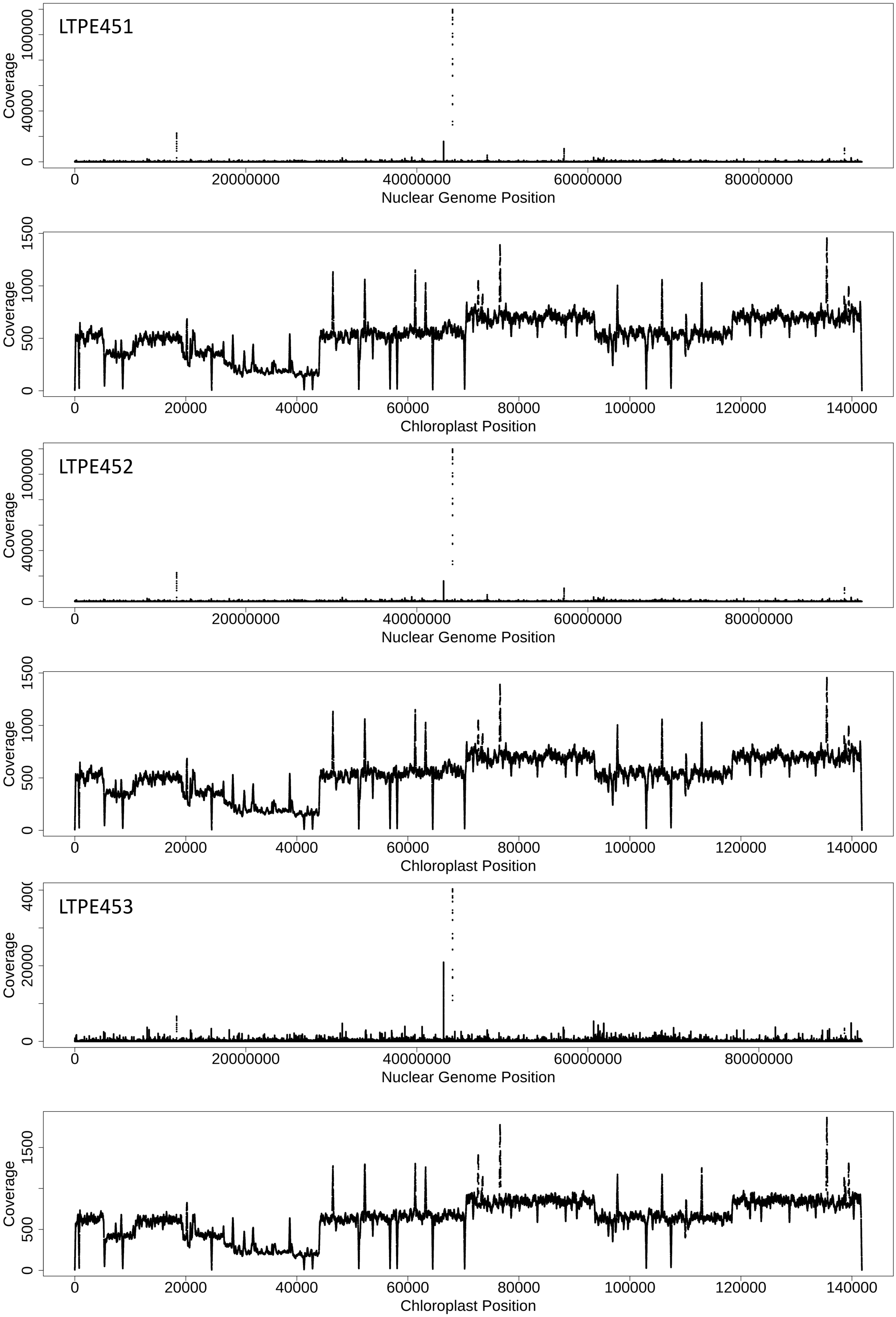
LTPE451
Coverage
0
40000
100000
0
20000000
40000000
60000000
80000000
Nuclear Genome Position
Coverage
0
500
1000
1500
0
20000
40000
60000
80000
100000
120000
140000
Chloroplast Position
LTPE452
Coverage
0
40000
100000
0
20000000
40000000
60000000
80000000
Nuclear Genome Position
Coverage
0
500
1000
1500
0
20000
40000
60000
80000
100000
120000
140000
Chloroplast Position
LTPE453
Coverage
0
20000
0
20000000
40000000
60000000
80000000
Nuclear Genome Position
Coverage
0
500
1500
0
20000
40000
60000
80000
100000
120000
140000
Chloroplast Position

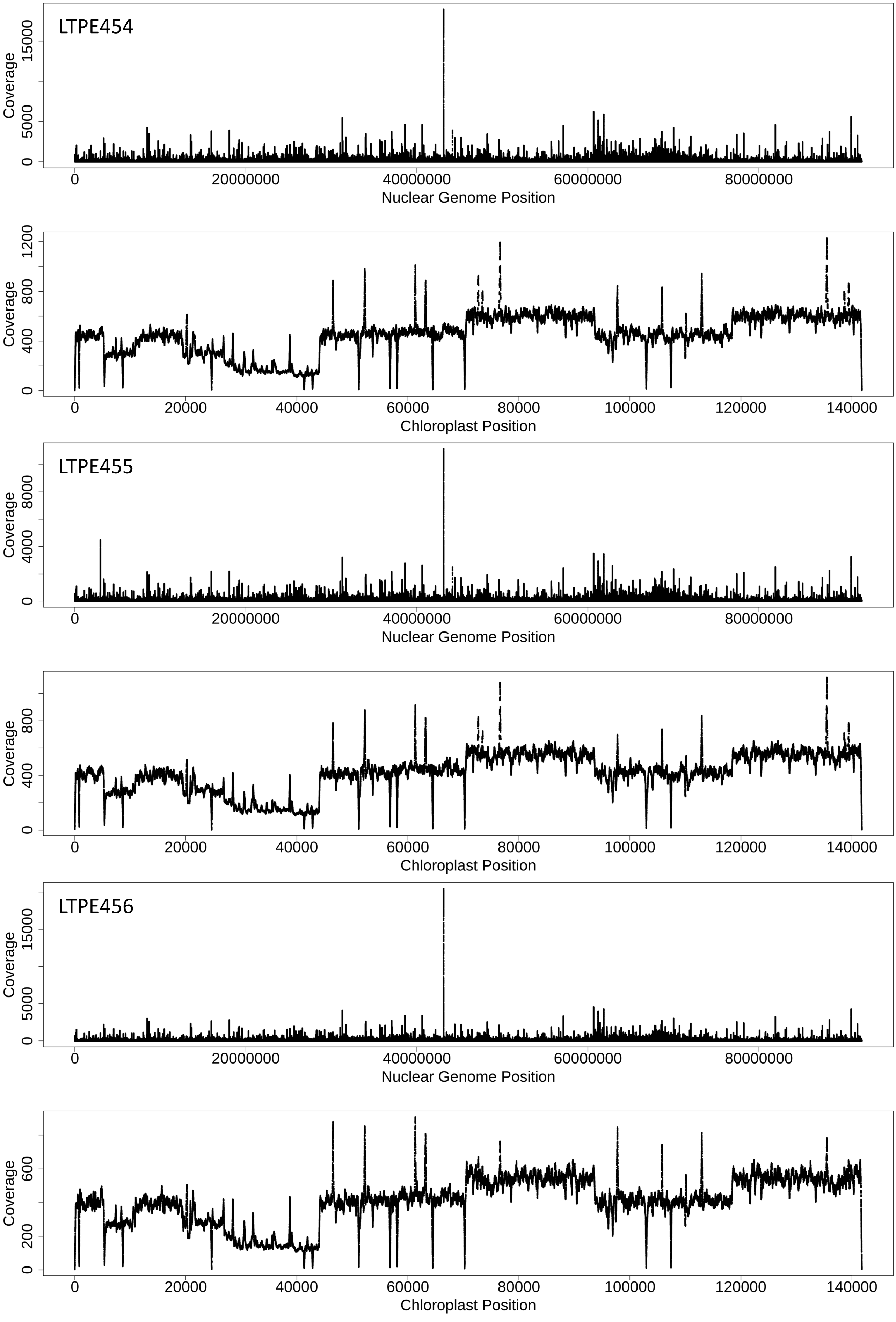
LTPE454
Coverage
0
5000
15000
0
20000000
40000000
60000000
80000000
Nuclear Genome Position
Coverage
0
400
800
1200
0
20000
40000
60000
80000
100000
120000
140000
Chloroplast Position
LTPE455
Coverage
0
4000
8000
0
20000000
40000000
60000000
80000000
Nuclear Genome Position
Coverage
0
400
800
0
20000
40000
60000
80000
100000
120000
140000
Chloroplast Position
LTPE456
Coverage
0
5000
15000
0
20000000
40000000
60000000
80000000
Nuclear Genome Position
Coverage
0
200
600
0
20000
40000
60000
80000
100000
120000
140000
Chloroplast Position

**Figure S3. Sequencing coverage for evolved *Thalassiosira oceanica* CCMP1005 genomes.** Strains LTPE445 to LTPE450 were evolved at 400 ppm $pCO_2$; strains LTPE451 to LTPE456 were evolved at 800 ppm $pCO_2$. For each strain, the nuclear genome and chloroplast genome are shown separately. The CCMP1005 reference genome was a collection of contigs, all of which were concatenated linearly in order to create this plot.

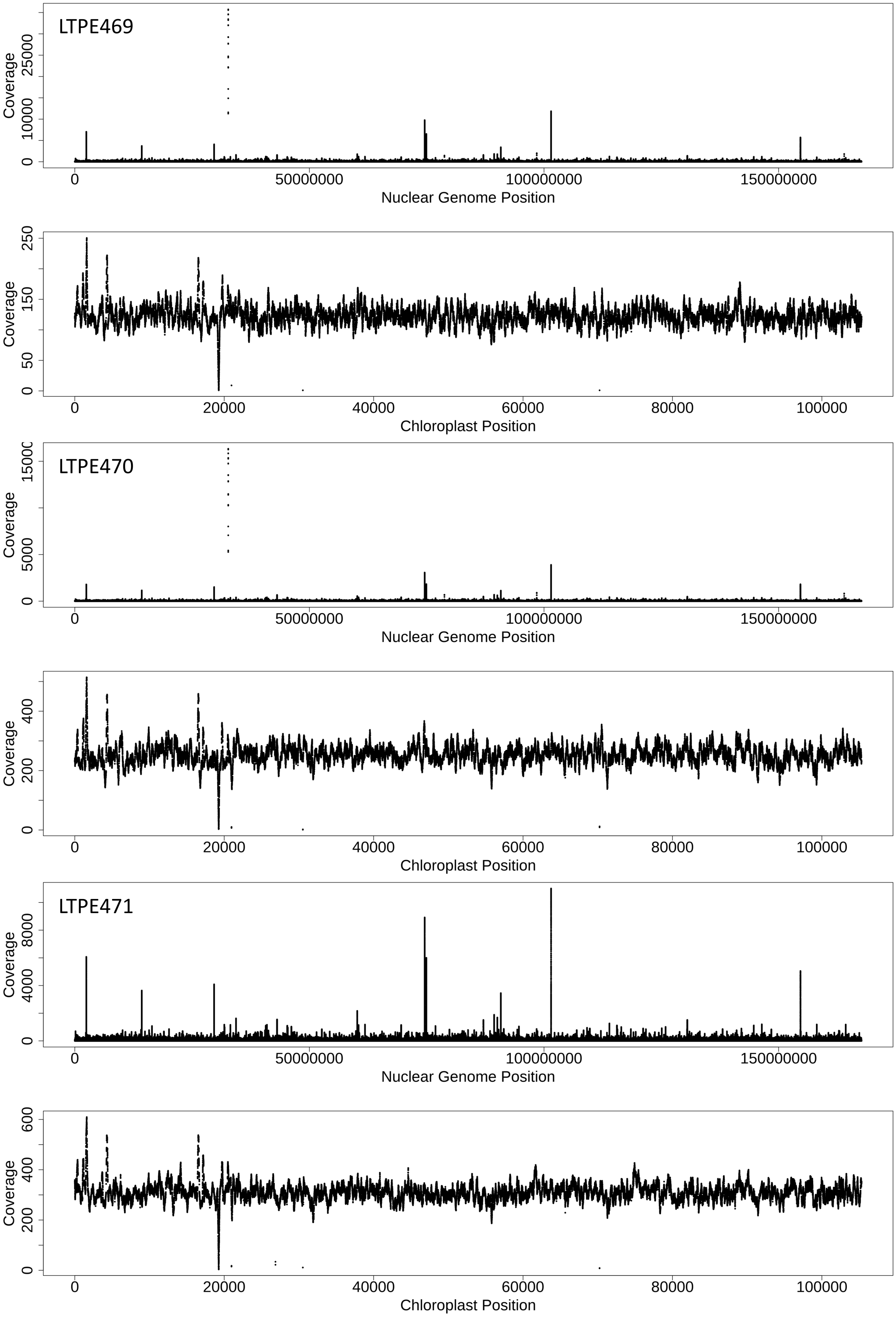
LTPE469
Coverage
0
10000
25000
0
50000000
100000000
150000000
Nuclear Genome Position
Coverage
0
50
150
250
0
20000
40000
60000
80000
100000
Chloroplast Position
LTPE470
Coverage
0
5000
15000
0
50000000
100000000
150000000
Nuclear Genome Position
Coverage
0
200
400
0
20000
40000
60000
80000
100000
Chloroplast Position
LTPE471
Coverage
0
4000
8000
0
50000000
100000000
150000000
Nuclear Genome Position
Coverage
0
200
400
600
0
20000
40000
60000
80000
100000
Chloroplast Position

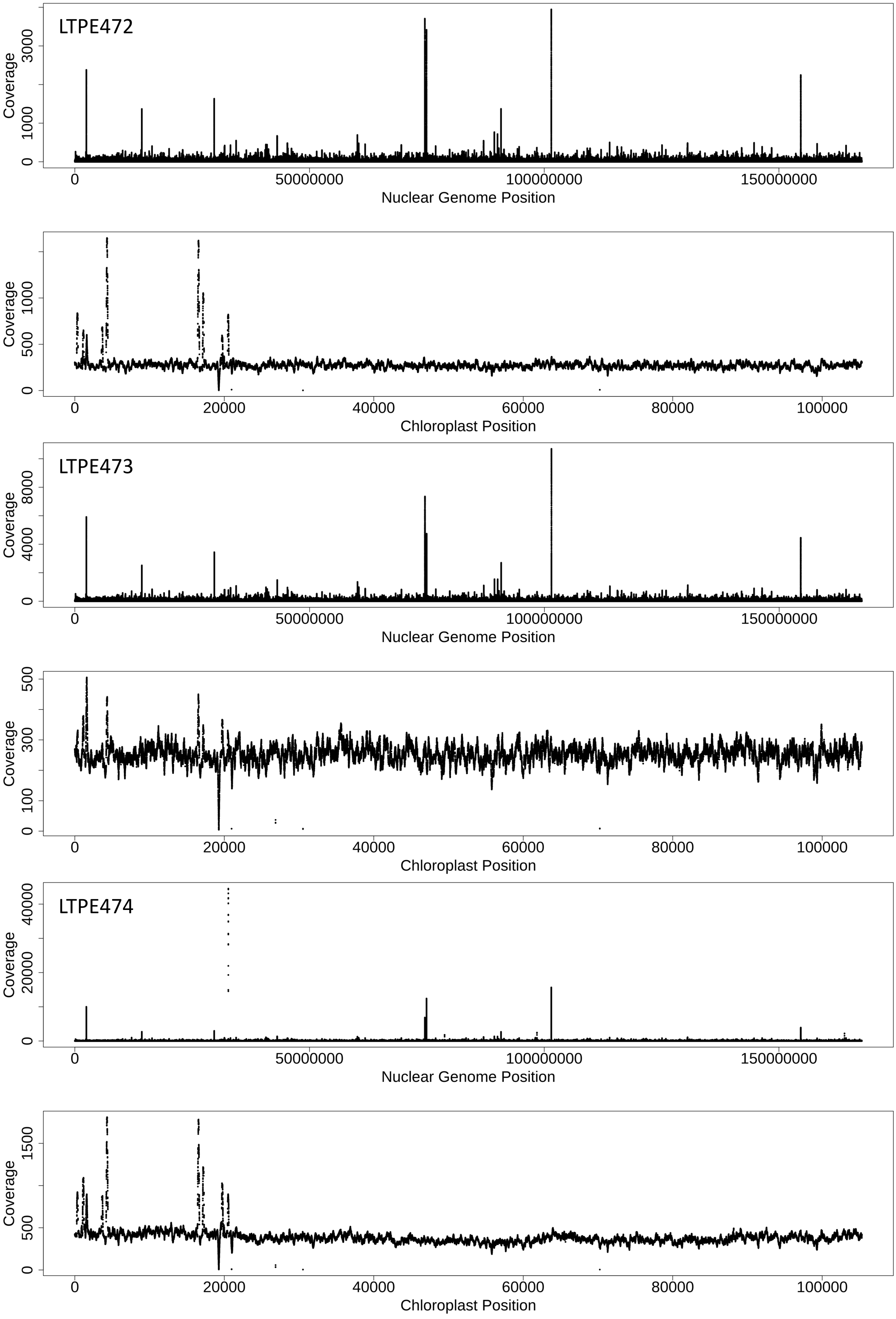

LTPE472
Coverage
0
1000
3000
0
50000000
100000000
150000000
Nuclear Genome Position
Coverage
0
500
1000
0
20000
40000
60000
80000
100000
Chloroplast Position
LTPE473
Coverage
0
4000
8000
0
50000000
100000000
150000000
Nuclear Genome Position
Coverage
0
100
300
500
0
20000
40000
60000
80000
100000
Chloroplast Position
LTPE474
Coverage
0
20000
40000
0
50000000
100000000
150000000
Nuclear Genome Position
Coverage
0
500
1500
0
20000
40000
60000
80000
100000
Chloroplast Position

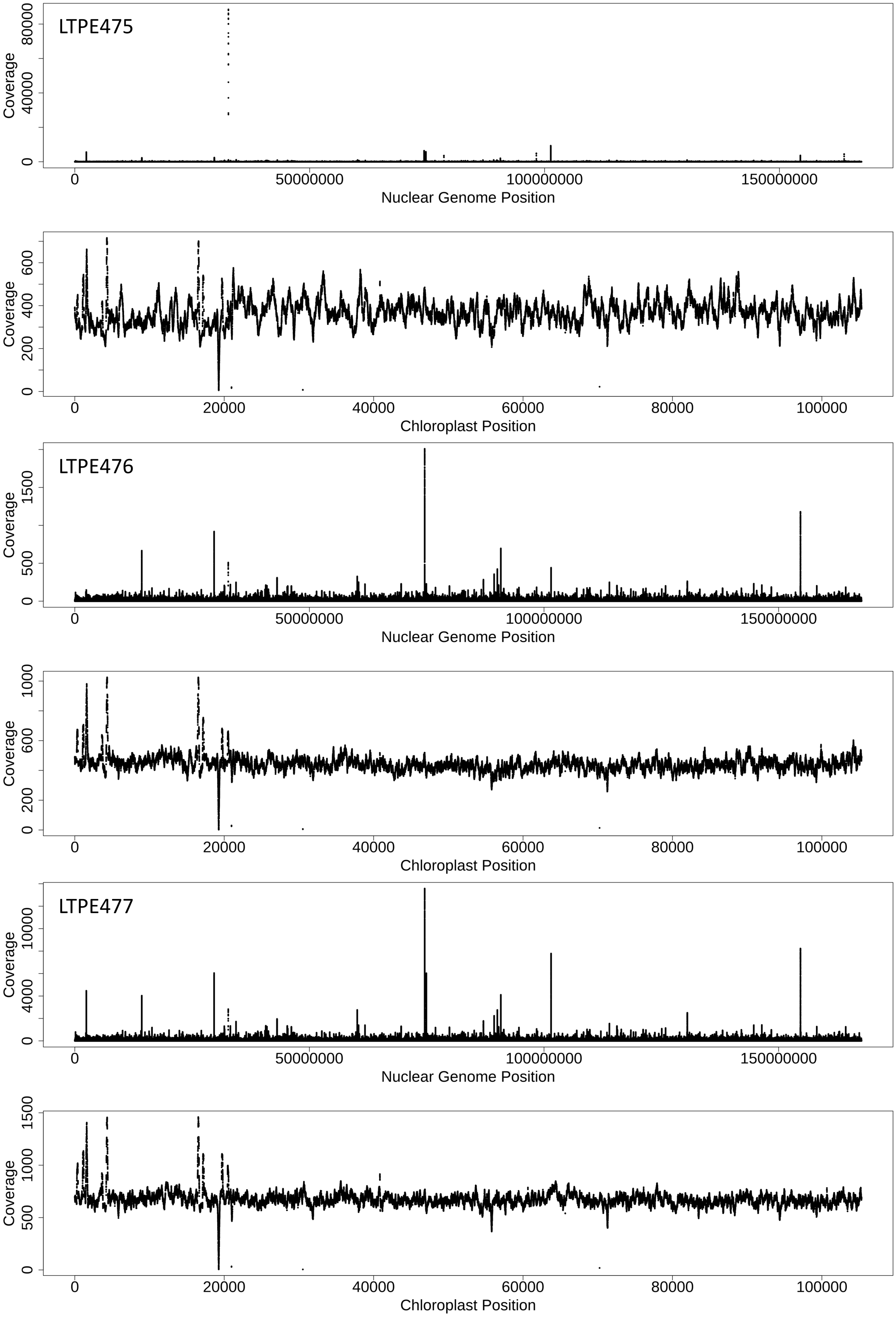
LTPE475
Coverage
0
40000
80000
0
50000000
100000000
150000000
Nuclear Genome Position
Coverage
0
200
400
600
0
20000
40000
60000
80000
100000
Chloroplast Position
LTPE476
Coverage
0
500
1500
0
50000000
100000000
150000000
Nuclear Genome Position
Coverage
0
200
600
1000
0
20000
40000
60000
80000
100000
Chloroplast Position
LTPE477
Coverage
0
4000
10000
0
50000000
100000000
150000000
Nuclear Genome Position
Coverage
0
500
1000
1500
0
20000
40000
60000
80000
100000
Chloroplast Position

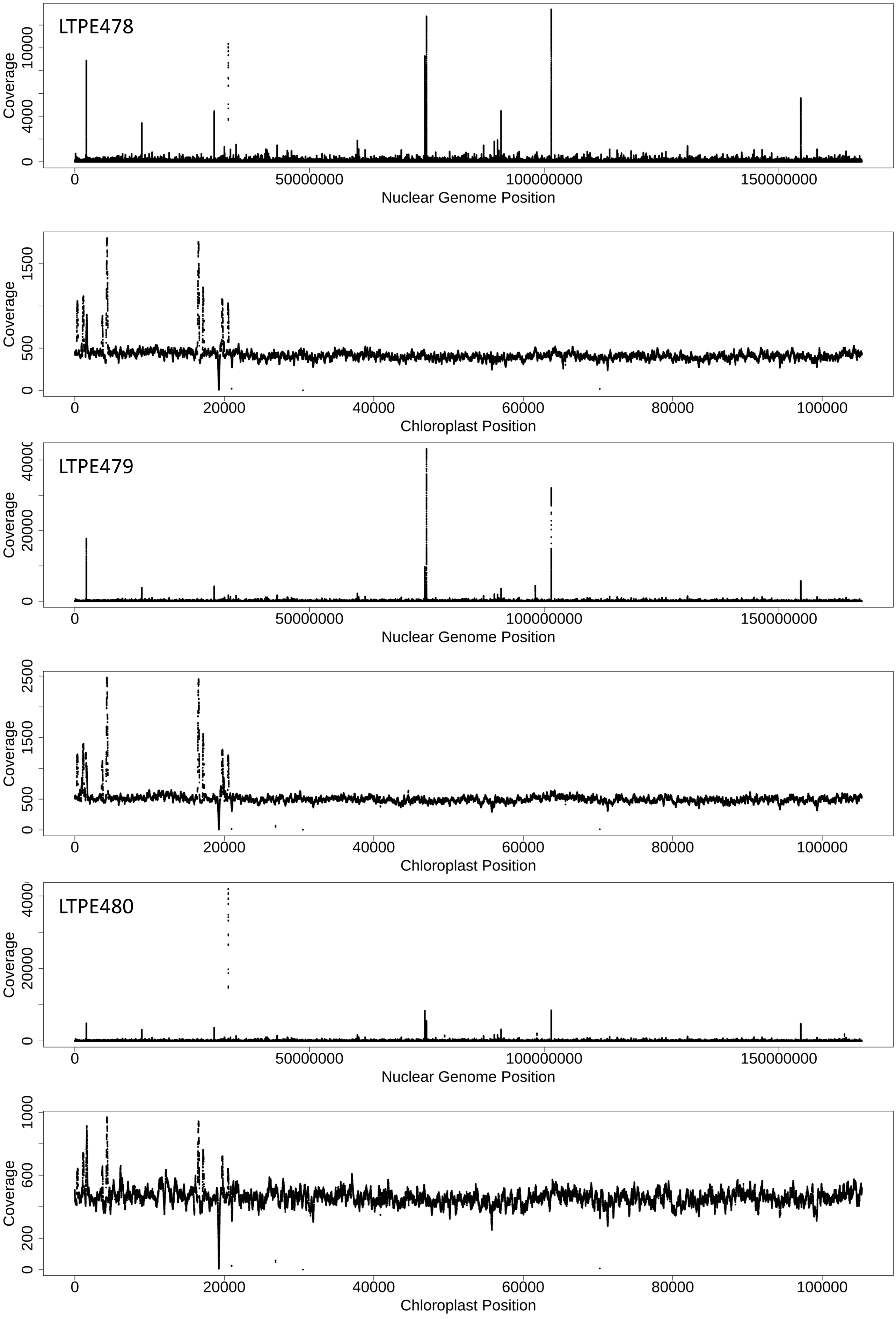
LTPE478
Coverage
0
4000
10000
0
50000000
100000000
150000000
Nuclear Genome Position
Coverage
0
500
1500
0
20000
40000
60000
80000
100000
Chloroplast Position
LTPE479
Coverage
0
20000
0
50000000
100000000
150000000
Nuclear Genome Position
Coverage
0
500
1500
2500
0
20000
40000
60000
80000
100000
Chloroplast Position
LTPE480
Coverage
0
20000
0
50000000
100000000
150000000
Nuclear Genome Position
Coverage
0
200
600
1000
0
20000
40000
60000
80000
100000
Chloroplast Position

**Figure S4. Sequencing coverage for evolved *Emiliania huxleyi* CCMP371 genomes.** Strains LTPE469 to LTPE474 were evolved at 400 ppm $pCO_2$; strains LTPE475 to LTPE480 were evolved at 800 ppm $pCO_2$. For each strain, the nuclear genome and chloroplast genome are shown separately. The CCMP371 reference genome was a collection of contigs, all of which were concatenated linearly in order to create this plot.

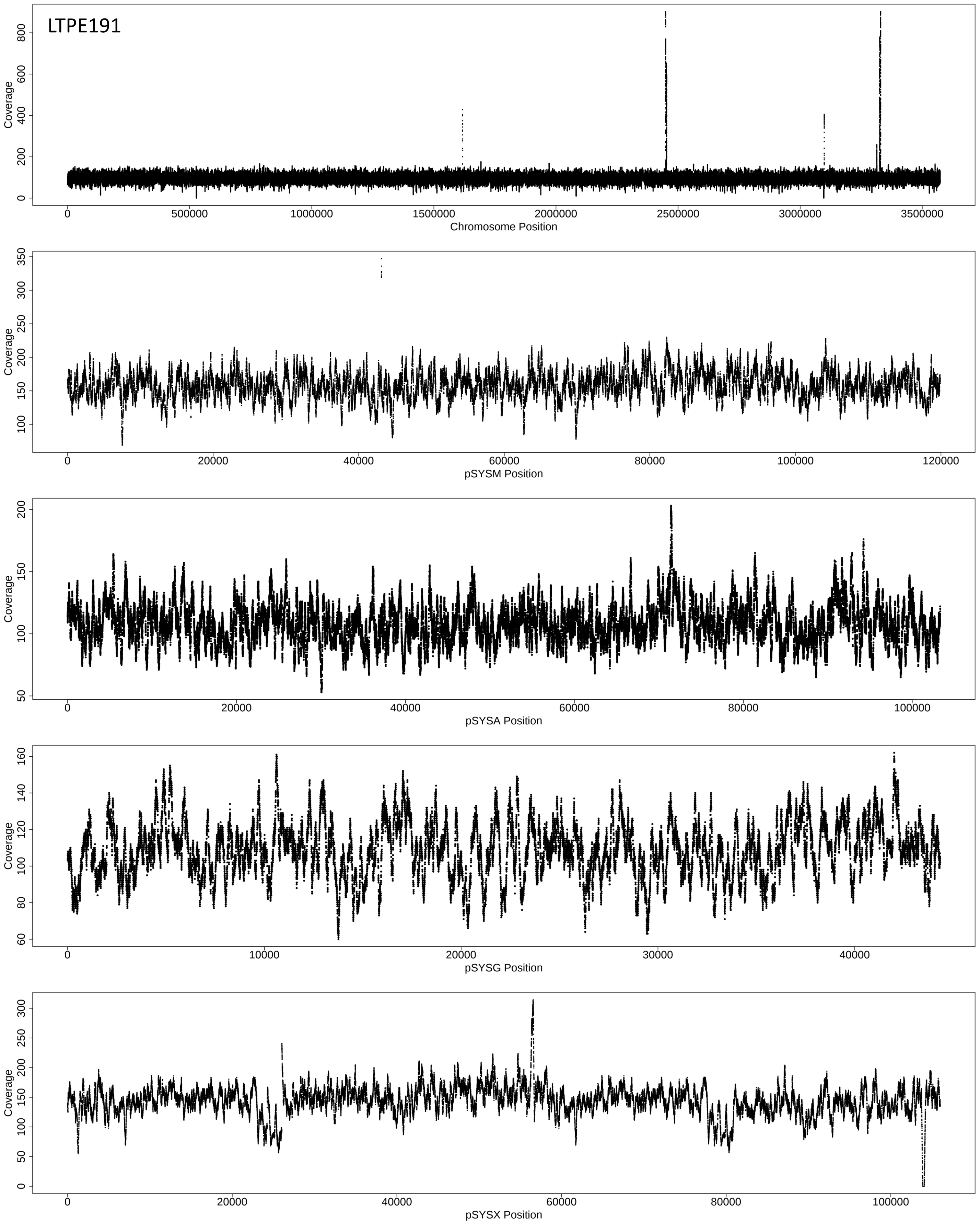
LTPE191
Coverage
0 200 400 600 800
0 500000 1000000 1500000 2000000 2500000 3000000 3500000
Chromosome Position
Coverage
100 150 200 250 300 350
0 20000 40000 60000 80000 100000 120000
pSYSM Position
Coverage
50 100 150 200
0 20000 40000 60000 80000 100000
pSYSA Position
Coverage
60 80 100 120 140 160
0 10000 20000 30000 40000
pSYSG Position
Coverage
0 50 100 150 200 250 300
0 20000 40000 60000 80000 100000
pSYSX Position

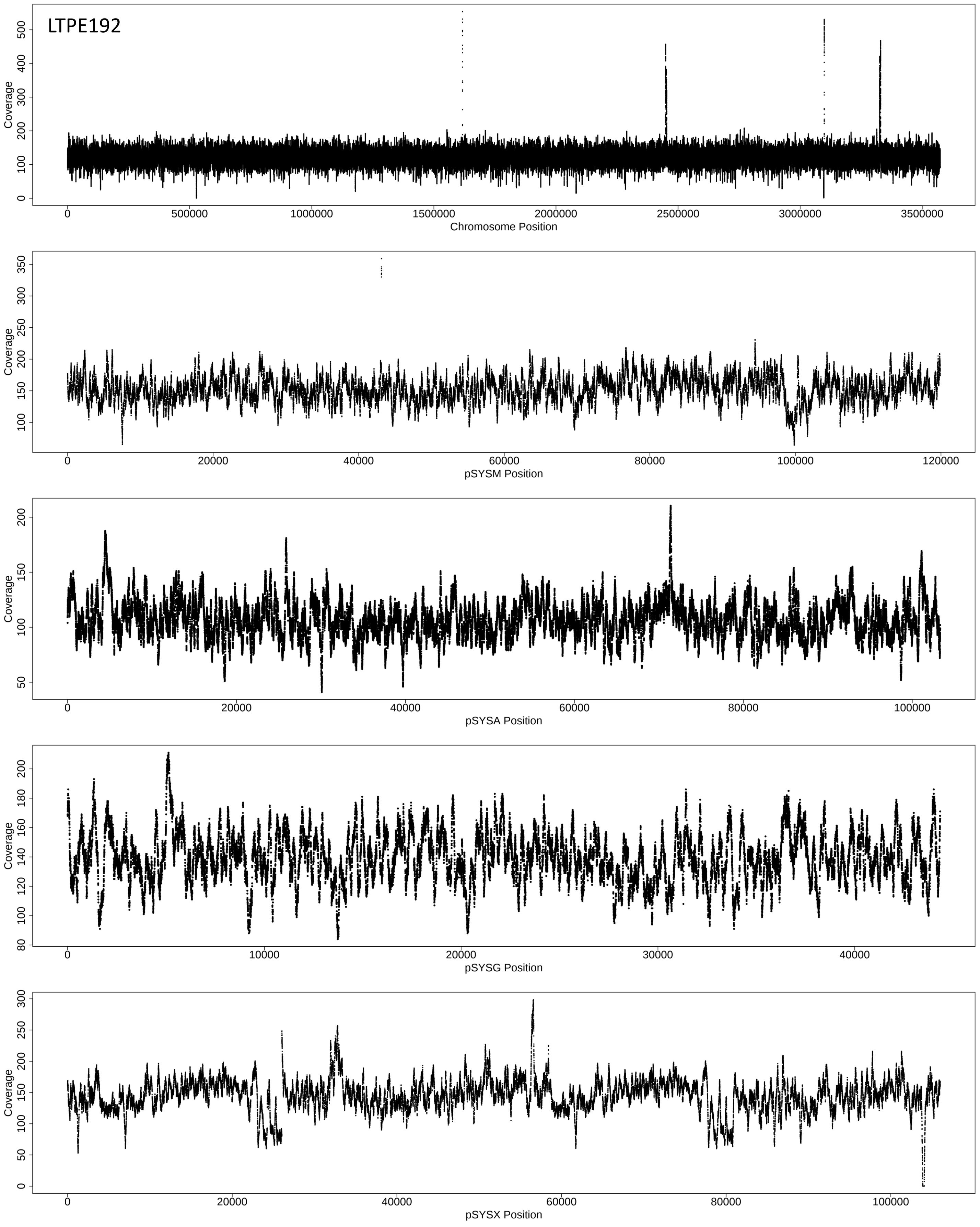
LTPE192
Coverage
0 100 200 300 400 500
0 500000 1000000 1500000 2000000 2500000 3000000 3500000
Chromosome Position
Coverage
100 150 200 250 300 350
0 20000 40000 60000 80000 100000 120000
pSYSM Position
Coverage
50 100 150 200
0 20000 40000 60000 80000 100000
pSYSA Position
Coverage
80 100 120 140 160 180 200
0 10000 20000 30000 40000
pSYSG Position
Coverage
0 50 100 150 200 250 300
0 20000 40000 60000 80000 100000
pSYSX Position

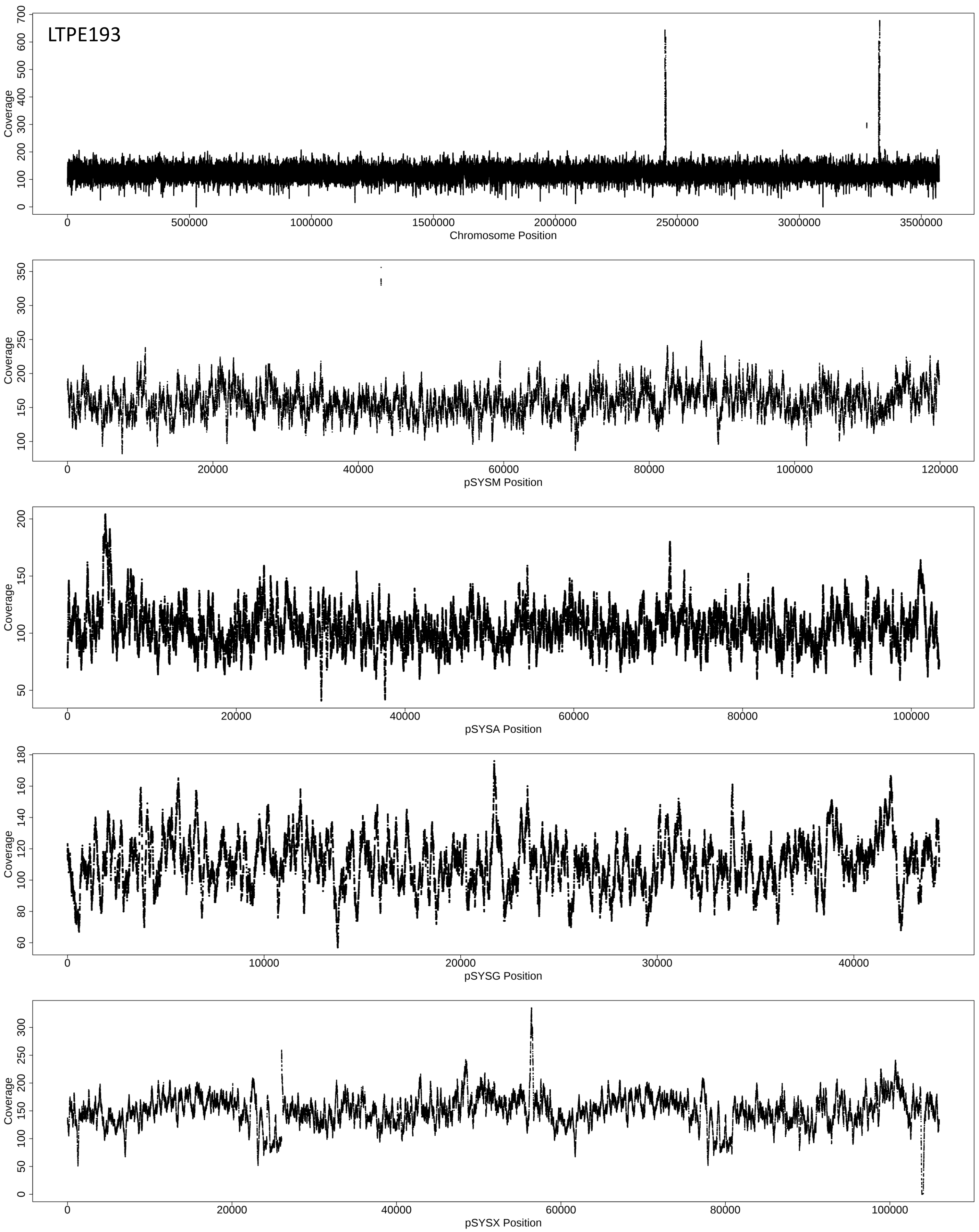
LTPE193
Coverage
0
100
200
300
400
500
600
700
0
500000
1000000
1500000
2000000
2500000
3000000
3500000
Chromosome Position
Coverage
100
150
200
250
300
350
0
20000
40000
60000
80000
100000
120000
pSYSM Position
Coverage
50
100
150
200
0
20000
40000
60000
80000
100000
pSYSA Position
Coverage
60
80
100
120
140
160
180
0
10000
20000
30000
40000
pSYSG Position
Coverage
0
50
100
150
200
250
300
0
20000
40000
60000
80000
100000
pSYSX Position

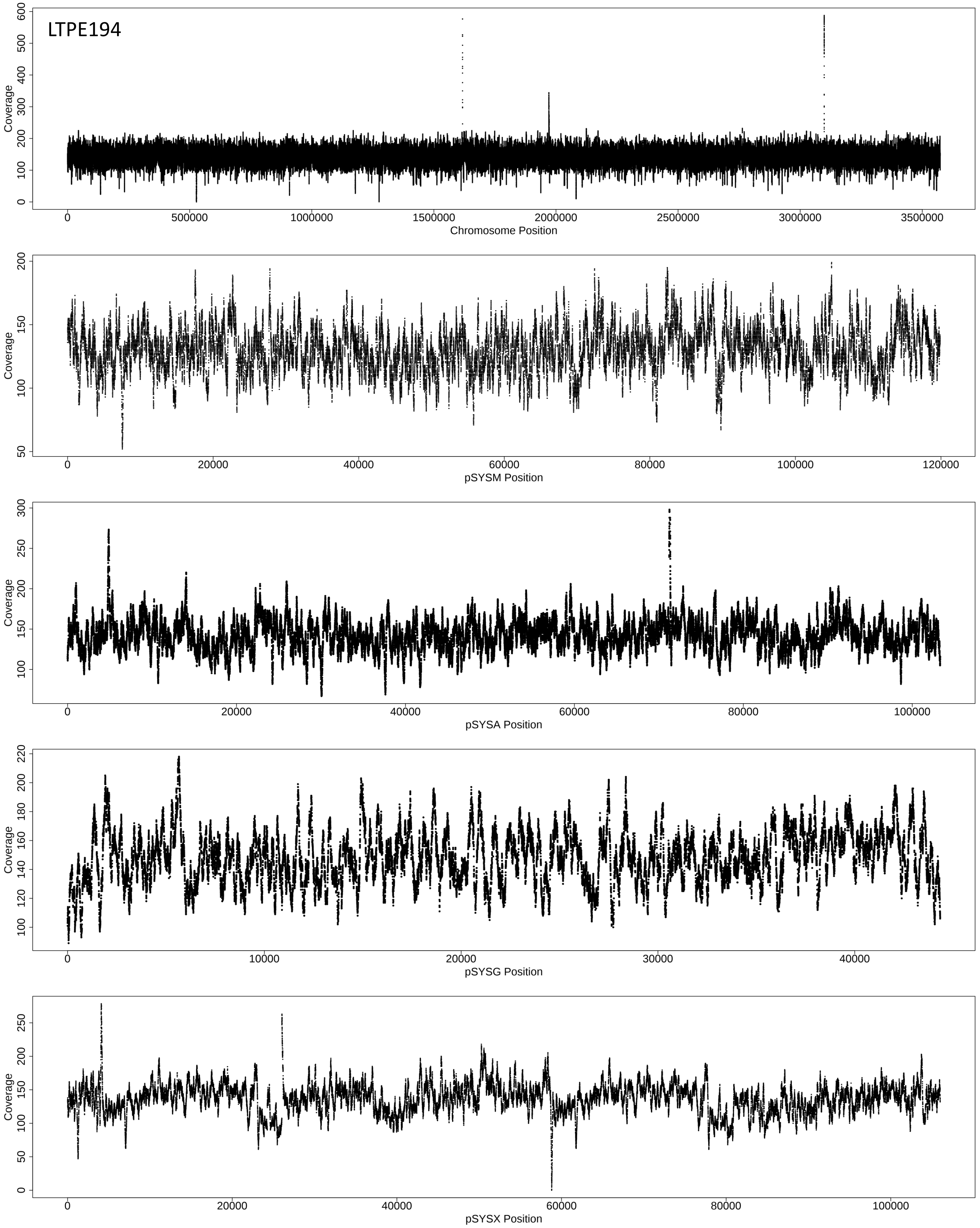
LTPE194
Coverage
0
100
200
300
400
500
600
0
500000
1000000
1500000
2000000
2500000
3000000
3500000
Chromosome Position
Coverage
50
100
150
200
0
20000
40000
60000
80000
100000
120000
pSYSM Position
Coverage
100
150
200
250
300
0
20000
40000
60000
80000
100000
pSYSA Position
Coverage
100
120
140
160
180
200
220
0
10000
20000
30000
40000
pSYSG Position
Coverage
0
50
100
150
200
250
0
20000
40000
60000
80000
100000
pSYSX Position

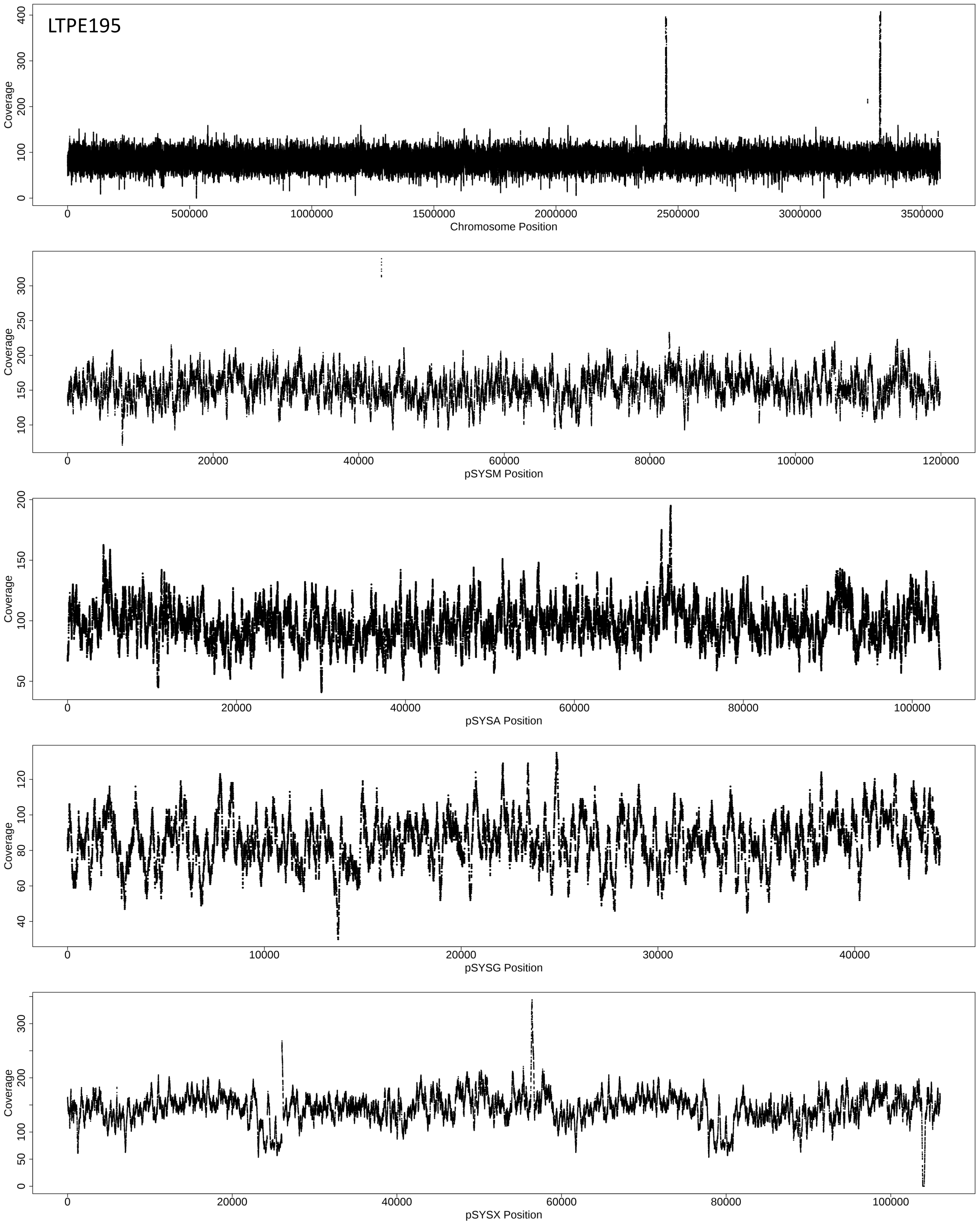

LTPE195
Coverage
0
100
200
300
400
0
500000
1000000
1500000
2000000
2500000
3000000
3500000
Chromosome Position
Coverage
100
150
200
250
300
0
20000
40000
60000
80000
100000
120000
pSYSM Position
Coverage
50
100
150
200
0
20000
40000
60000
80000
100000
pSYSA Position
Coverage
40
60
80
100
120
0
10000
20000
30000
40000
pSYSG Position
Coverage
0
50
100
200
300
0
20000
40000
60000
80000
100000
pSYSX Position

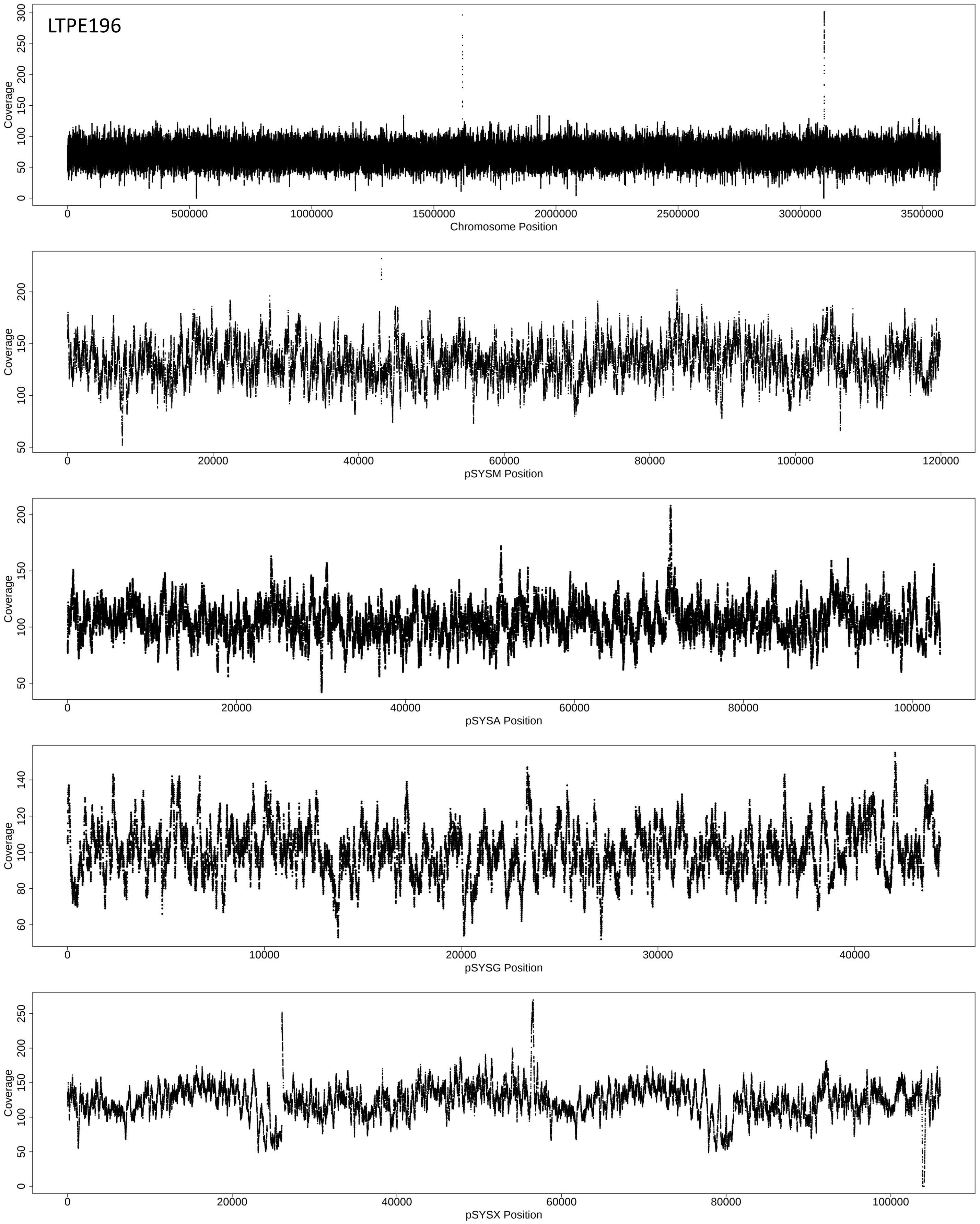
LTPE196
Coverage
Chromosome Position
Coverage
pSYSM Position
Coverage
pSYSA Position
Coverage
pSYSG Position
Coverage
pSYSX Position

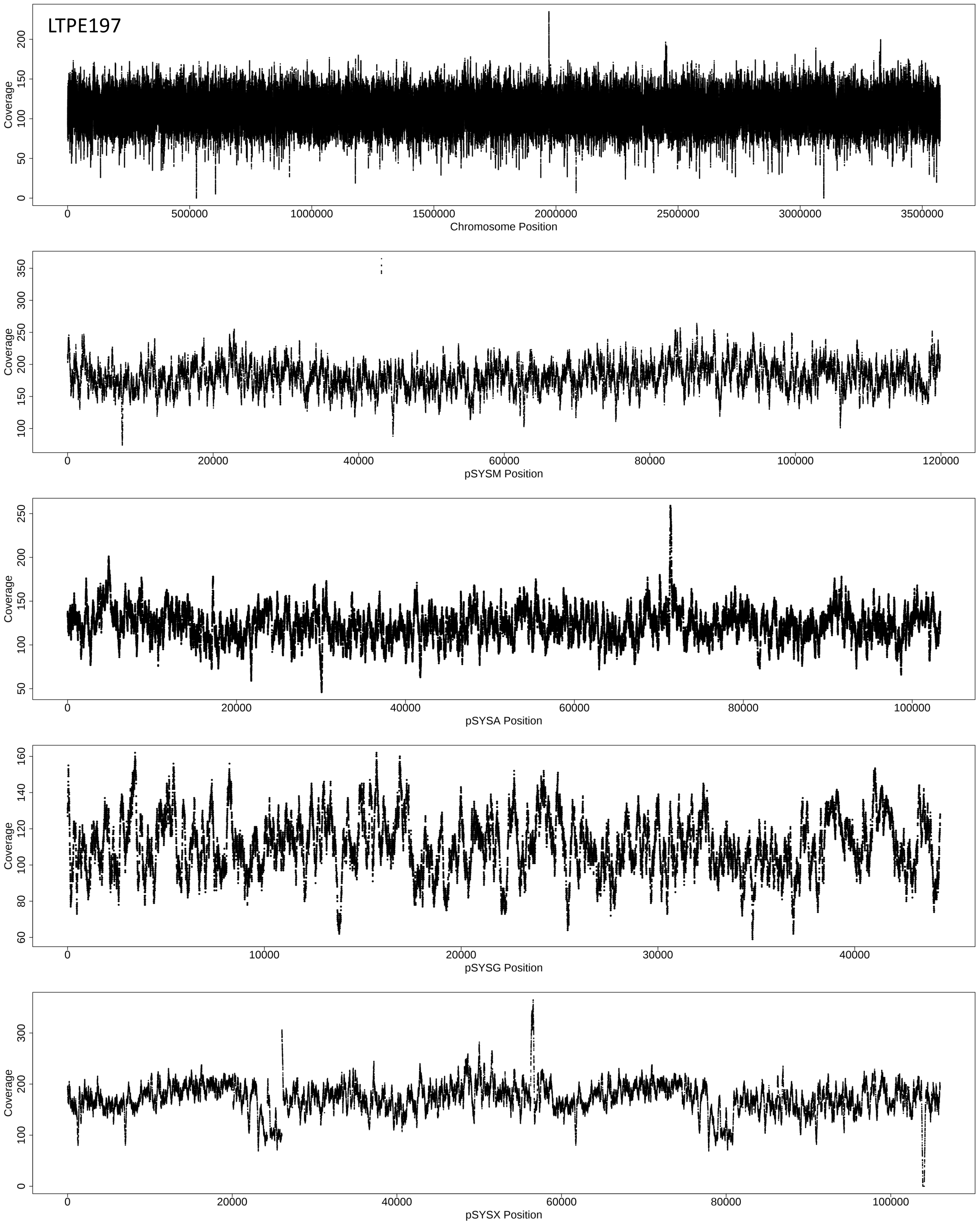
LTPE197
Coverage
0 50 100 150 200
0 500000 1000000 1500000 2000000 2500000 3000000 3500000
Chromosome Position
Coverage
100 150 200 250 300 350
0 20000 40000 60000 80000 100000 120000
pSYSM Position
Coverage
50 100 150 200 250
0 20000 40000 60000 80000 100000
pSYSA Position
Coverage
60 80 100 120 140 160
0 10000 20000 30000 40000
pSYSG Position
Coverage
0 100 200 300
0 20000 40000 60000 80000 100000
pSYSX Position

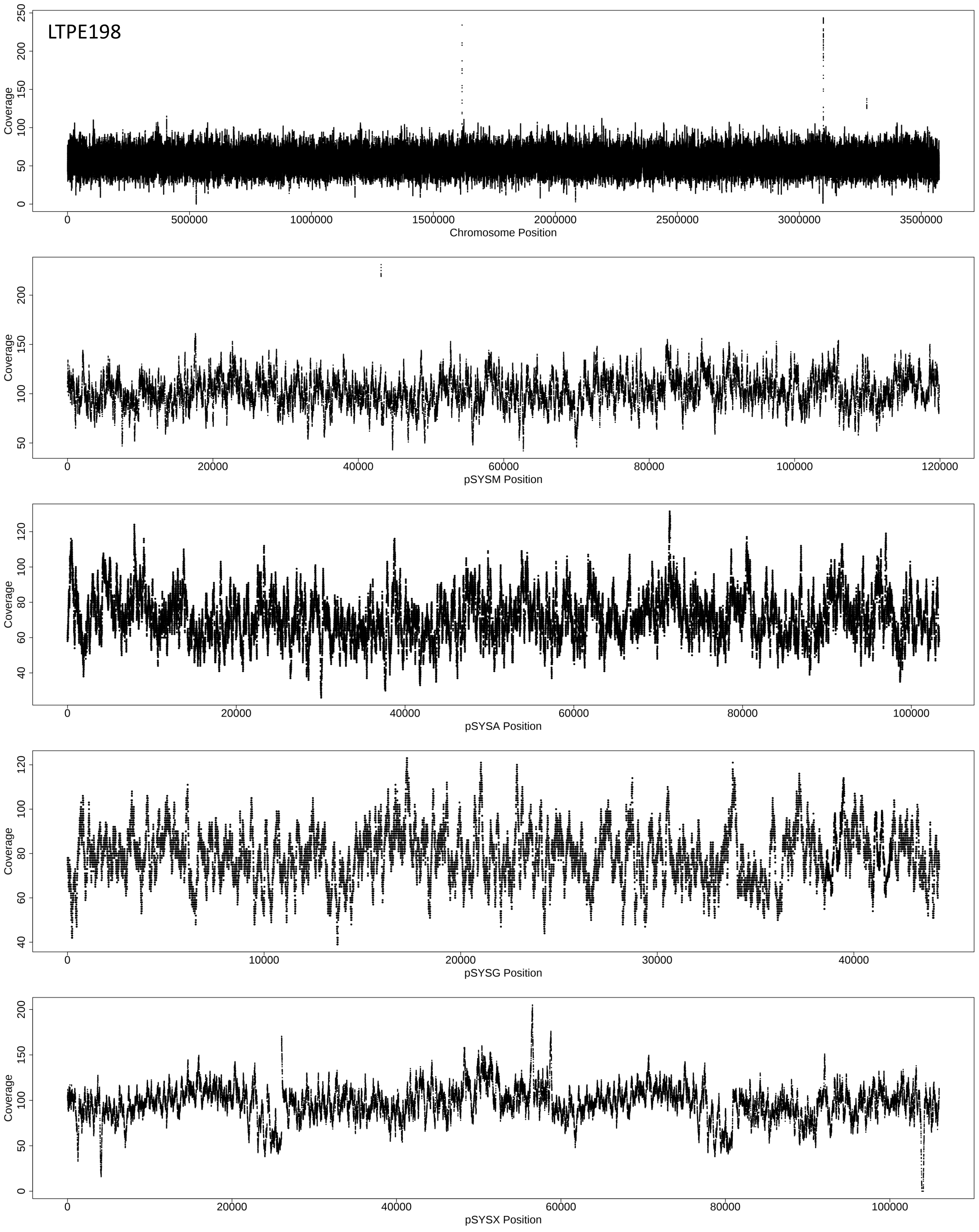
LTPE198
Coverage
0
50
100
150
200
250
0
500000
1000000
1500000
2000000
2500000
3000000
3500000
Chromosome Position
Coverage
50
100
150
200
0
20000
40000
60000
80000
100000
120000
pSYSM Position
Coverage
40
60
80
100
120
0
20000
40000
60000
80000
100000
pSYSA Position
Coverage
40
60
80
100
120
0
10000
20000
30000
40000
pSYSG Position
Coverage
0
50
100
150
200
0
20000
40000
60000
80000
100000
pSYSX Position

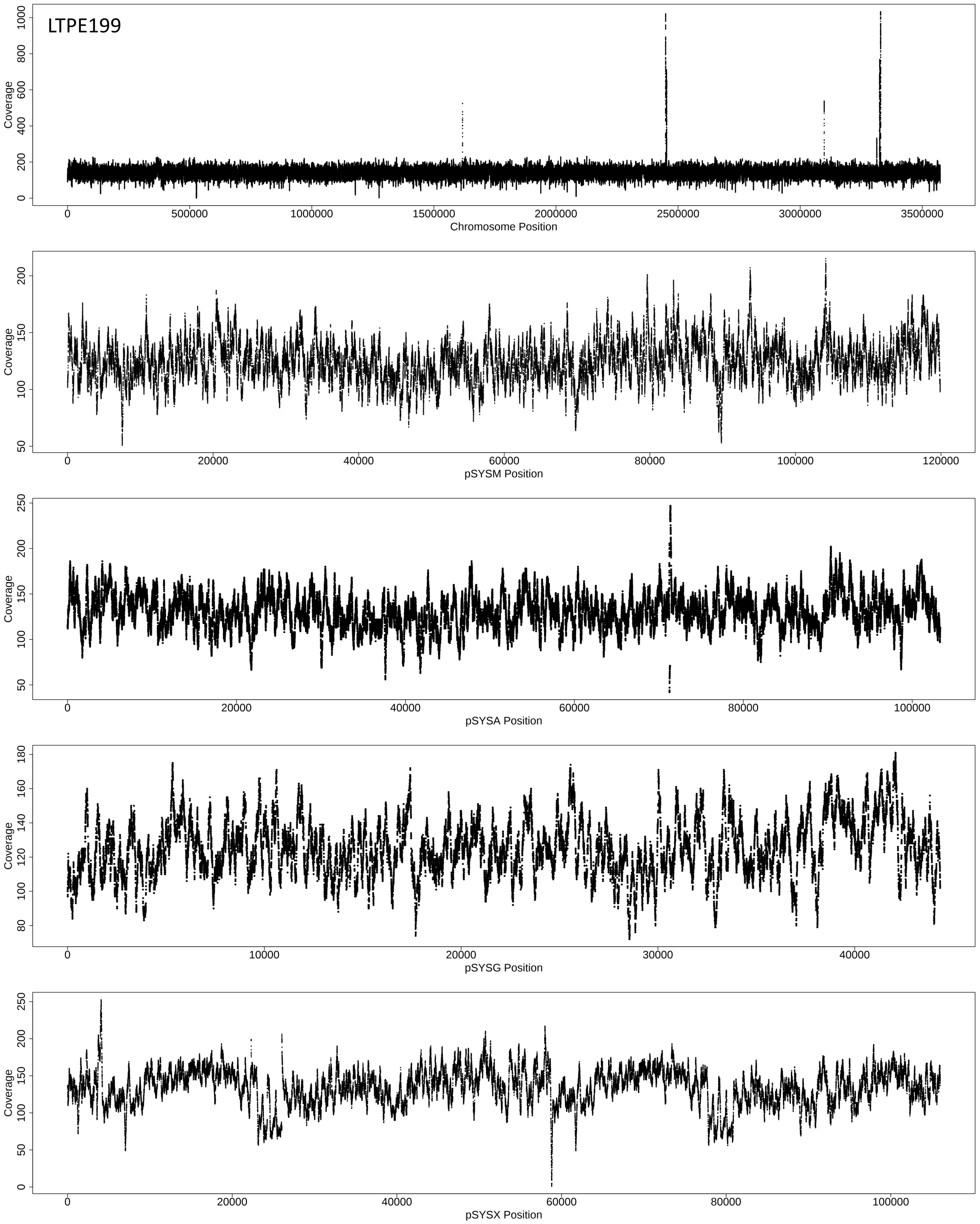
LTPE199
Coverage
0
200
400
600
800
1000
0
500000
1000000
1500000
2000000
2500000
3000000
3500000
Chromosome Position
Coverage
50
100
150
200
0
20000
40000
60000
80000
100000
120000
pSYSM Position
Coverage
50
100
150
200
250
0
20000
40000
60000
80000
100000
pSYSA Position
Coverage
80
100
120
140
160
180
0
10000
20000
30000
40000
pSYSG Position
Coverage
0
50
100
150
200
250
0
20000
40000
60000
80000
100000
pSYSX Position

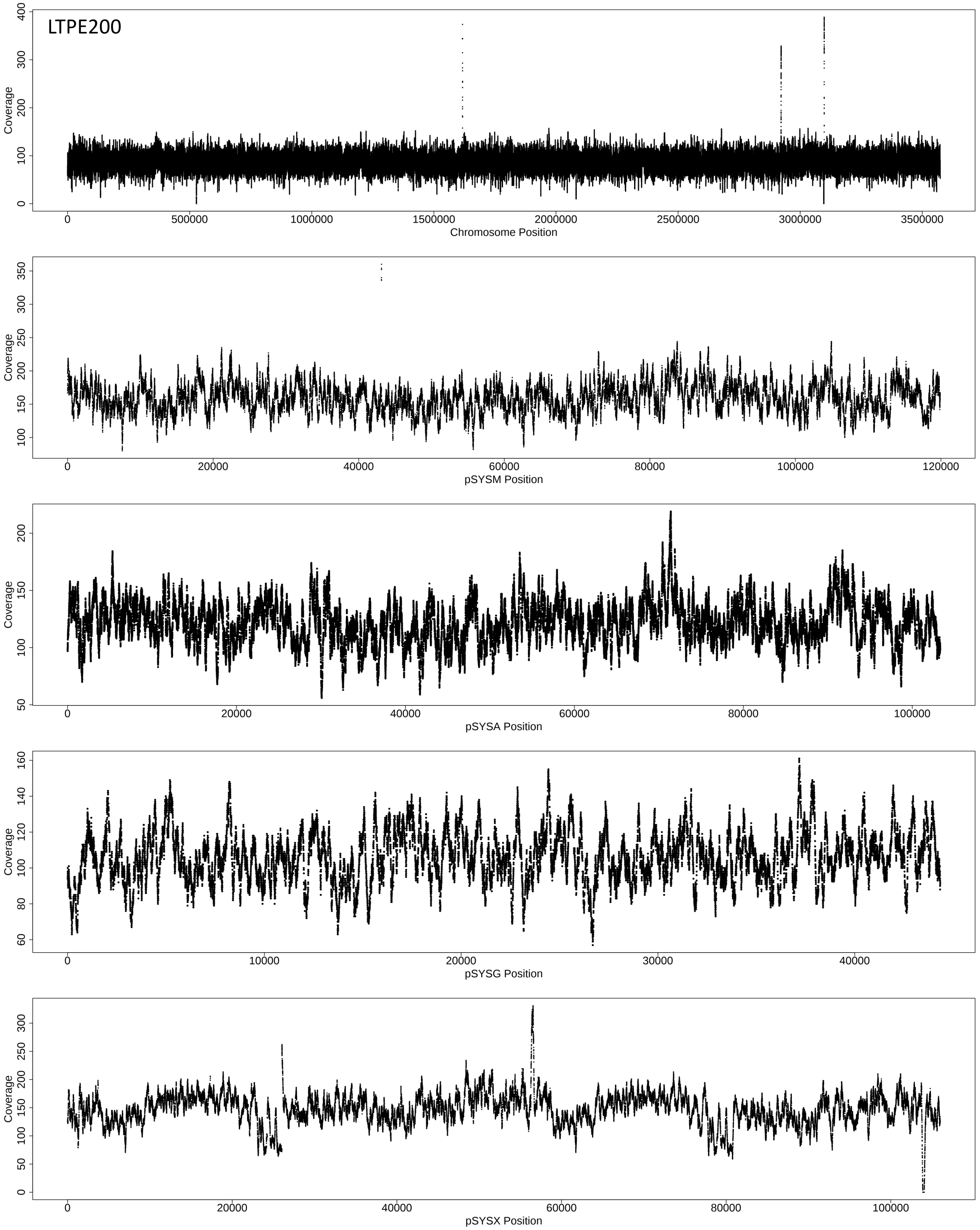

LTPE200
Coverage
Chromosome Position
pSYSM Position
pSYSA Position
pSYSG Position
pSYSX Position

**Figure S5. Sequencing coverage for evolved *Synechocystis* PCC6803 genomes.** Strains LTPE191 to LTPE195 were evolved at 400 ppm $pCO_2$; strains LTPE196 to LTPE200 were evolved at 800 ppm $pCO_2$. For each strain, the main chromosome and each of the four plasmids from the reference genome assembly are shown separately.

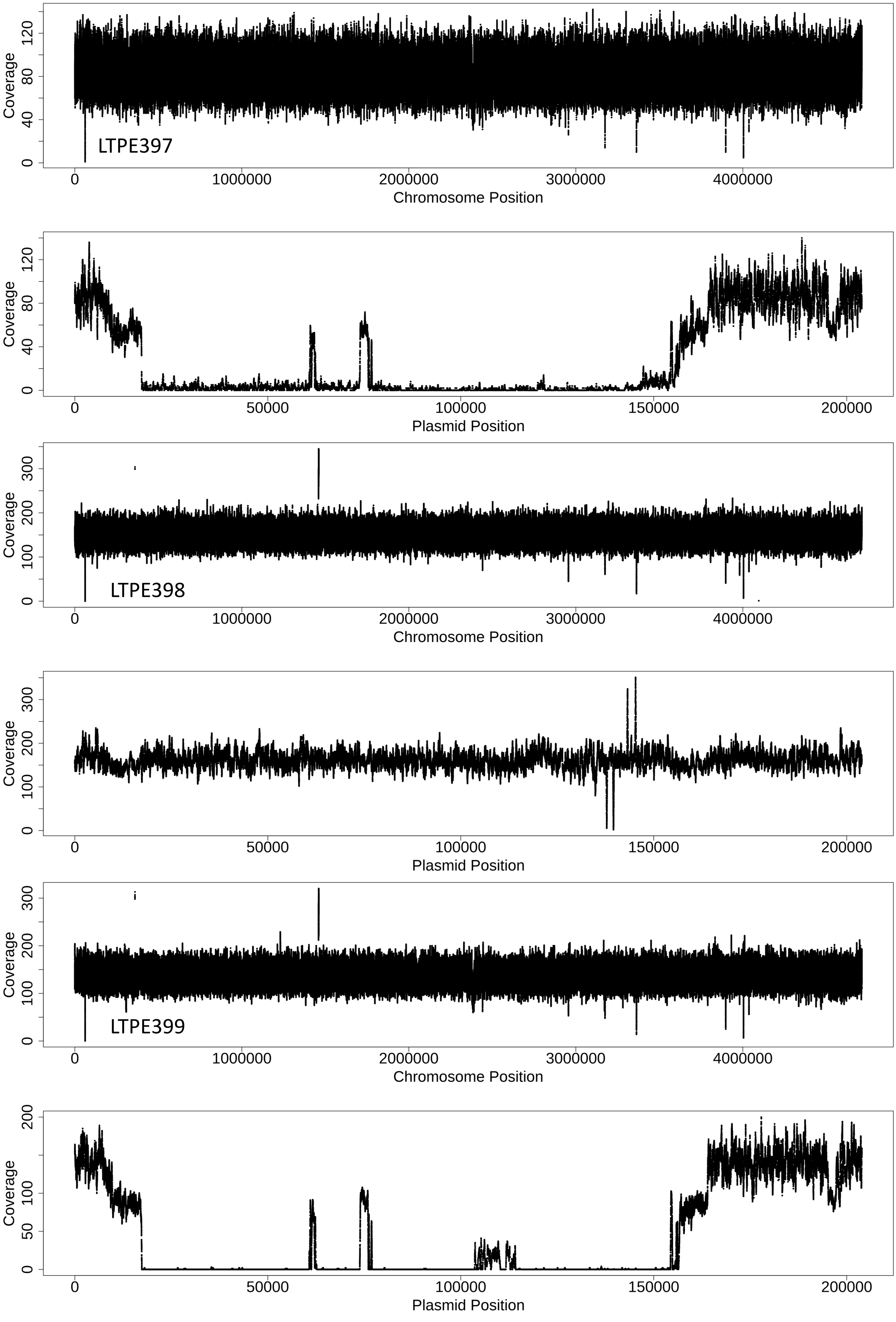

LTPE397
Coverage
0
40
80
120
0
1000000
2000000
3000000
4000000
Chromosome Position
Coverage
Plasmid Position
0
50000
100000
150000
200000
LTPE398
Coverage
0
100
200
300
Chromosome Position
Coverage
Plasmid Position
LTPE399
Coverage
Chromosome Position
Coverage
0
50
100
150
200
Plasmid Position

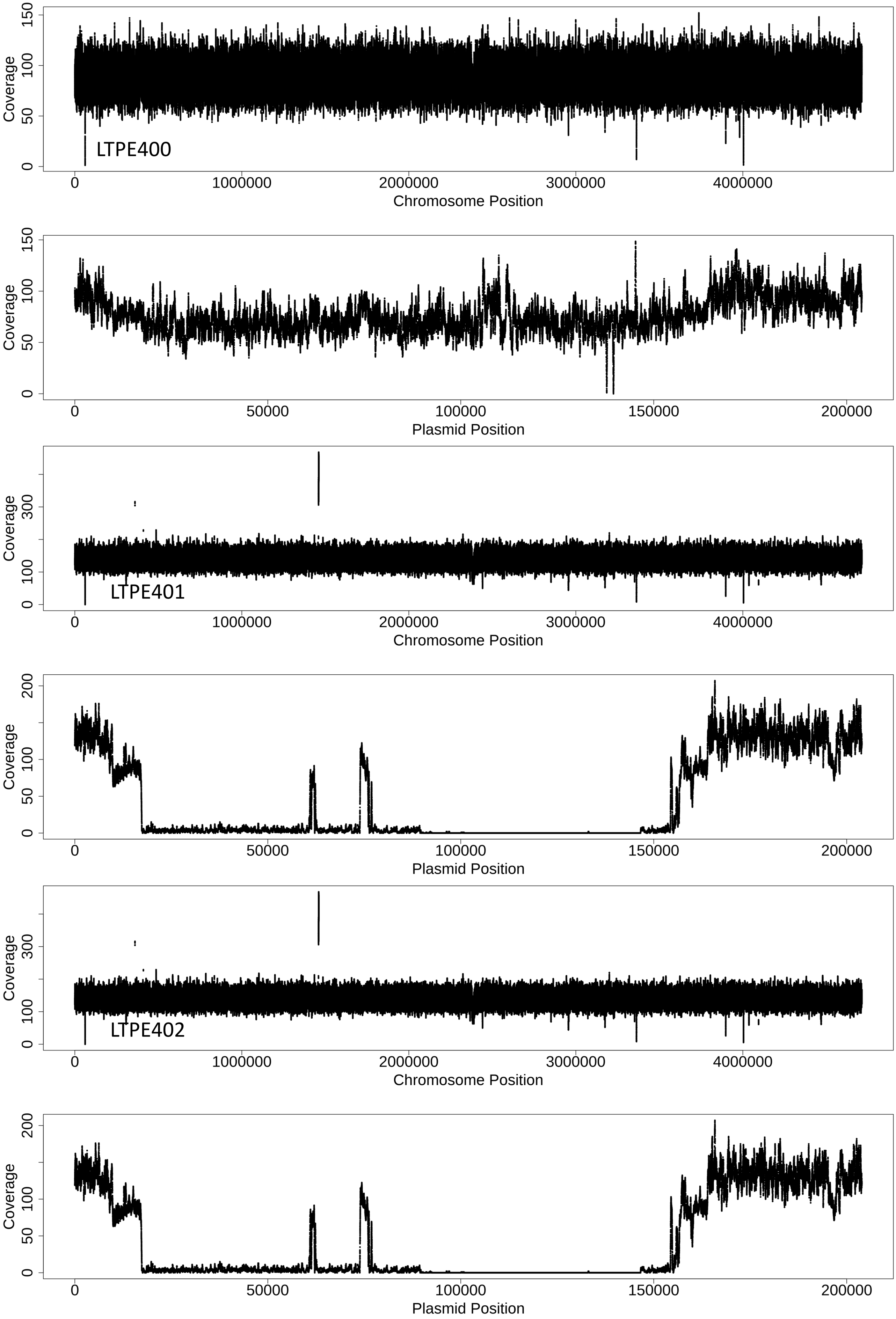
LTPE400
Coverage
0
50
100
150
0
1000000
2000000
3000000
4000000
Chromosome Position
Coverage
0
50
100
150
0
50000
100000
150000
200000
Plasmid Position
LTPE401
Coverage
0
100
300
0
1000000
2000000
3000000
4000000
Chromosome Position
Coverage
0
50
100
200
0
50000
100000
150000
200000
Plasmid Position
LTPE402
Coverage
0
100
300
0
1000000
2000000
3000000
4000000
Chromosome Position
Coverage
0
50
100
200
0
50000
100000
150000
200000
Plasmid Position

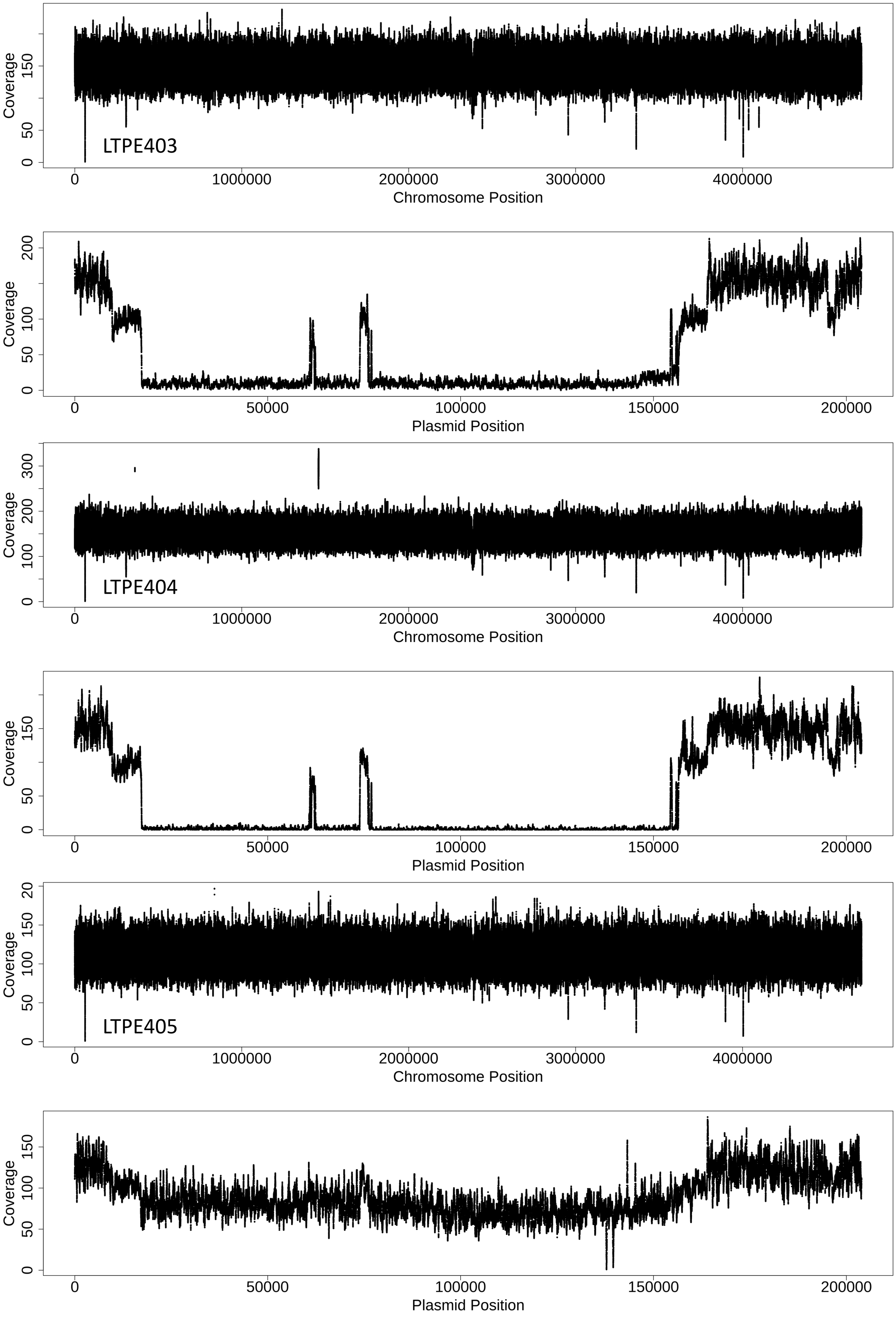
LTPE403
Coverage
0
50
150
Chromosome Position
0
1000000
2000000
3000000
4000000
Coverage
0
50
100
200
Plasmid Position
0
50000
100000
150000
200000
LTPE404
Coverage
0
100
200
300
Chromosome Position
0
1000000
2000000
3000000
4000000
Coverage
0
50
150
Plasmid Position
0
50000
100000
150000
200000
LTPE405
Coverage
0
50
100
150
20
Chromosome Position
0
1000000
2000000
3000000
4000000
Coverage
0
50
100
150
Plasmid Position
0
50000
100000
150000
200000

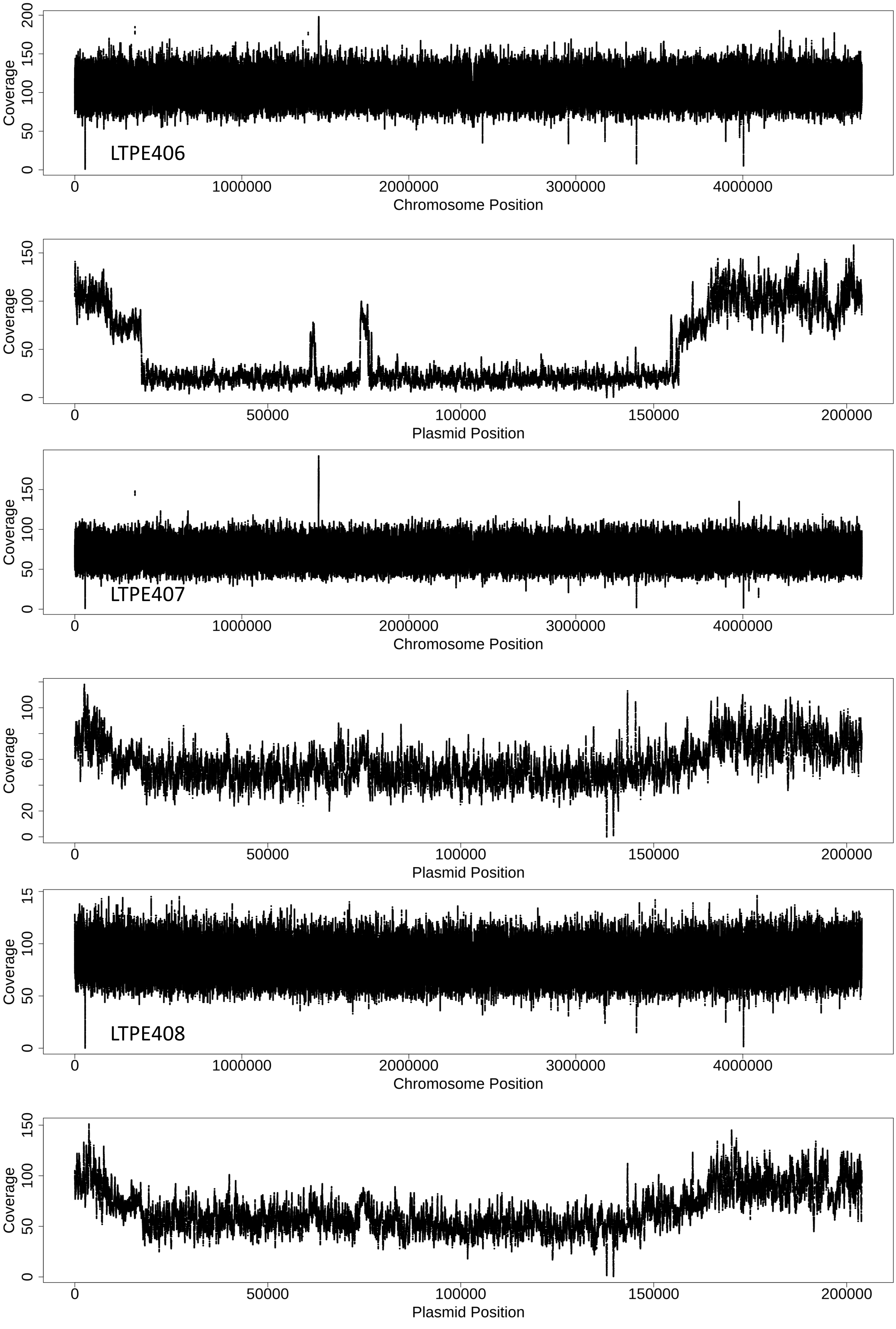
LTPE406
Coverage
0
50
100
150
200
0
1000000
2000000
3000000
4000000
Chromosome Position
Coverage
0
50
100
150
0
50000
100000
150000
200000
Plasmid Position
LTPE407
Coverage
0
50
100
150
0
1000000
2000000
3000000
4000000
Chromosome Position
Coverage
0
20
60
100
0
50000
100000
150000
200000
Plasmid Position
LTPE408
Coverage
0
50
100
15
0
1000000
2000000
3000000
4000000
Chromosome Position
Coverage
0
50
100
150
0
50000
100000
150000
200000
Plasmid Position

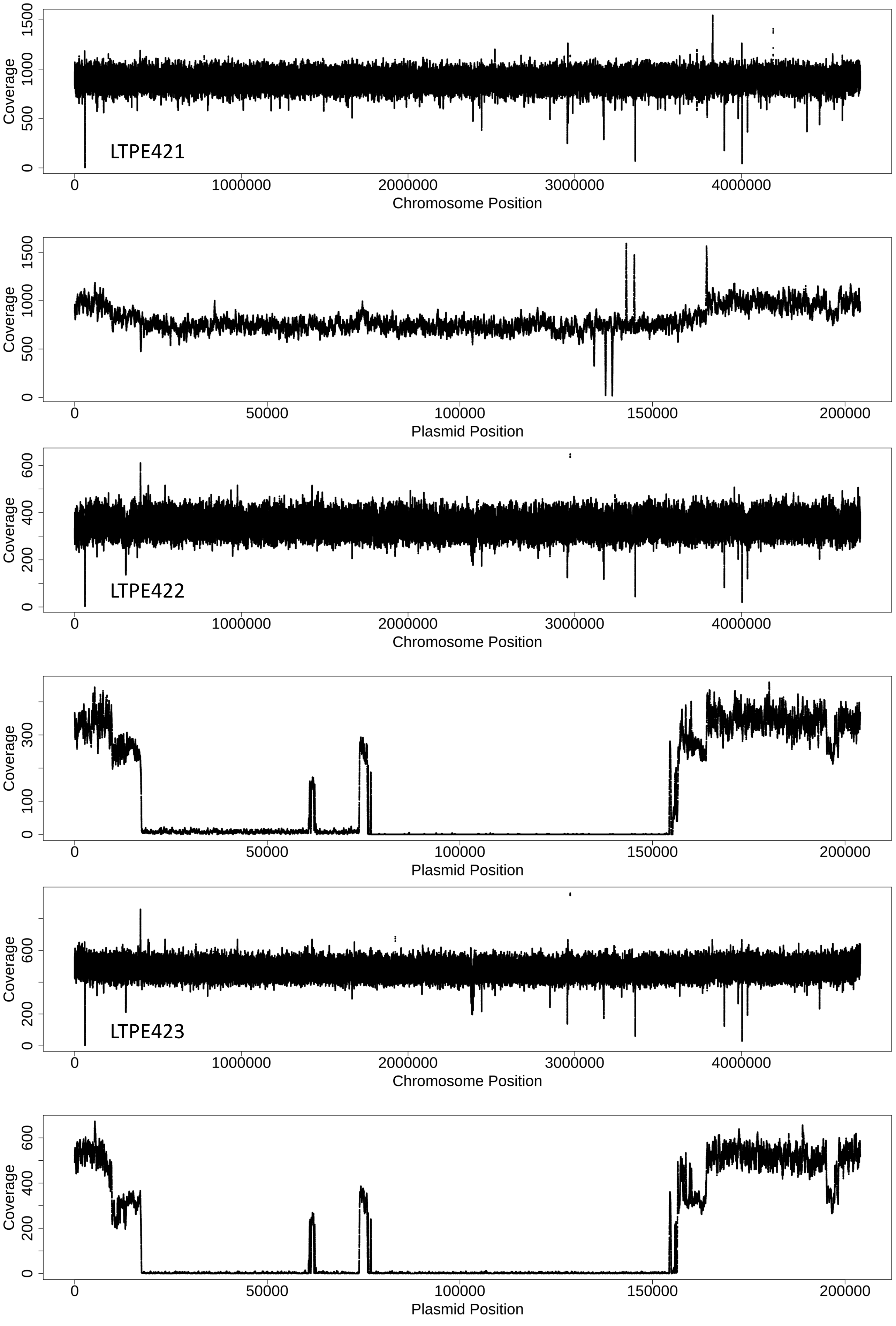
LTPE421
Coverage
0
500
1000
1500
0
1000000
2000000
3000000
4000000
Chromosome Position
Coverage
0
500
1000
1500
0
50000
100000
150000
200000
Plasmid Position
LTPE422
Coverage
0
200
400
600
0
1000000
2000000
3000000
4000000
Chromosome Position
Coverage
0
100
300
0
50000
100000
150000
200000
Plasmid Position
LTPE423
Coverage
0
200
600
0
1000000
2000000
3000000
4000000
Chromosome Position
Coverage
0
200
400
600
0
50000
100000
150000
200000
Plasmid Position

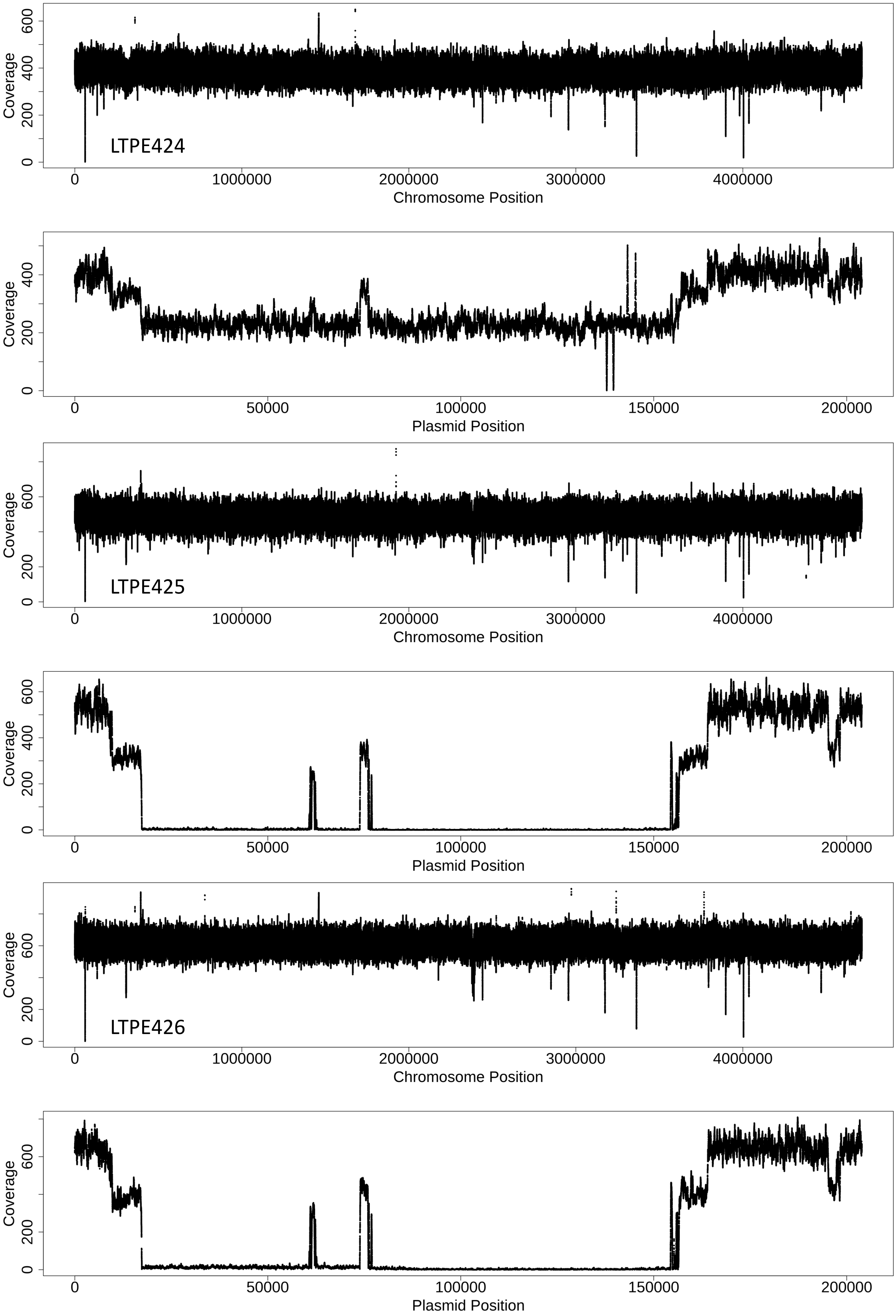
Coverage
0
200
400
600
LTPE424
0
1000000
2000000
3000000
4000000
Chromosome Position
Coverage
0
200
400
0
50000
100000
150000
200000
Plasmid Position
Coverage
0
200
400
600
LTPE425
0
1000000
2000000
3000000
4000000
Chromosome Position
Coverage
0
200
400
600
0
50000
100000
150000
200000
Plasmid Position
Coverage
0
200
400
600
LTPE426
0
1000000
2000000
3000000
4000000
Chromosome Position
Coverage
0
200
400
600
0
50000
100000
150000
200000
Plasmid Position

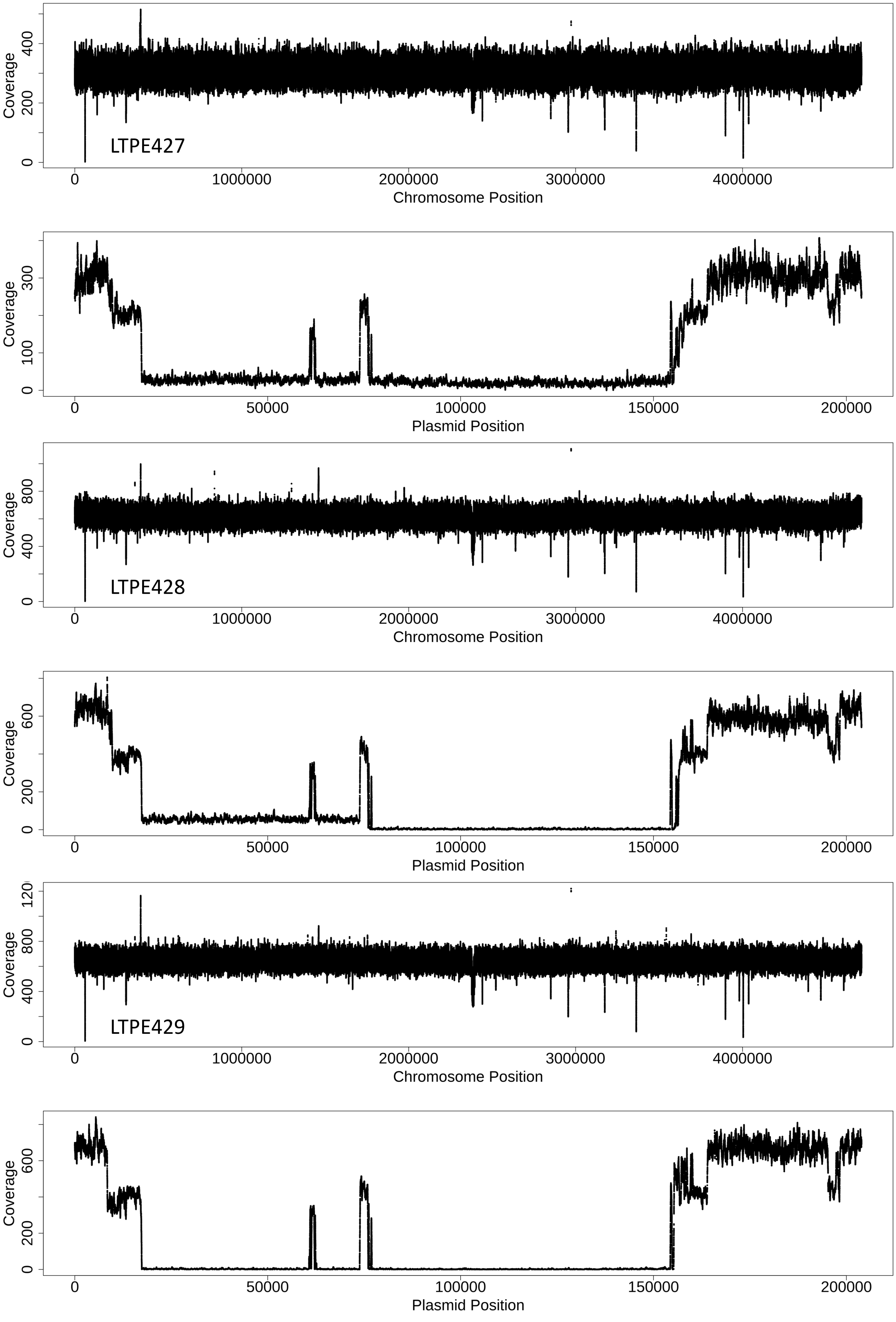
LTPE427
Coverage
0
200
400
0
1000000
2000000
3000000
4000000
Chromosome Position
Coverage
0
100
300
0
50000
100000
150000
200000
Plasmid Position
LTPE428
Coverage
0
400
800
0
1000000
2000000
3000000
4000000
Chromosome Position
Coverage
0
200
600
0
50000
100000
150000
200000
Plasmid Position
LTPE429
Coverage
0
400
800
120
0
1000000
2000000
3000000
4000000
Chromosome Position
Coverage
0
200
600
0
50000
100000
150000
200000
Plasmid Position

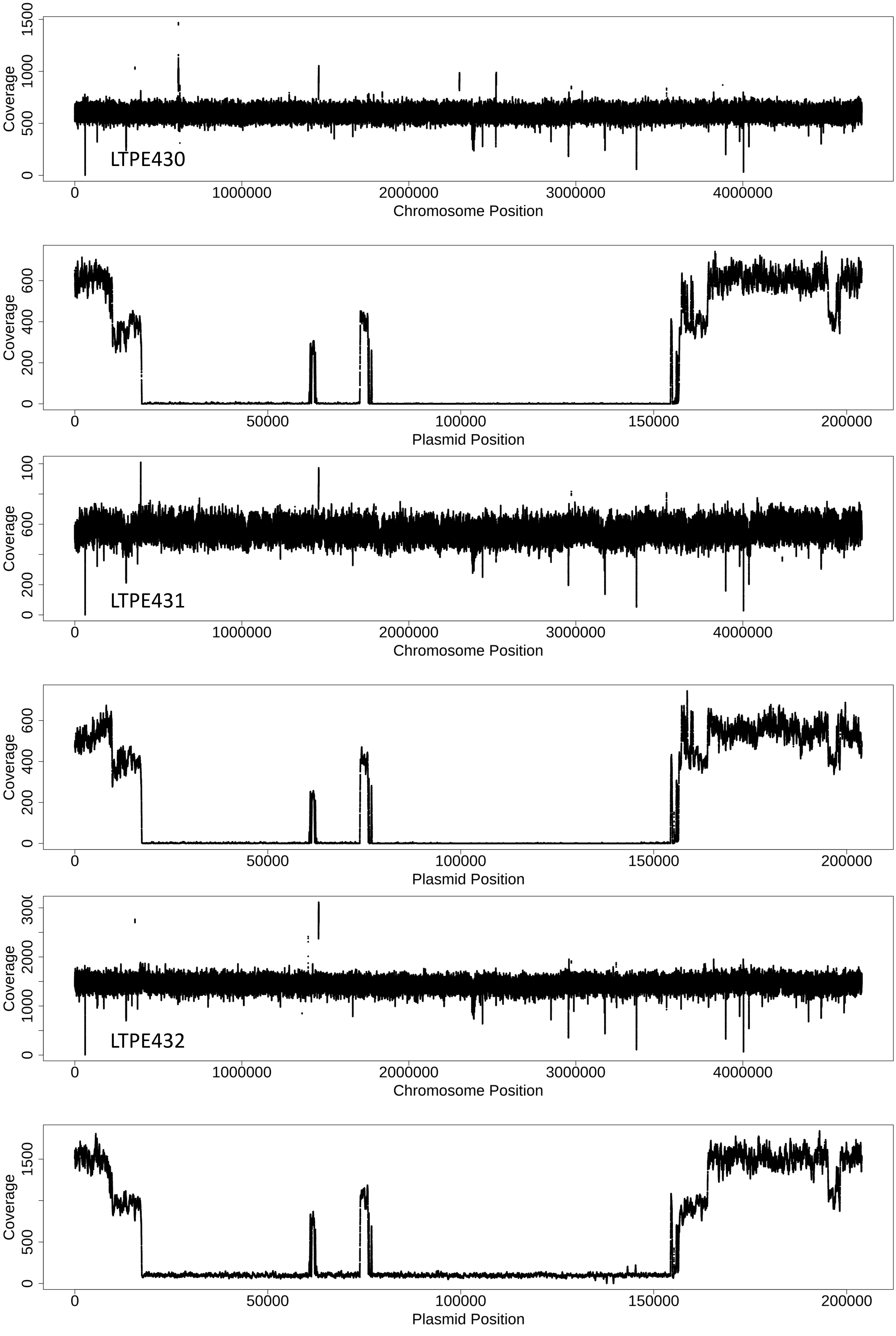

LTPE430
Coverage
0
500
1000
1500
0
1000000
2000000
3000000
4000000
Chromosome Position
Coverage
0
200
400
600
0
50000
100000
150000
200000
Plasmid Position
LTPE431
Coverage
0
200
600
100
0
1000000
2000000
3000000
4000000
Chromosome Position
Coverage
0
200
400
600
0
50000
100000
150000
200000
Plasmid Position
LTPE432
Coverage
0
1000
2000
300
0
1000000
2000000
3000000
4000000
Chromosome Position
Coverage
0
500
1500
0
50000
100000
150000
200000
Plasmid Position

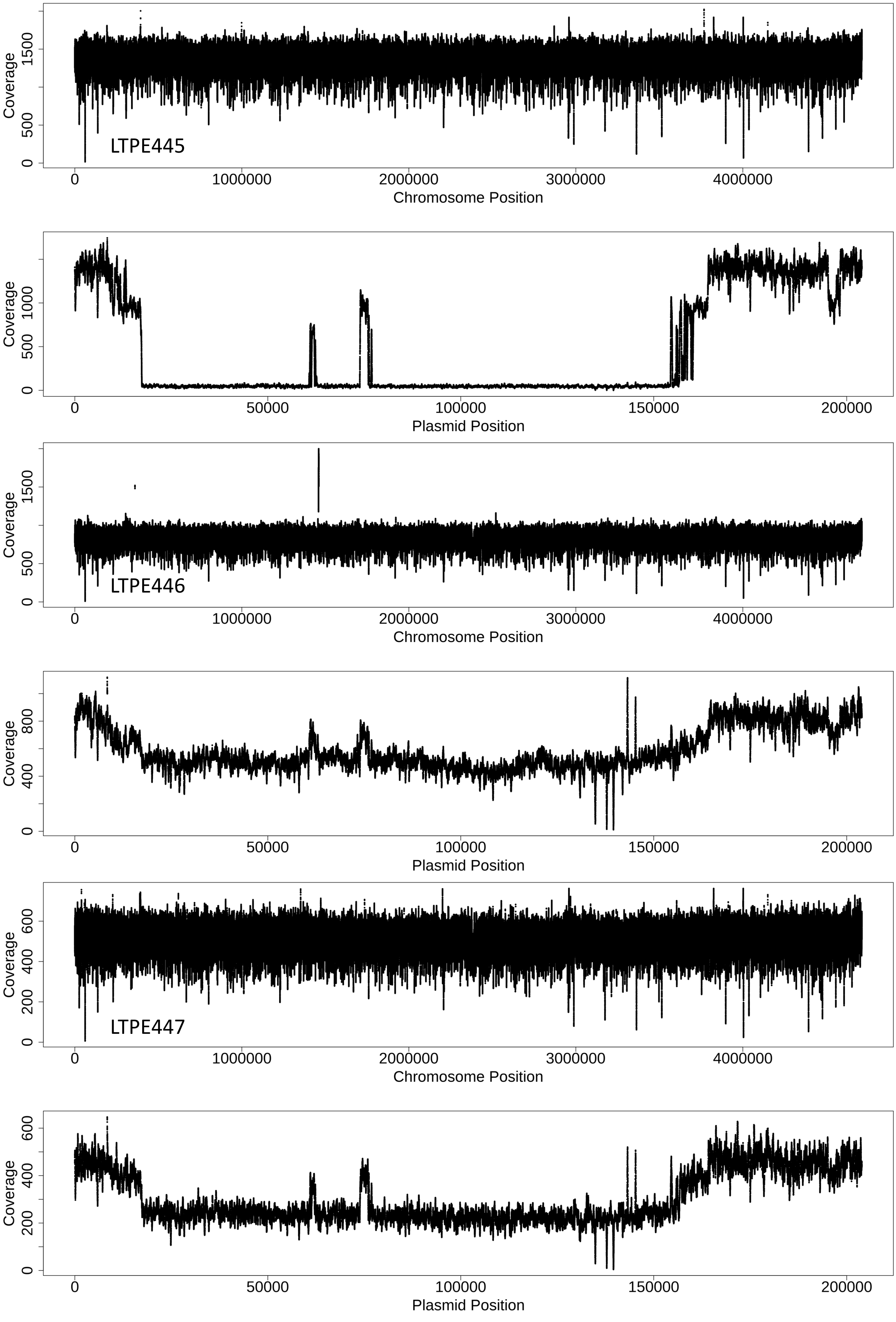
LTPE445
Coverage
0
500
1500
0
1000000
2000000
3000000
4000000
Chromosome Position
Coverage
0
500
1000
0
50000
100000
150000
200000
Plasmid Position
LTPE446
Coverage
0
500
1500
0
1000000
2000000
3000000
4000000
Chromosome Position
Coverage
0
400
800
0
50000
100000
150000
200000
Plasmid Position
LTPE447
Coverage
0
200
400
600
0
1000000
2000000
3000000
4000000
Chromosome Position
Coverage
0
200
400
600
0
50000
100000
150000
200000
Plasmid Position

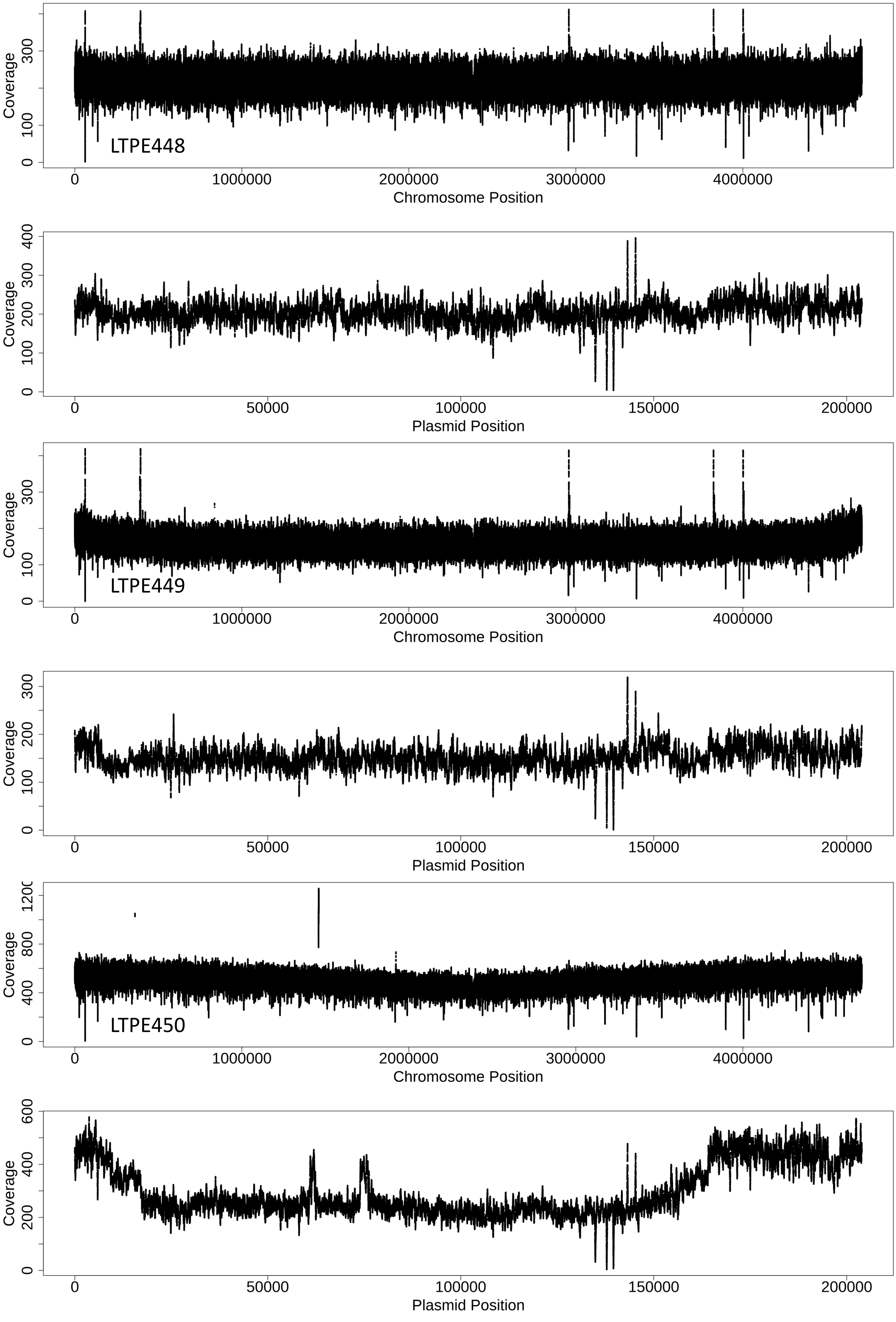
LTPE448
Coverage
0
100
300
0
1000000
2000000
3000000
4000000
Chromosome Position
Coverage
0
100
200
300
400
0
50000
100000
150000
200000
Plasmid Position
LTPE449
Coverage
0
100
300
0
1000000
2000000
3000000
4000000
Chromosome Position
Coverage
0
100
200
300
0
50000
100000
150000
200000
Plasmid Position
LTPE450
Coverage
0
400
800
1200
0
1000000
2000000
3000000
4000000
Chromosome Position
Coverage
0
200
400
600
0
50000
100000
150000
200000
Plasmid Position

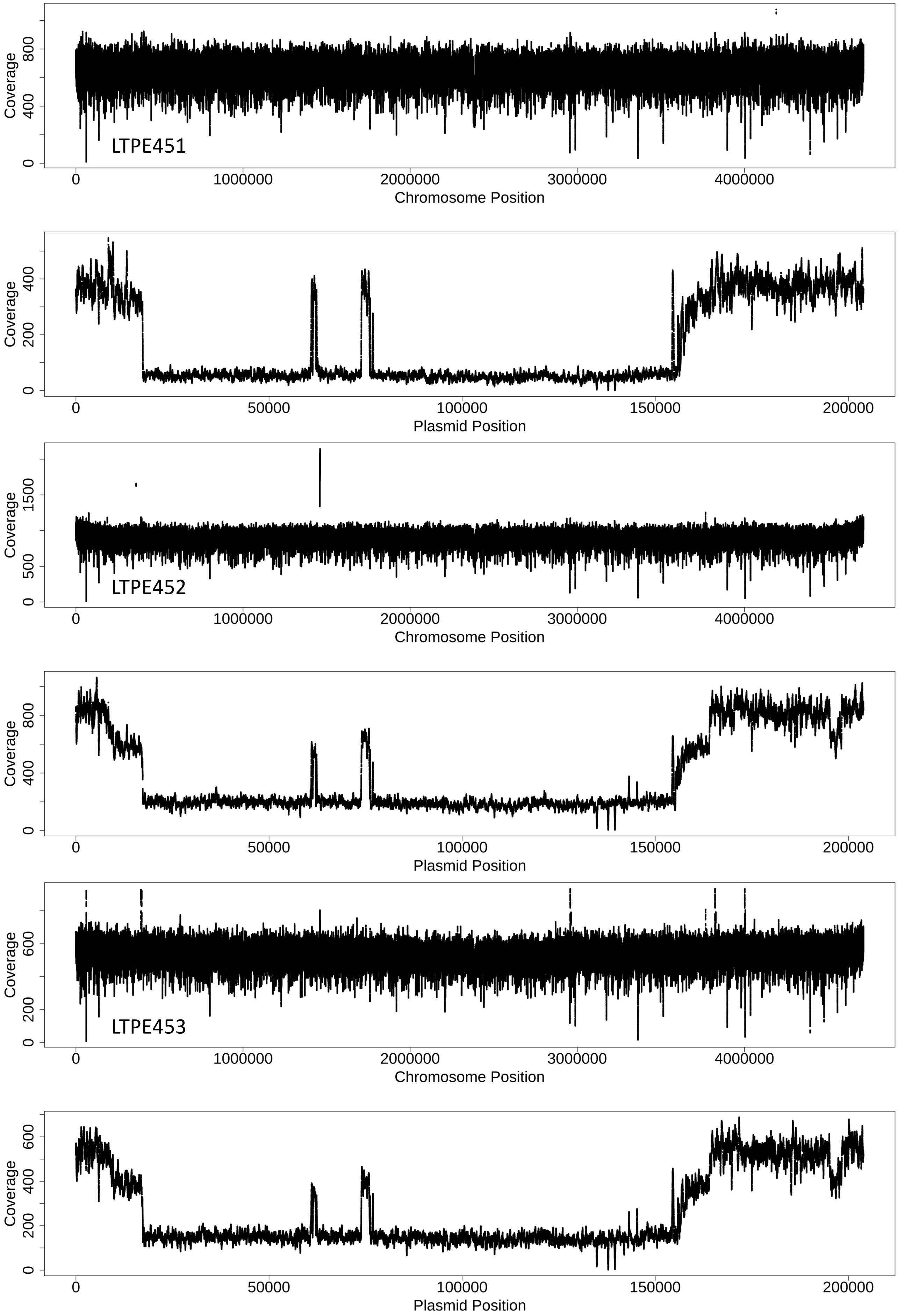

LTPE451
Coverage
Chromosome Position
Plasmid Position
LTPE452
LTPE453
0
200
400
500
600
800
1500
1000000
2000000
3000000
4000000
50000
100000
150000
200000

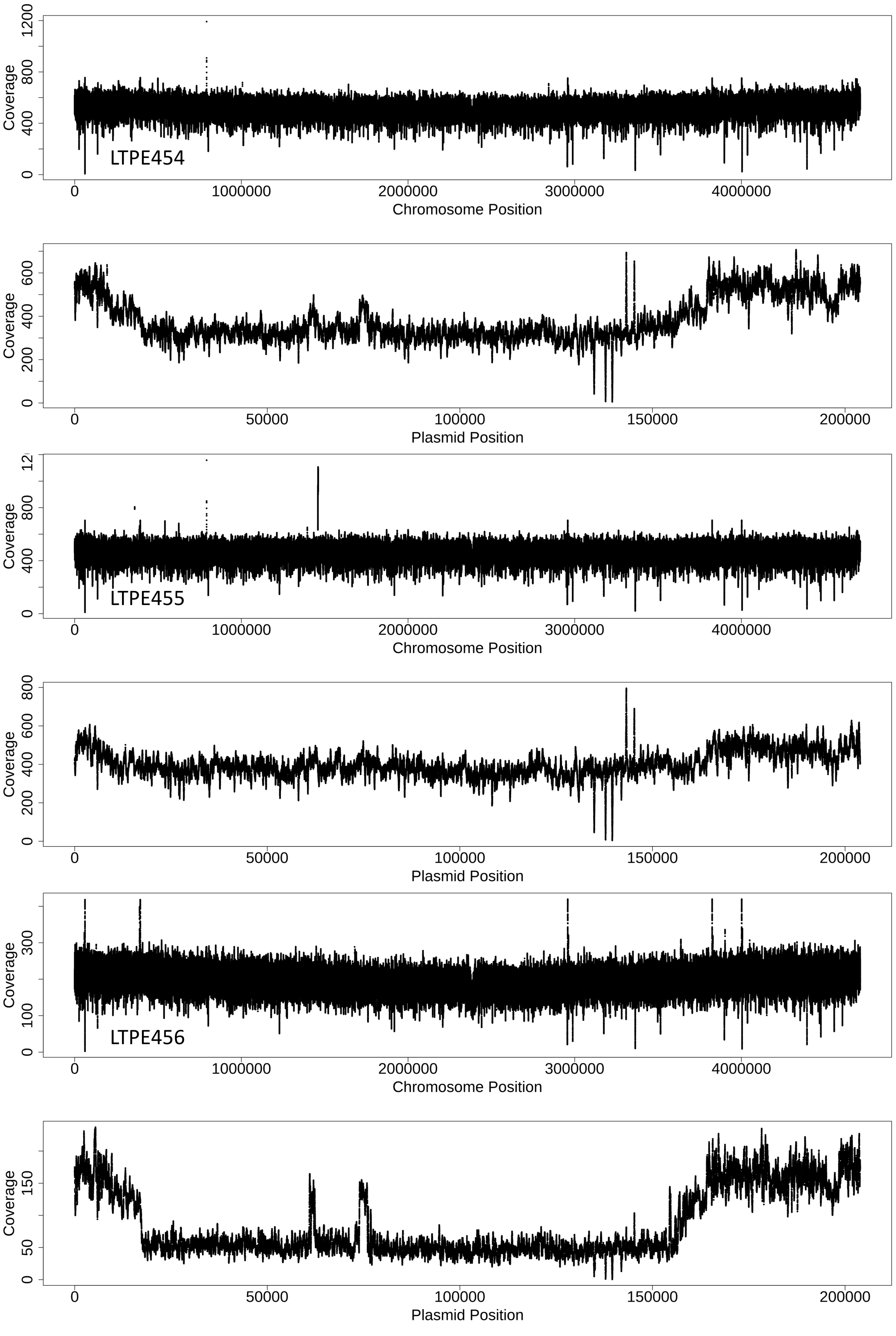
Coverage
LTPE454
Chromosome Position
Coverage
Plasmid Position
Coverage
LTPE455
Chromosome Position
Coverage
Plasmid Position
Coverage
LTPE456
Chromosome Position
Coverage
Plasmid Position

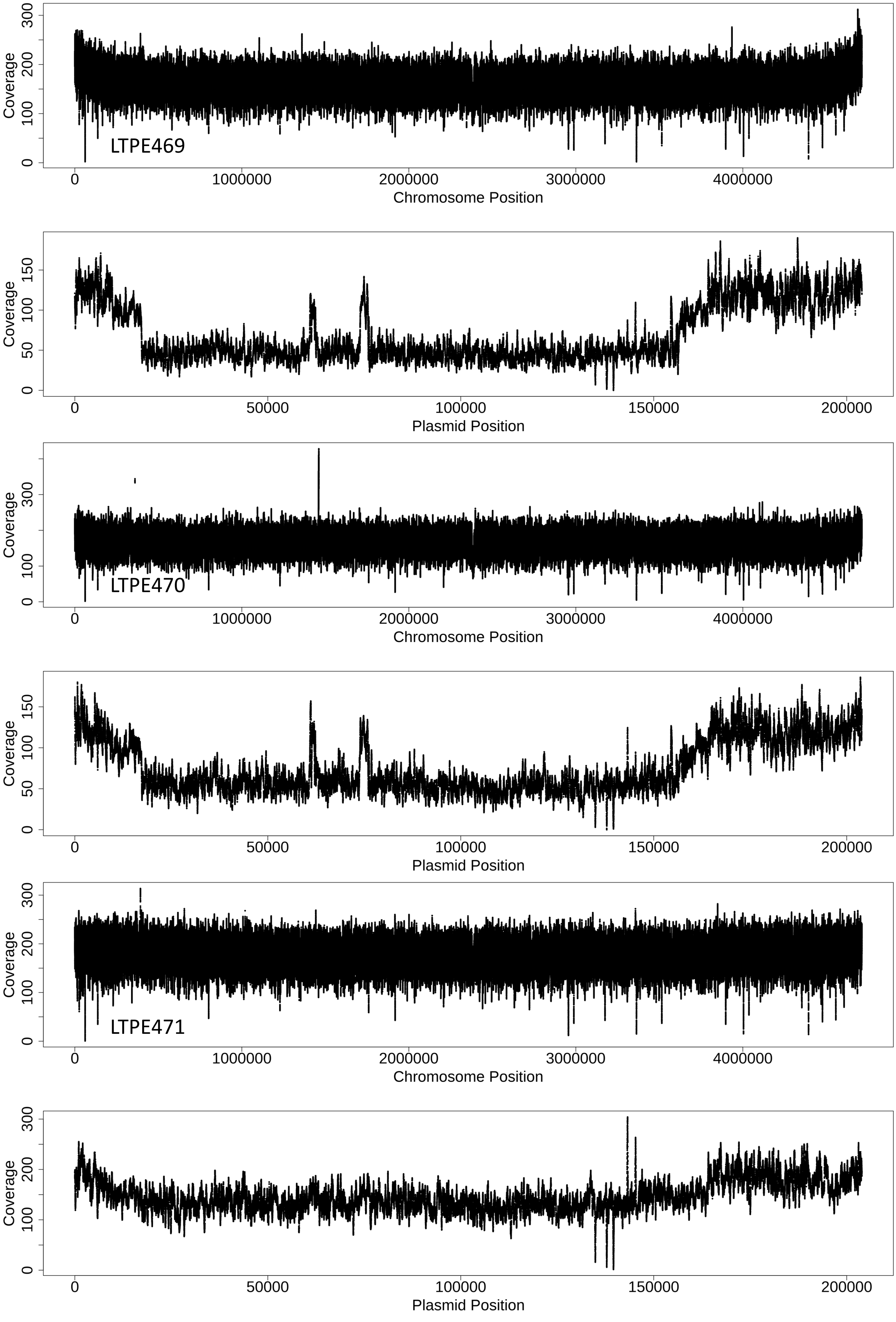
LTPE469
Coverage
Chromosome Position
Plasmid Position
LTPE470
LTPE471
0
50000
100000
150000
200000
1000000
2000000
3000000
4000000
100
200
300
50
150

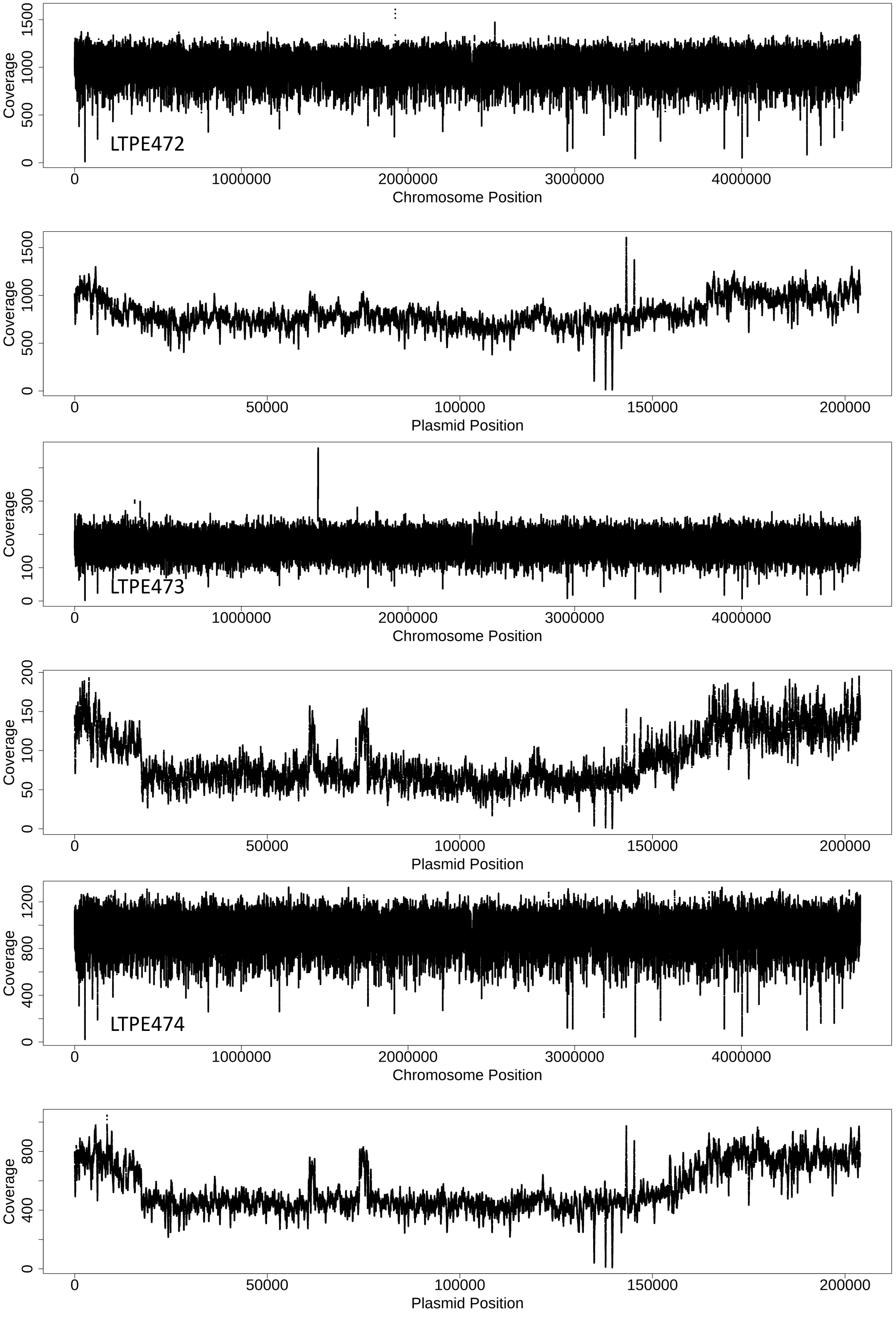

LTPE472
Coverage
0
500
1000
1500
0
1000000
2000000
3000000
4000000
Chromosome Position
Coverage
0
500
1000
1500
0
50000
100000
150000
200000
Plasmid Position
LTPE473
Coverage
0
100
300
0
1000000
2000000
3000000
4000000
Chromosome Position
Coverage
0
50
100
150
200
0
50000
100000
150000
200000
Plasmid Position
LTPE474
Coverage
0
400
800
1200
0
1000000
2000000
3000000
4000000
Chromosome Position
Coverage
0
400
800
0
50000
100000
150000
200000
Plasmid Position

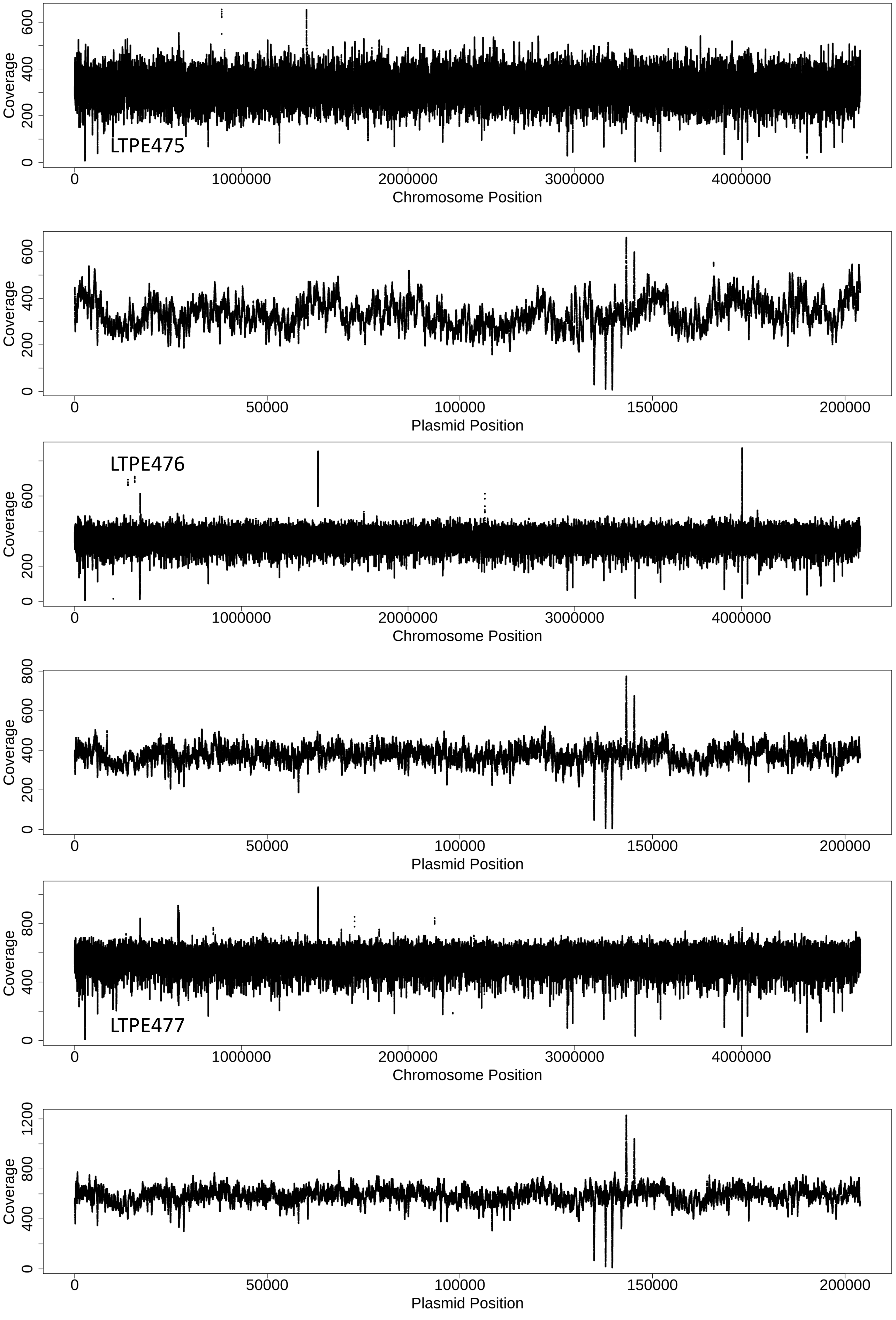
LTPE475
Coverage
Chromosome Position
0
1000000
2000000
3000000
4000000
200
400
600
Plasmid Position
50000
100000
150000
200000
LTPE476
800
LTPE477
1200

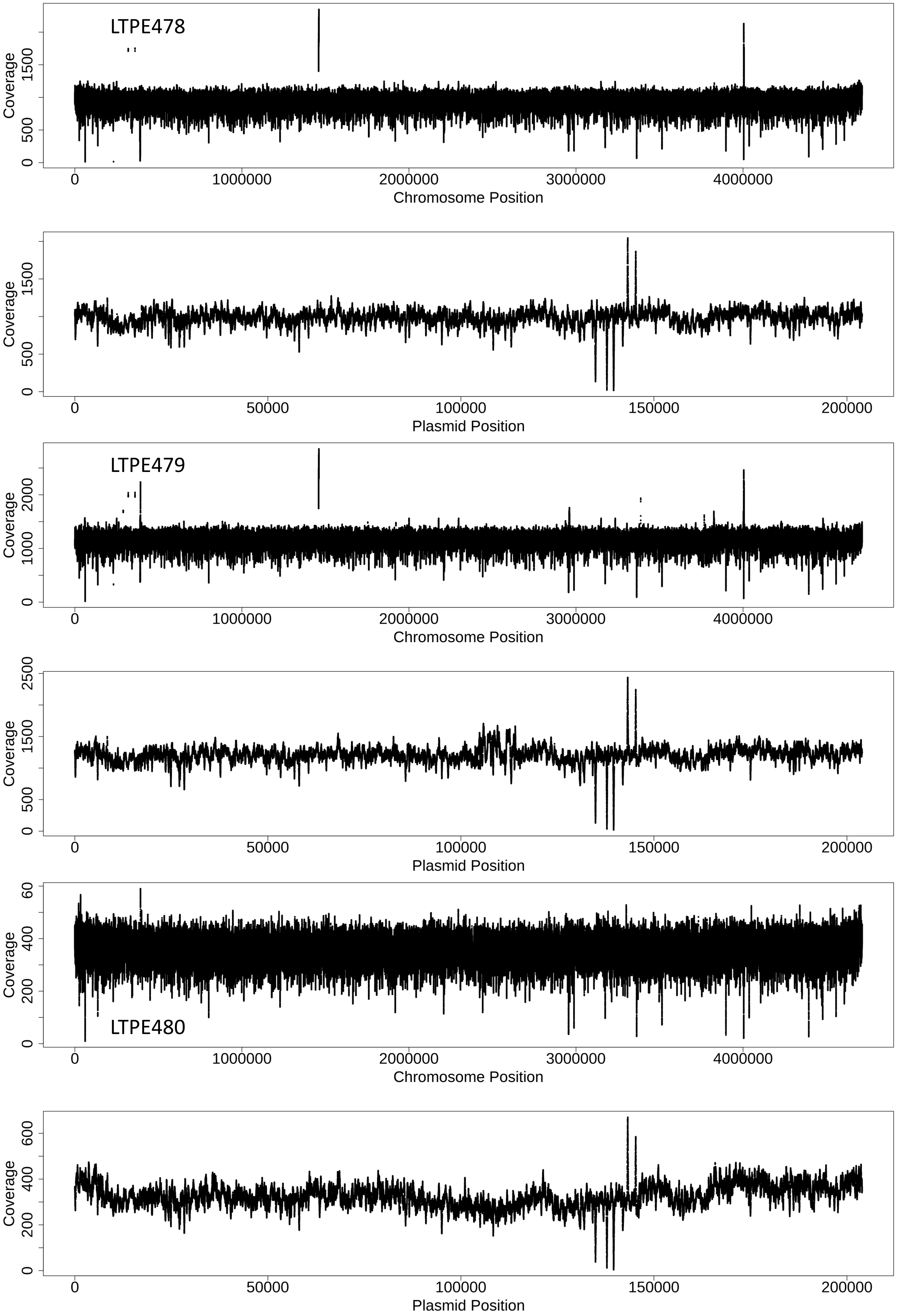

LTPE478
Coverage
0
500
1500
0
1000000
2000000
3000000
4000000
Chromosome Position
Coverage
0
500
1500
0
50000
100000
150000
200000
Plasmid Position
LTPE479
Coverage
0
1000
2000
0
1000000
2000000
3000000
4000000
Chromosome Position
Coverage
0
500
1500
2500
0
50000
100000
150000
200000
Plasmid Position
LTPE480
Coverage
0
200
400
60
0
1000000
2000000
3000000
4000000
Chromosome Position
Coverage
0
200
400
600
0
50000
100000
150000
200000
Plasmid Position

**Figure S6. Sequencing coverage for evolved *Alteromonas* EZ55 genomes.** Strain numbers indicate the designation of the phytoplankton partner. For each strain, the main chromosome and plasmid are shown separately. “Islands” of coverage in the middle portion of the plasmid represent areas of homology with the main chromosome that caused breseq to mistakenly assign a fraction of coverage to regions of the plasmid that were likely deleted during evolution (see Supplemental Figures S20 and S21).

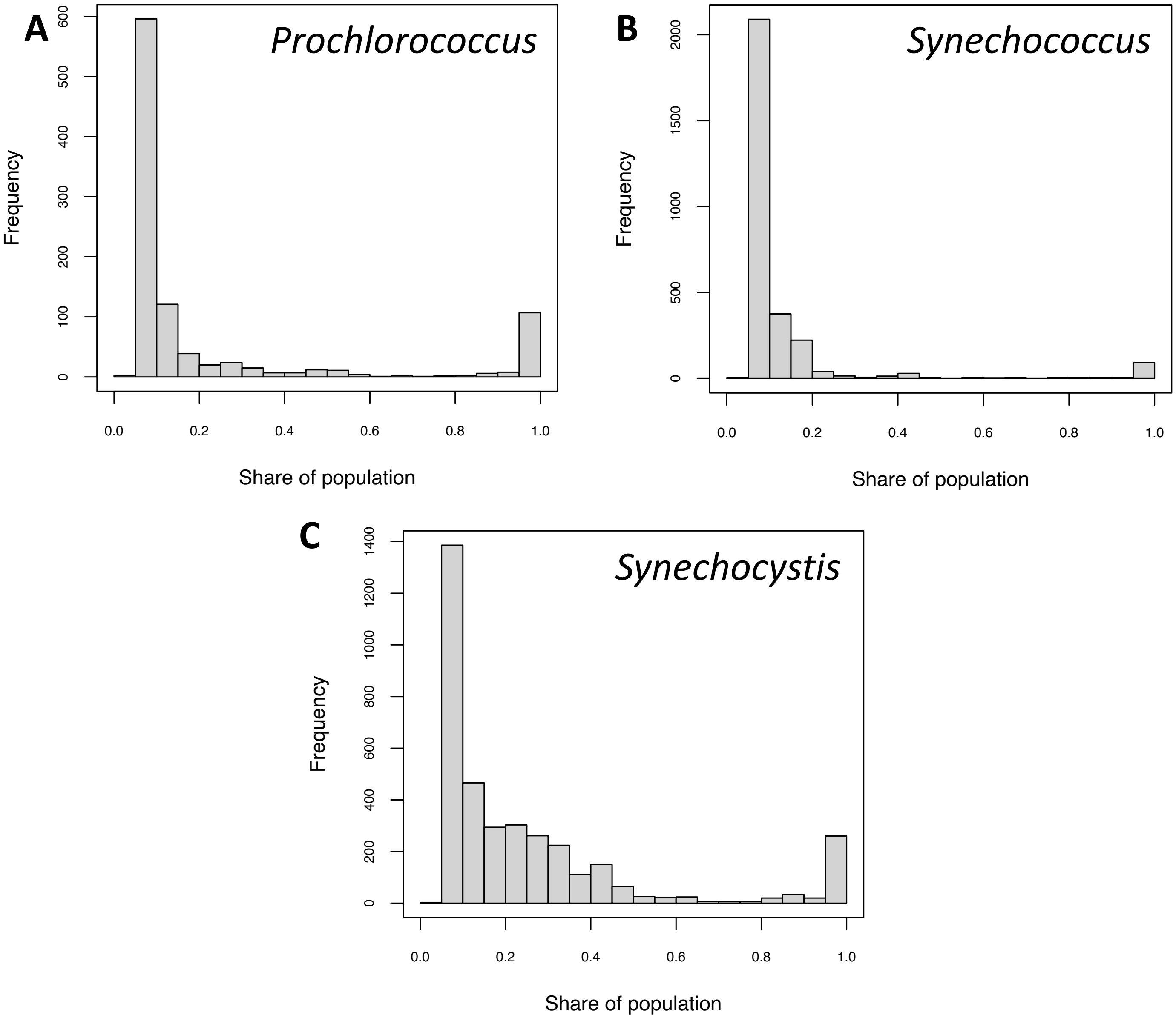


**Figure S7. Frequency distributions of mutations in cyanobacterial genomes.** Histograms show the frequencies of mutations present as a given percentage share of the population, summed across all replicate evolved lineages for A) *Prochlorococcus* MIT9312, B) *Synechococcus* CC9311, or C) *Synechocystis* PCC6803.

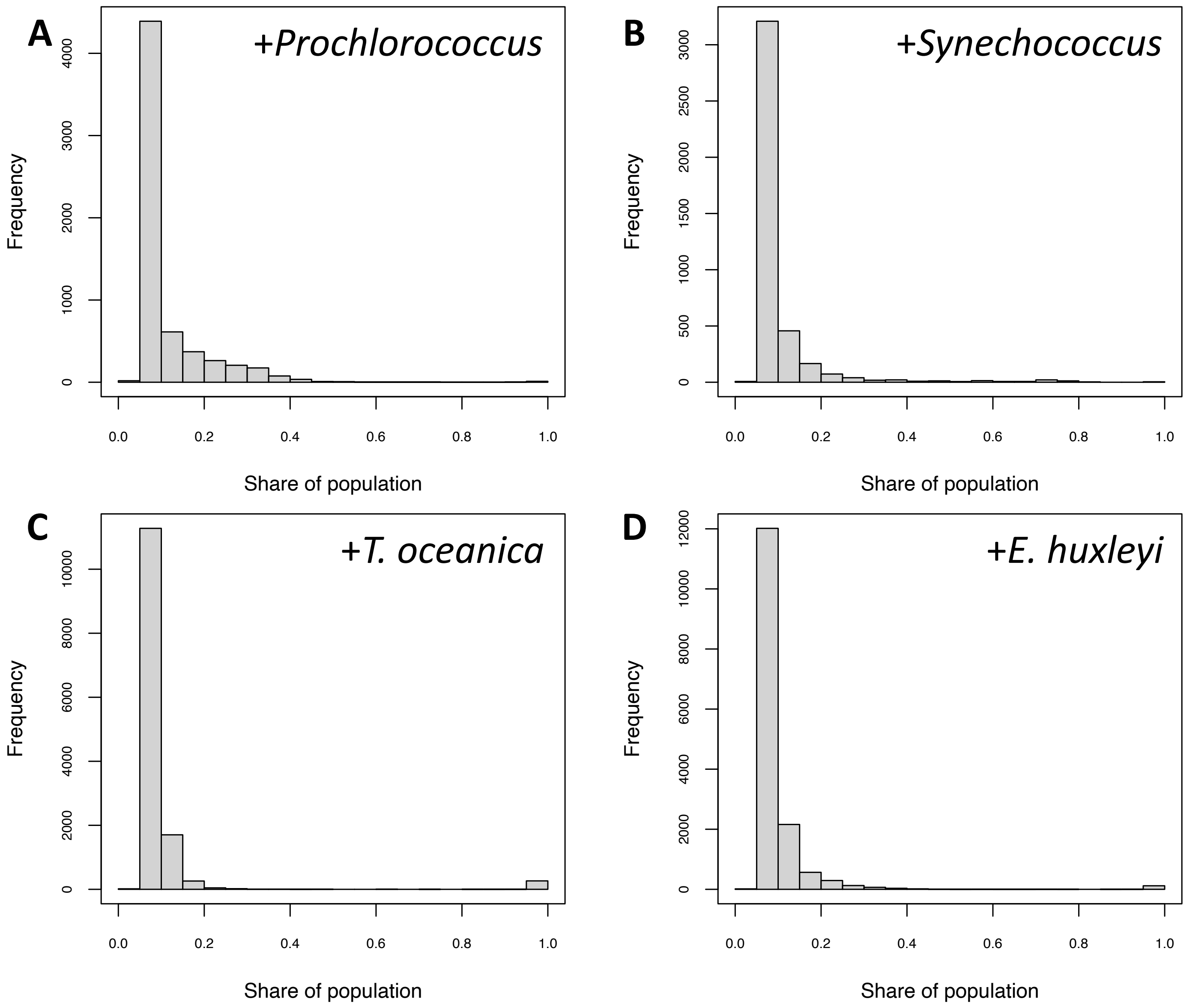


**Figure S8. Frequency distributions of mutations in *Alteromonas* genomes.** Histograms show the frequencies of mutations present as a given percentage share of the population, summed across all replicate evolved lineages for a given organism. Individual plots represent *Alteromonas* EZ55 genomes evolved alongside A) *Prochlorococcus* MIT9312, B) *Synechococcus* CC9311, C) *Thalassiosira oceanica* CCMP1005, or D) *Emiliania huxleyi* CCMP371.

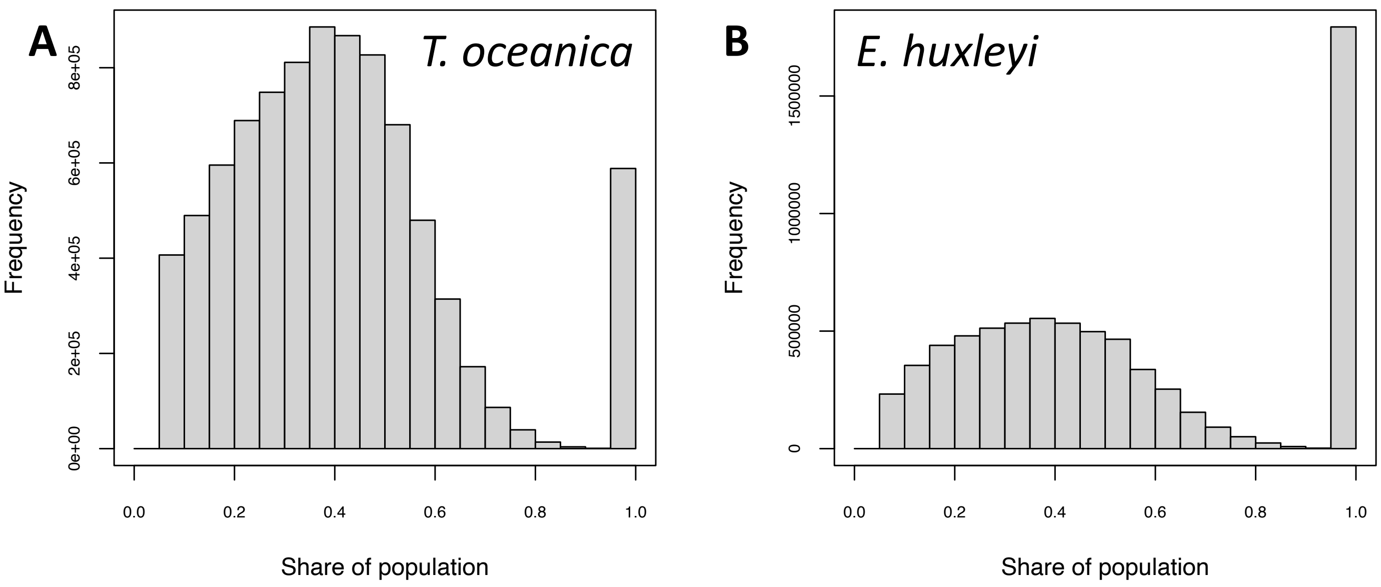


**Figure S9. Frequency distributions of mutations in eukaryotic phytoplankton genomes.** Histograms show the frequencies of mutations present as a given percentage share of the population, summed across all replicate evolved lineages for A) *Thalassiosira oceanica* CCMP1005 or B) *emiliania huxleyi* CCMP371.

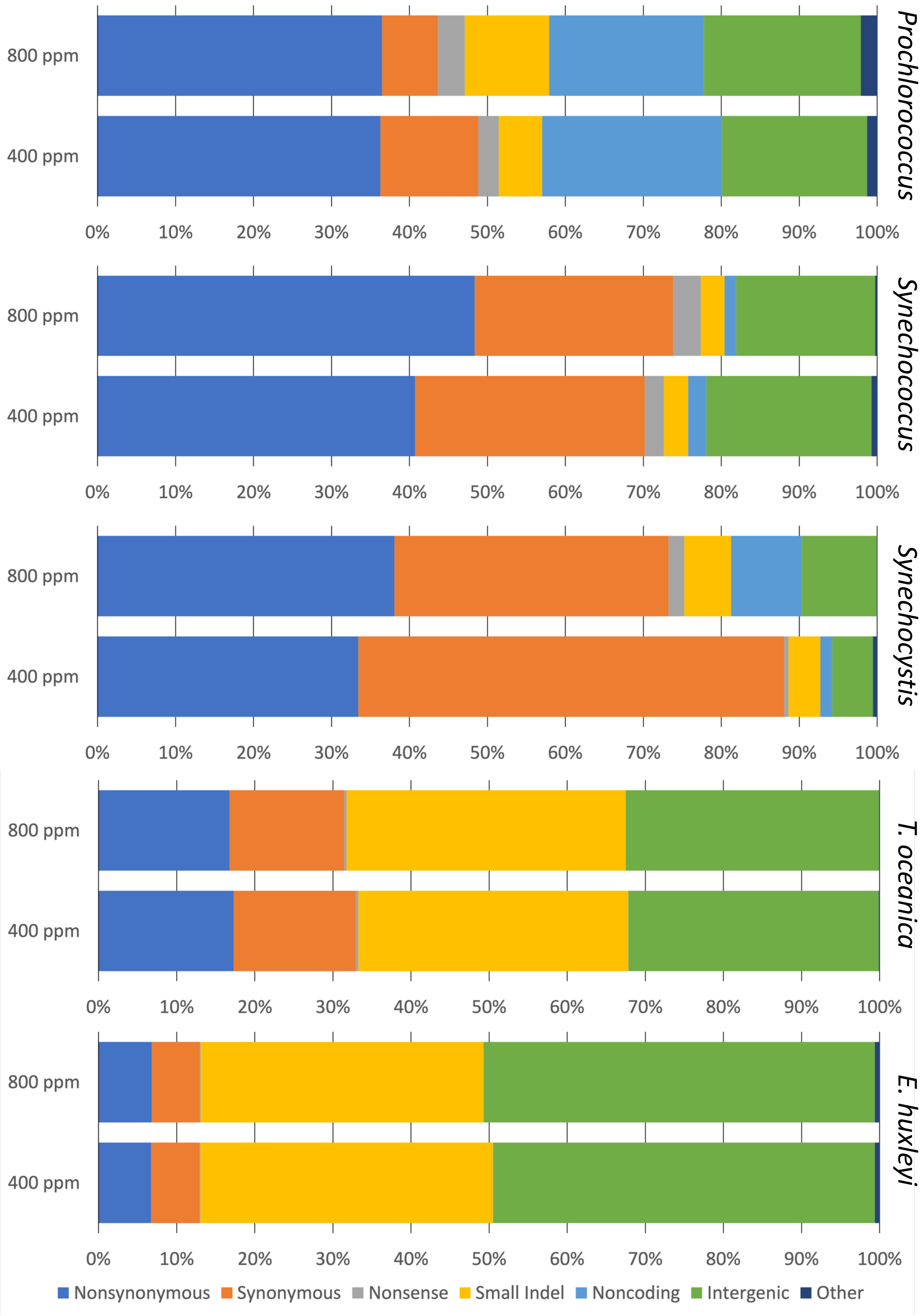


**Figure S10. Distributions of mutation types in phytoplankton genomes evolved under different $pCO_2$ regimes.**

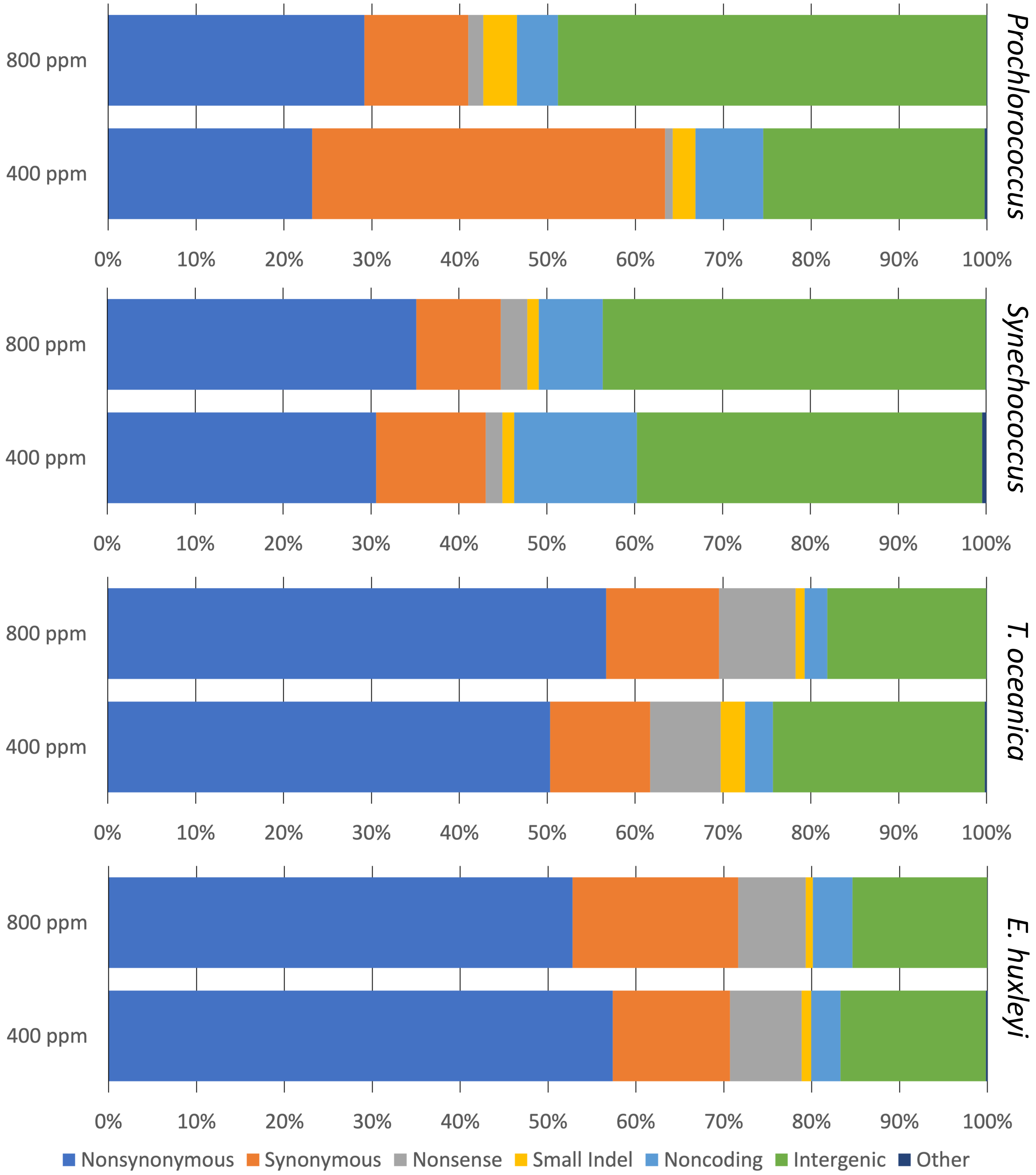


**Figure S11. Distributions of mutation types in *Alteromonas* genomes evolved under different $pCO_2$ regimes in partnership with the indicated phytoplankton strain.**

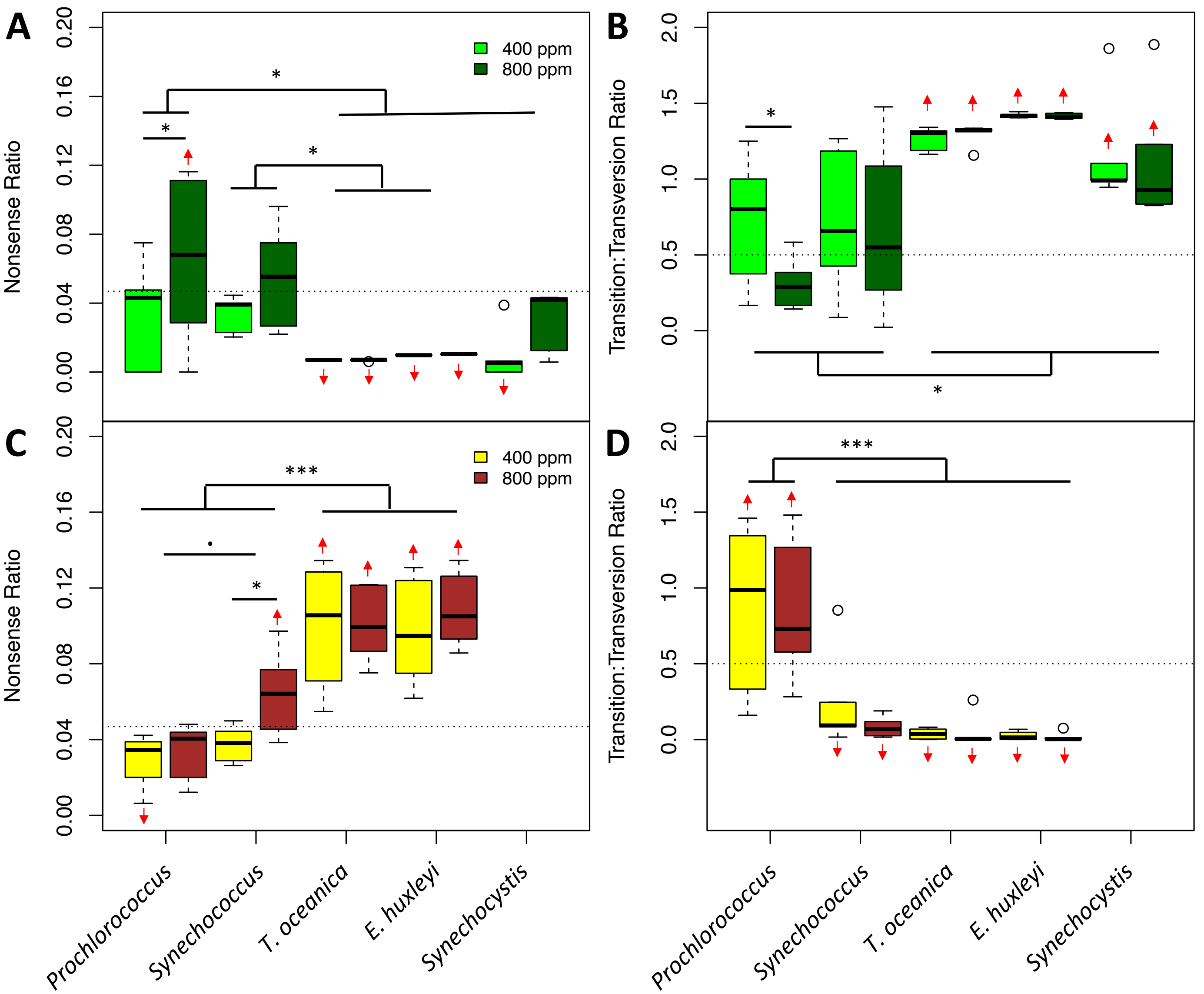


**Figure S12. Genomic evidence of adaptive evolution.** Plots in A and C show the proportions of codon changes resulting in stop codons (i.e., nonsense mutations) in coding sequences of phytoplankton species (A) or *Alteromonas* strains paired with the indicated phytoplankton species (C). Plots in B and D show the ratios of transitions to transversions. Dashed lines indicate the expected value under neutral evolution; red arrows indicate predicted means significantly higher or lower than this expected value (linear model, 95% confidence interval of the extended marginal mean). Asterisks indicate significant differences between $pCO_2$ treatments within a species or between species or groups of species: *, $p < 0.05$; ***, $p < 0.001$, ., $p < 0.1$.

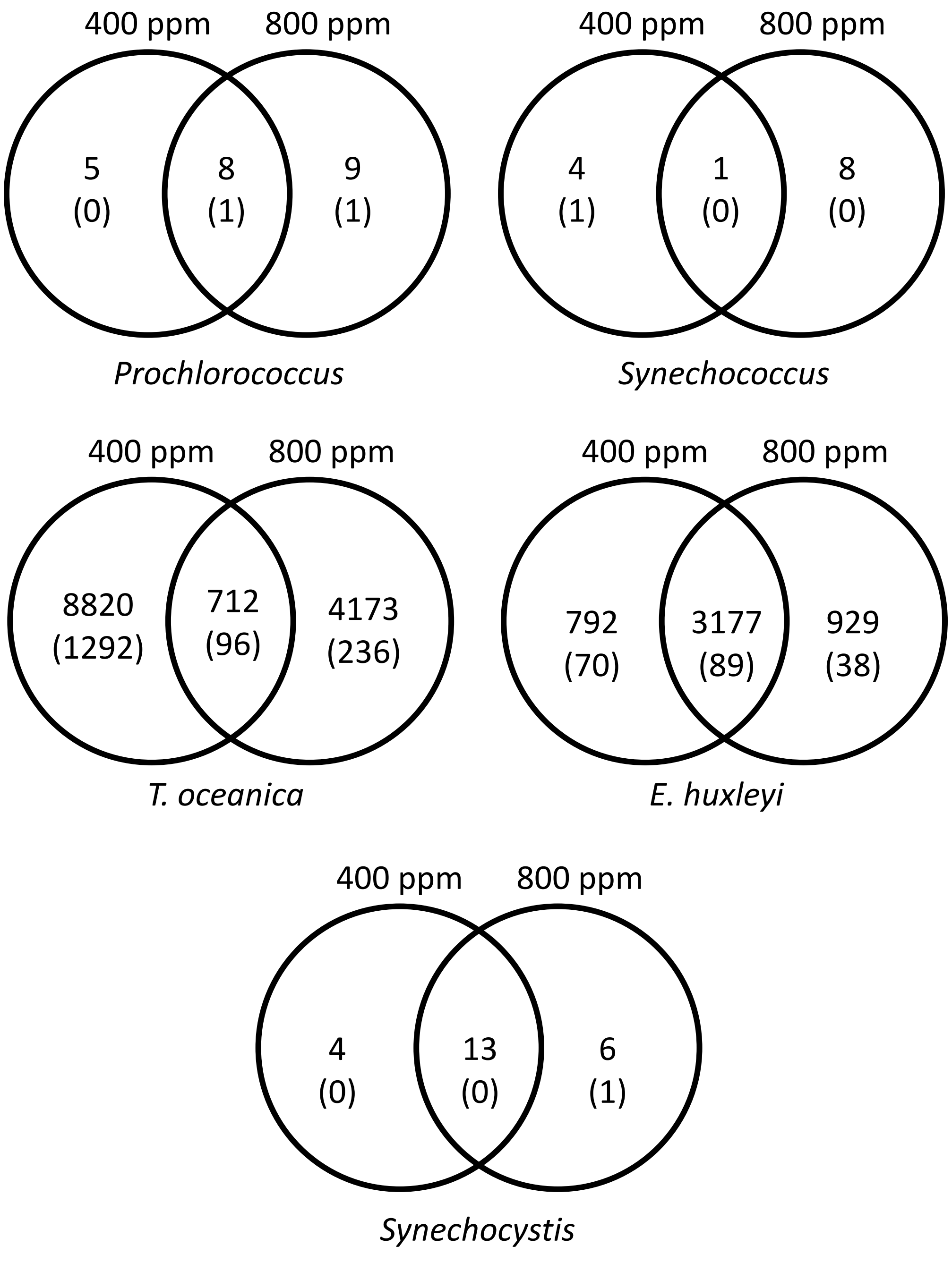


**Figure S13. Genes in phytoplankton genomes that were significantly more mutated, by $pCO_2$ treatment.** In order to be considered significantly multiply mutated, a gene had to have either i) more observed nonsynonymous, indel, or promoter mutations than in any of our bootstrapped dummy datasets (see Methods), and to not also have more synonymous mutations than in the dummy datasets, or ii) it had to have at least one observed nonsynonymous mutation in at least 50% of replicately evolved lineages. Values indicate the number of genes passing these criteria in only one of the $pCO_2$ treatments versus in both; values in parentheses indicate the number of thus identified genes that were also marked as statistically significantly differentially mutated between $pCO_2$ treatments in a linear model.

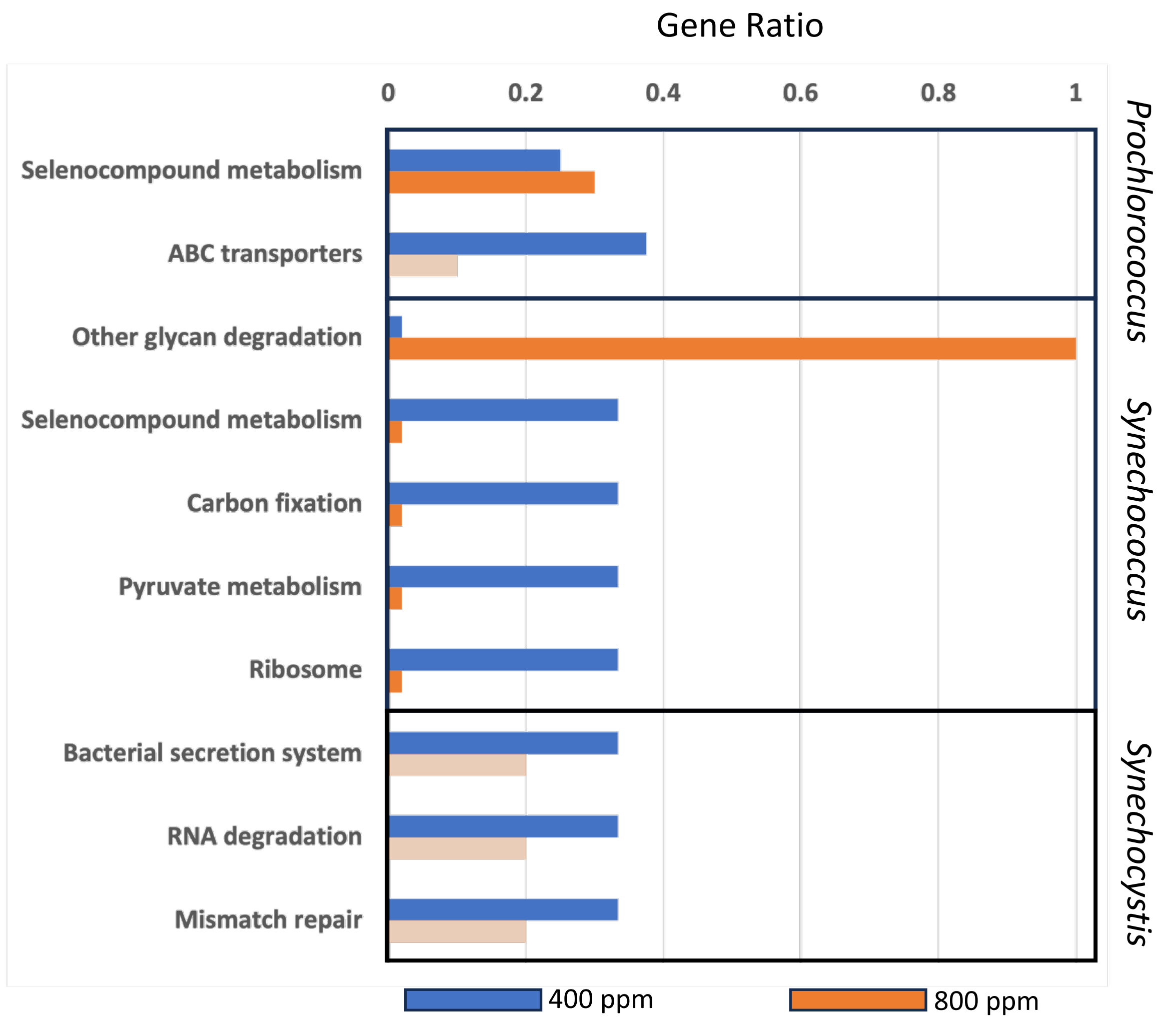


**Figure S14. Over-representation analysis of cyanobacterial mutations.** Genes passing our multiple mutation screening criteria were assigned to KEGG pathways as described in the Methods. Bold colors indicate that the pathway is statistically significantly overrepresented; paler colors indicate a lack of significance. Very small bars are placeholders only and correspond to absence of mutations observed in the indicated $pCO_2$ condition.

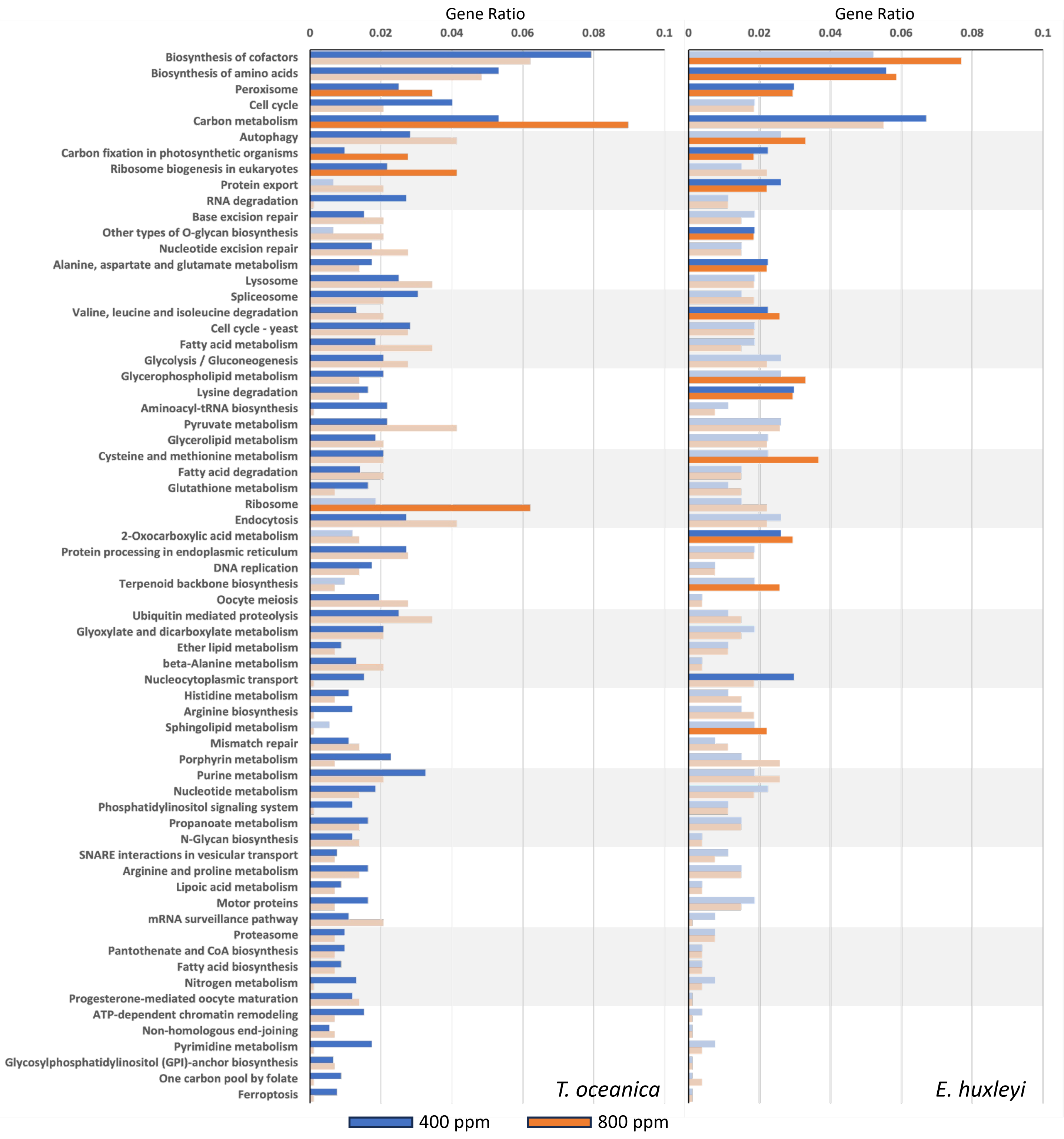


**Figure S15. Over-representation analysis of eukaryotic phytoplankton mutations.** Genes passing our multiple mutation screening criteria were assigned to KEGG pathways as described in the methods. Bold colors indicate that the pathway is statistically significantly overrepresented; paler colors indicate a lack of significance. Very small bars are placeholders only and correspond to absence of mutations observed in the indicated $pCO_2$ condition.

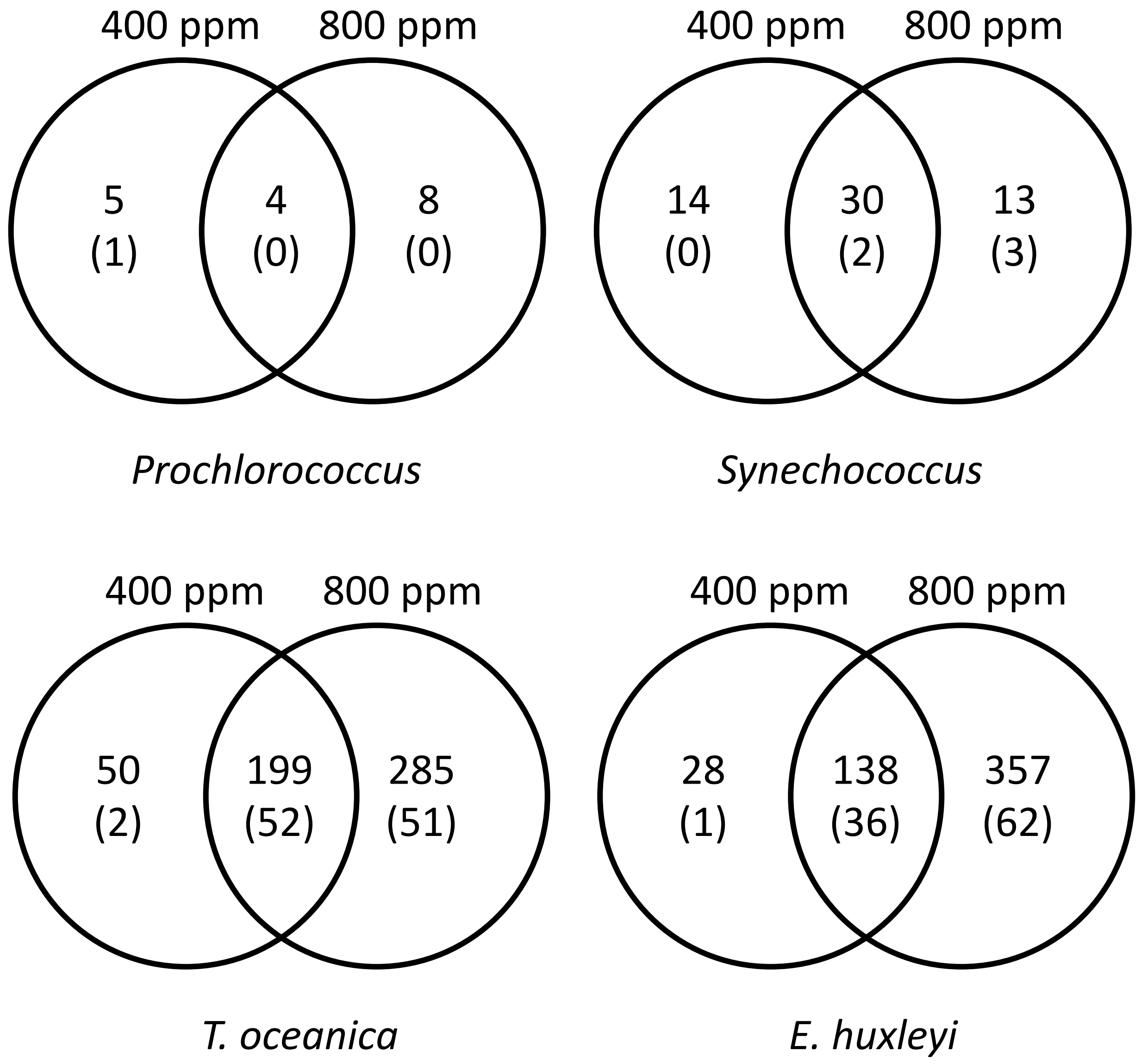


**Figure S16. Genes in *Alteromonas* genomes that were significantly more mutated, by $pCO_2$ treatment.** In order to be considered significantly multiply mutated, a gene had to have either i) more observed nonsynonymous, indel, or promoter mutations than in any of our bootstrapped dummy datasets (see Methods), and to not also have more synonymous mutations than in the dummy datasets, or ii) it had to have at least one observed nonsynonymous mutation in at least 50% of replicately evolved lineages. Values indicate the number of genes passing these criteria in only one of the $pCO_2$ treatments versus in both; values in parentheses indicate the number of thus identified genes that were also marked as statistically significantly differentially mutated between $pCO_2$ treatments in a linear model.

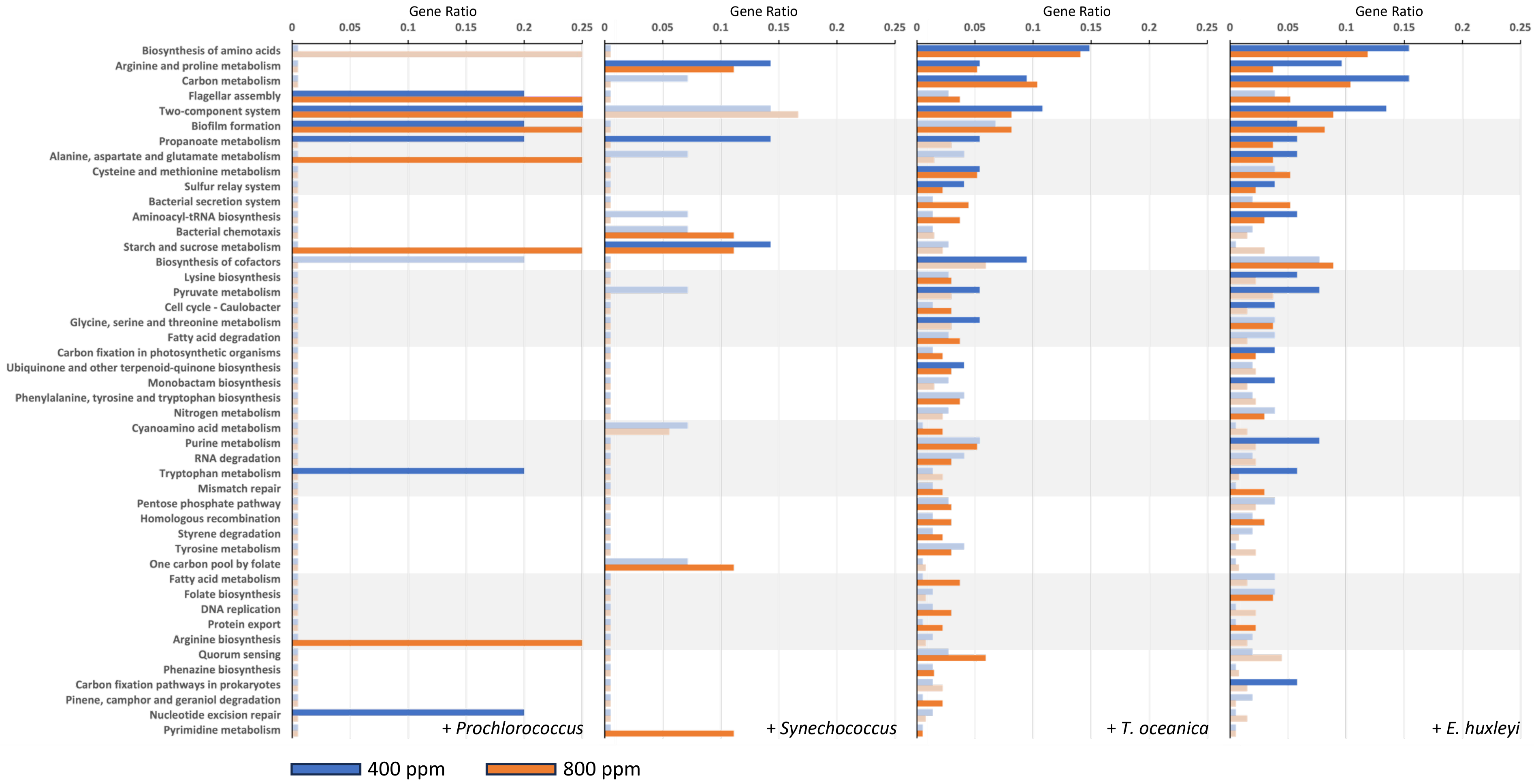


**Figure S17. Over-representation analysis of *Alteromonas* EZ55 mutations.** Genes passing our multiple mutation screening criteria were assigned to KEGG pathways as described in the Methods. Bold colors indicate that the pathway is statistically significantly overrepresented; paler colors indicate a lack of significance. Very small bars are placeholders only and correspond to absence of mutations observed in the indicated $pCO_2$ condition.

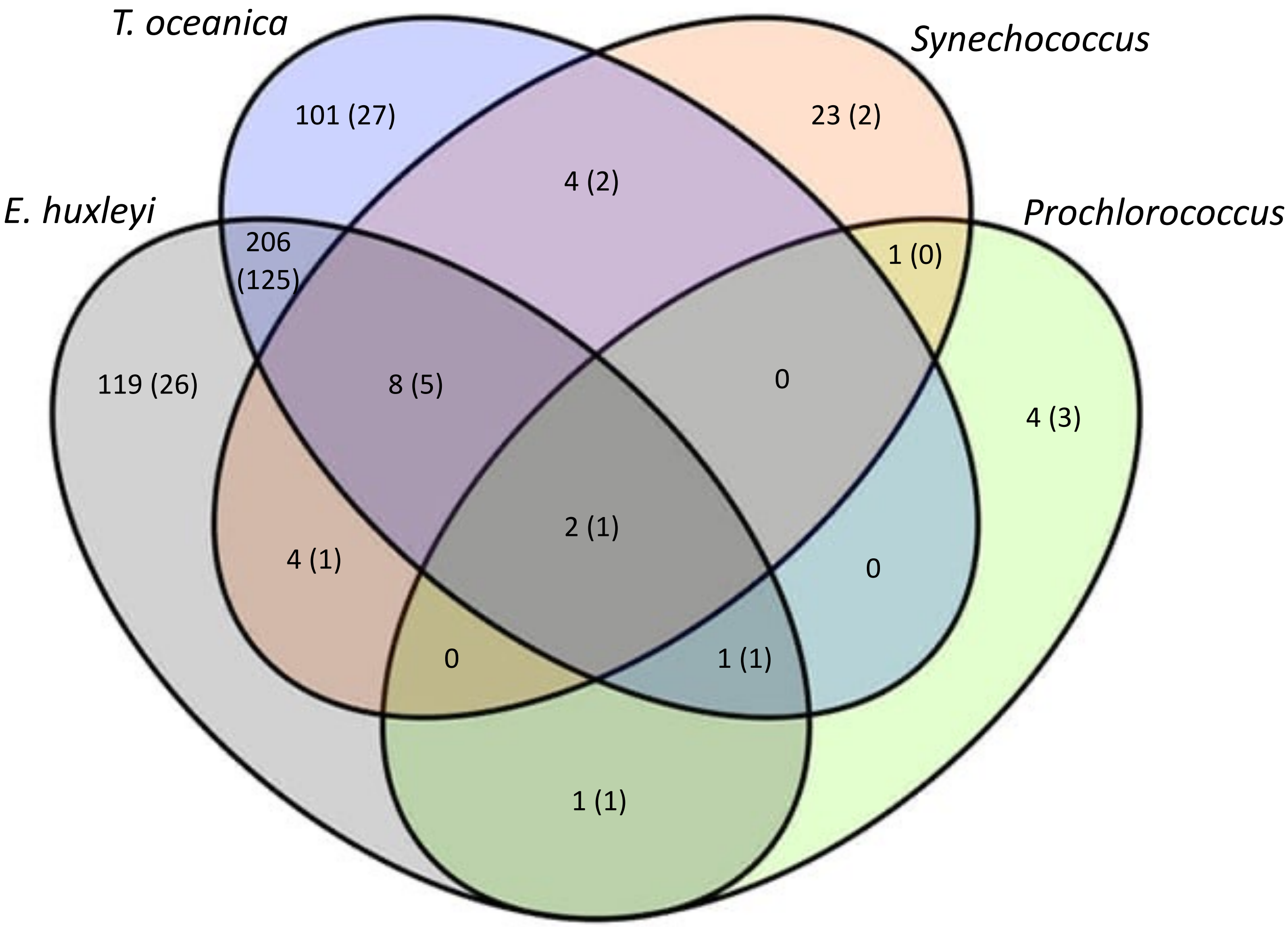


**Figure S18. Genes in *Alteromonas* genomes that were significantly more mutated, by phytoplankton partner.** In order to be considered significantly multiply mutated, a gene had to have either i) more observed nonsynonymous, indel, or promoter mutations than in any of our bootstrapped dummy datasets (see Methods), and to not also have more synonymous mutations than in the dummy datasets, or ii) it had to have at least one observed nonsynonymous mutation in at least 50% of replicately evolved lineages. Values indicate the number of genes passing these criteria in for the partners indicated by overlapping ovals; values in parentheses indicate the number of thus identified genes that were also marked as statistically significantly differentially mutated between at least one pair of partners in a linear model.

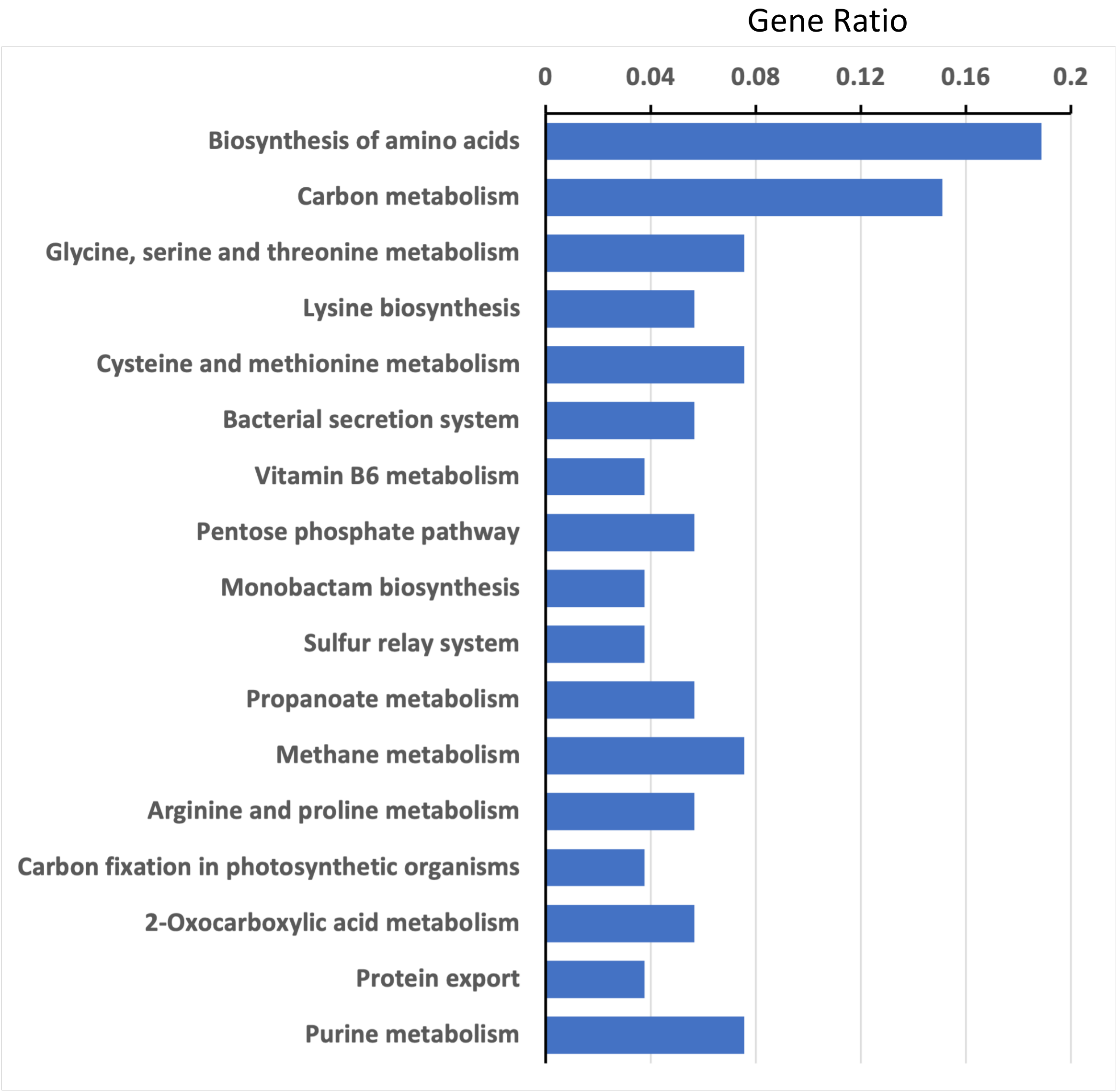


**Figure S19. Pathways analysis of *Alteromonas* EZ55 genes convergently mutated in co-culture with eukaryotic phytoplankton.** Genes passing our multiple mutation screening criteria only for *T. oceanica* CCMP1005 and *E. huxleyi* CCMP371 were assigned to KEGG pathways as described in the methods. Only pathways with adjusted p values $< 0.05$ for the statistical test of overrepresentation are depicted.

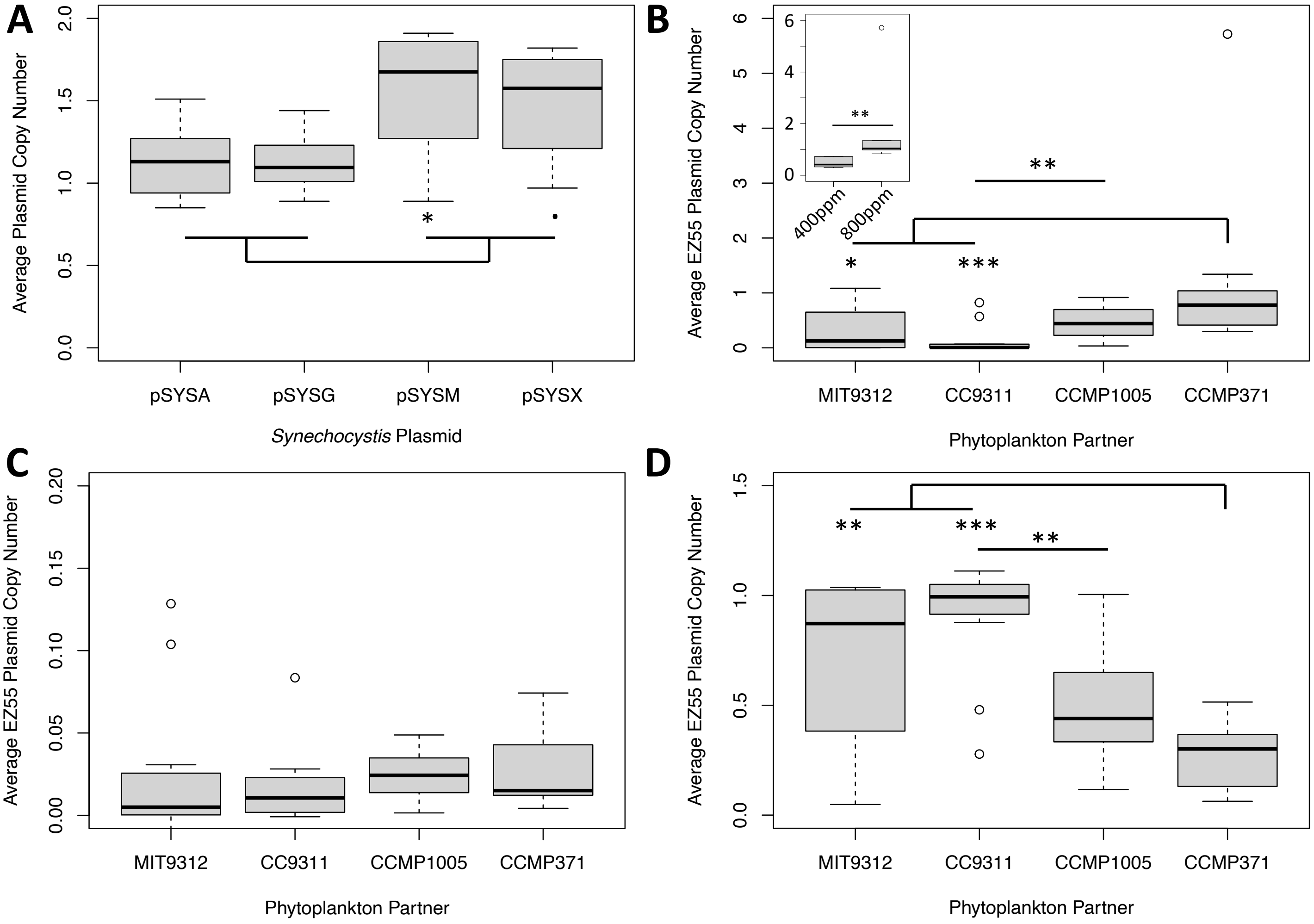


**Figure S20. Plasmid copy number.** Copy number was determined by dividing the average coverage of plasmid sequences by the average coverage of chromosomal sequences. A) Four plasmids from the *Synechocystis* PCC6803 genome. B) Copy number of the full, free *Alteromonas* EZ55 plasmid in co-cultures with *Prochlorococcus* MIT9312*, Synechococcus* CC9311, *T. oceanica* CCMP1005, and *E. huxleyi* CCMP371. The inset figure shows the copy number for EZ55 paired with *E. huxleyi* under different $pCO_2$ treatments, which were significantly different (Mann-Whitney test, $p = 0.008$). With the exception of strains partnered with *E. huxleyi*, all evolved EZ55 strains had mean copy numbers significantly less than 1, indicating the presence of plasmid-free segregants in the population. C) Copy number of the hypothetical reduced-size version of the EZ55 plasmid. D) Proportion of EZ55 cells with evidence of plasmid insertion in the chromosome. *, $p < 0.05$; **, $p < 0.01$; ***, $p < 0.001$; ., $p < 0.1$.

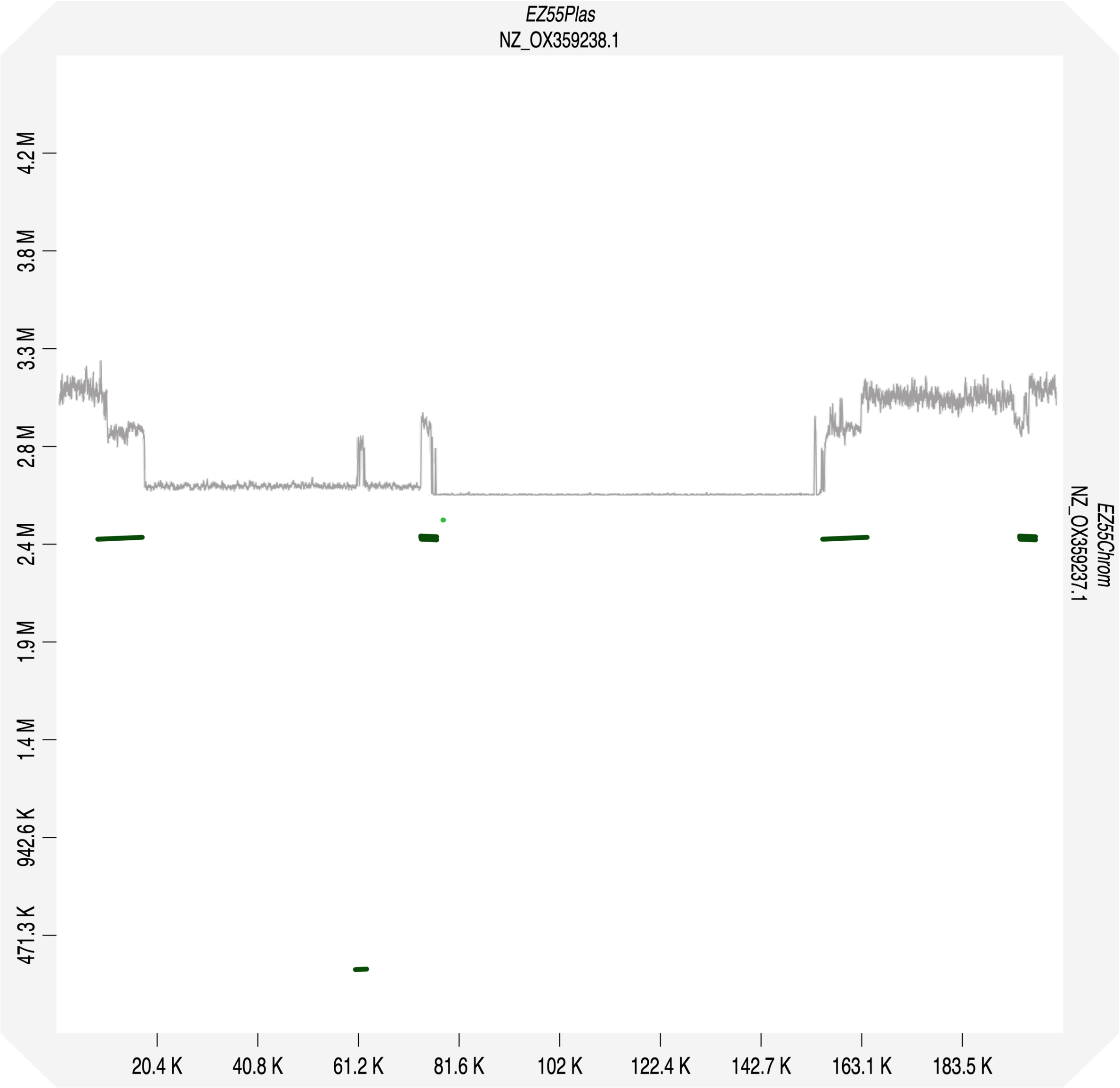


**Figure S21. Homology between EZ55 chromosome and plasmid DNA sequences.** Positions along the EZ55 main chromosome are represented on the y-axis, and positions along the plasmid are shown on the x-axis. Black dots indicate >95% homology between the paired sequences. Overlayed in light gray is a map of EZ55 plasmid coverage from LTPE428 (Fig. S6) scaled to match the x-axis, showing the erroneous mapping of chromosomal reads to deleted portions of the plasmid. The map also suggests potential targets for homologous recombination and insertion into the main chromosome that may explain the patterns of coverage and deletion observed.

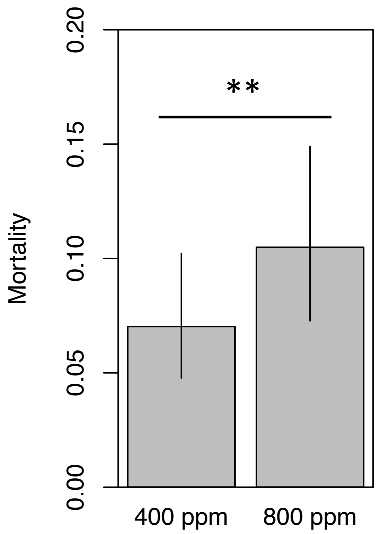


**Figure S22. Impact of $pCO_2$ on *Prochlorococcus* mortality.** Ancestral and evolved *Prochlorococcus* cultures were grown either axenically or in co-culture with either ancestral or evolved *Alteromonas* EZ55. The impact of the heterotrophic helper bacterium (or lack thereof) is shown in Figure 4A. Based on model predictions, cultures grown at 800 ppm experienced significantly greater mortality, but there was no interaction between the impact of $pCO_2$ and heterotrophic bacterium treatment on mortality. **, $p < 0.01$.

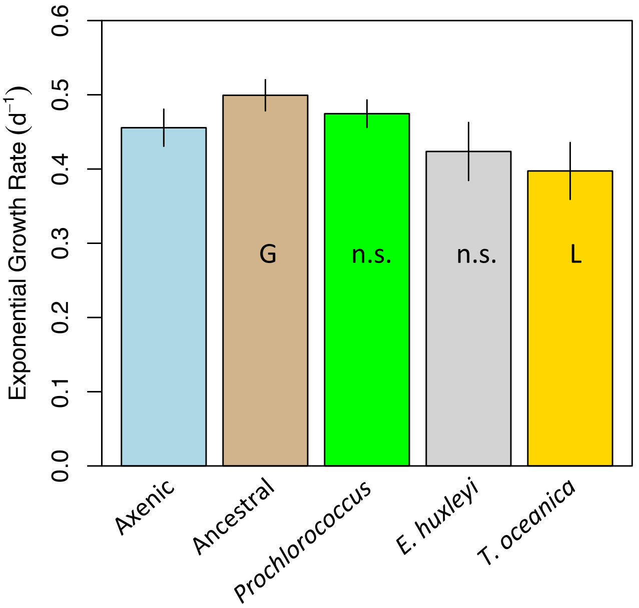


**Figure S23. Impact of adaptation to co-culture on ability of EZ55 to improve exponential growth rates of *Prochlorococcus* cultures**. Ancestral and evolved populations of *Prochlorococcus* MIT9312 were grown either axenically, in co-culture with ancestral *Alteromonas* EZ55, or with clones of EZ55 isolated from cultures of MIT9312, *E. huxleyi* CCMP371, or *T. oceanica* CCMP1005 after 500 generations of evolution at 400 ppm $pCO_2$. G and L indicate significantly ($p < 0.05$) greater or lower parameters based on the results of a Dunnett's test comparing each EZ55 treatment to the axenic control, whereas n.s. indicates the result of the comparison was nonsignificant ($p > 0.05$). There was not a significant difference between the impact of EZ55 on ancestral and evolved MIT9312, so the model estimated values are averaged between those treatments. Error bars represent the 95% confidence interval of the extended marginal means estimate of the indicated parameter.
